# Identification of CD4^+^ Sub-population of Resident Cardiac Fibroblasts Linked to Myocardial Fibrosis

**DOI:** 10.1101/2021.02.26.433023

**Authors:** Jamila H. Siamwala, Francesco S. Pagano, Patrycja M Dubielecka, Alexander Zhao, Sonja Chen, Haley Granston, Sakthivel Sadayappan, Sharon Rounds, Richard J. Gilbert

## Abstract

Infiltration with inflammatory T-cells and accumulation of cardiac myofibroblasts are hallmarks of cardiac fibrosis and maladaptive remodeling. The origin, identity, and functions of the resident cardiac cells involved in this process are, however, unclear. To determine the identity of cells contained in regions exhibiting fibrosis, mass cytometry profiling was performed using resident human ventricular cardiac fibroblasts and right ventricle autopsy tissues from individuals diagnosed with pulmonary hypertension and SUGEN/hypoxia rats. Results showed that a subpopulation of resident myocardial fibroblasts expresses increased levels of CD4^+^, a helper T-cell surface marker, in addition to mesenchymal markers in humans and rats. Characterization of the resident cardiac fibroblast subpopulation, both structurally and functionally, using transcriptome and secretome analysis of the secreted cytokines, chemokines, proteins, and metabolites, evidenced that IL-1β induces a phenotypic switch of human cardiac fibroblasts from mesenchymal to CD4^+^ lymphoidal lineage *in vitro*. RNA sequencing (RNA-seq) analysis of FACS-sorted CD4-expressing cardiac fibroblasts further revealed that the transcriptome of such IL-1β-induced CD4^+^ fibroblast population exhibited classical lymphoidal and stem cell-like signatures. Lastly, reversal of cell clustering, phosphorylation of MAPK p38 and NF-κB p65, and phenotypic switching was achieved with the administration of an IL-1R antagonist. In conclusion, we have identified a subpopulation of cardiac fibroblasts which exhibits structural and functional attributes of both mesenchymal and lymphoid cells which is induced by IL-1β-IL-1R-NFkB pathway for differentiation of cardiac fibroblast cells. These data suggest that cardiac fibroblast transdifferentiation during inflammation may form the basis for maladaptive remodeling during myocardial fibrosis.

## Introduction

Perivascular inflammation is commonly observed in the lungs of patients with pulmonary hypertension (PH) and is believed to play a critical role in determining their prognosis and pathology (1–5). Histological features of these lesions include perivascular inflammatory infiltrates involving T and B lymphocytes, macrophages, dendritic cells, and mast cells in the pulmonary arteries of both explanted human lungs and animal models of PH (2, 6). These lesions are, moreover, linked to elevated plasma concentrations of cytokines derived from infiltrating immune cells and serve as biomarkers of disease severity. However, the mechanism by which cytokines promote maladaptive cardiac and vascular remodeling in PH is unclear (7).

Among the cytokines reported to be elevated in PH, interleukin-1 (IL-1), a multipotent, inflammatory agonist of cardiac fibroblast cells, is known to enhance chemotaxis and adhesion molecule expression in response to myocardial injury (8). Interleukin-1β (IL-1β), a member of the IL-1 family, in particular, exacerbates tissue damage during acute and chronic injury by activating multiple signaling pathways, including phosphorylation of kinases, Ca^2+^ flux, cytoskeletal reorganization (9), and the release of inflammatory cytokines/chemokines, matrix metalloproteases (10) and nitric oxide (11). IL-1β binds to the Interleukin-1 receptor (IL-1R) to promote the release of secondary inflammatory cytokines, such as IL-6, IL-23, Tumor Necrosis Factor-Alpha (TNFα), and Granulocyte Colony-Stimulating Factor (G-CSF) (12). IL-1 may also drive the differentiation of T-cells into Th17 cells and effector cells to produce pathogenic cytokines, such as IFNγ, IL-17, and GM-CSF (13). Blocking the cytokine receptor IL-1R with an IL-1R antagonist reduces PH in a rat monocrotaline model (14). Similarly, the monoclonal antibody to IL-1β, canakinumab, decreases inflammation in right ventricular failure and PH in clinical Phase IB/II pilot studies, suggesting the critical role of IL-1β in PH-linked inflammation (15). While it is known that IL-1β is expressed by activated disease-derived fibroblasts (16), it is uncertain how IL-1β modulates cardiac fibroblast function or contributes to the inflammatory or fibrotic response in maladaptive remodeling typical in PH.

Fibroblasts are mesenchymal in origin and found in almost all cell and tissues. In myocardium, the cardiac fibroblasts exhibit heterogeneity in regard to their identity, origin, properties, signaling mechanisms, and response to environmental stimuli (17). Activated pulmonary myofibroblasts, for example, are posited to be the drivers of perivascular inflammation and temporarily assume an epigenetically altered pro-inflammatory phenotype, secreting cytokines, chemokines, and recruiting macrophages in calf, rat, and human models of pulmonary arterial hypertension (18). Activated resident cardiac fibroblasts can differentiate into myofibroblasts expressing smooth muscle cell actin (αSMA) and secreting collagen, fibronectin, and other extracellular matrix molecules (20, 21). Recent studies show that tissue resident cardiac myofibroblasts may also differentiate into endothelial cells as a response to local myocardial injury (1) and convert from a lipogenic cell to a myogenic cell during lung fibrosis (2). Lineage tracing experiments in a TCF-21^+^ resident cardiac fibroblast mice model further demonstrate a capacity for cardiac fibroblast cells to differentiate into cells expressing bone and cartilage markers suggesting plasticity and phenotype switching (3). They can also be reprogrammed into cardiomyocyte-like cells using transcription factors and miRNAs (22, 23). However, the modulators of resident cardiac fibroblasts plasticity and role of proinflammatory cytokine signals such as IL-1β in phenotype switching role and myocardial inflammation are poorly understood.

CD4^+^ T-cells infiltration levels are linked to overall survival rates in patients in other diseases and used as immunotherapy approach(4). Accumulating evidence suggests that CD4^+^ T cells participate in the repair of the myocardium after infarction, similar to the cardiac myofibroblast (24), by modulating collagen deposition, degradation, and the formation of fibrous tissues (25). In contrast to their reparative role, CD4^+^ T cells may also acquire a pro-inflammatory phenotype, thereby contributing to adverse cardiac remodeling in mice with chronic ischemic heart failure (26). Specifically, CD4^+^ T cells are involved in the transition from compensated cardiac hypertrophy to maladaptive cardiac remodeling and heart failure during chronic pressure overload (27). Blockade of T-cell co-stimulation in a transverse aortic constriction (TAC) mouse model of pressure overload reduces its severity and delays heart failure, suggesting a crucial role of CD4^+^ T-cells in the progression of this disease (28).

With the advent of state of art mass cytometry technique and viSNE and UMAP algorithms and improvement in isolation of primary human cardiac fibroblasts, it is now possible to scrutinize with greater depth the residing population and profile the cells to determine their phenotype and role in fibrosis. Using these tools, we present a discovery of a bridge population of cells bearing properties of both mesenchymal cells and lymphoidal cells referred to as fibrolymphocyte which we think unifies the isolated reports on CD4^+^ cells and cardiac myofibroblasts in fibrotic regions.

## Results

### Single-cell multidimensional mass cytometry identifies distinct cardiac fibroblast sub-populations expressing mesenchymal and lymphoid markers

PH patients have high levels of plasma IL-1β that correlate with the severity of PH (*15, 43*), and reduction of IL-1β reduces inflammation and improves right heart function (15). Our studies and those of others show that IL-1β is increased in the right ventricle of rats exhibiting pulmonary hypertension in concentrations that are known to activate cardiac fibroblasts isolated from the rat right ventricle with varying severity and outcomes (*35–38*). We also observed heterogeneic responses of rat cardiac fibroblasts to modulators and inhibitors raising the possibility of plasticity and phenotype switching in the presence of cytokines as typically observed in induced pluripotent stem cell derived from adult fibroblasts. To identify the cardiac fibroblast transitory states and functional diversity of human primary ventricular cardiac fibroblasts (hVCF), we employed advanced, highly specific mass cytometry (Figure 1A) technique on cultured primary human cardiac fibroblasts expressing the prototypical fibroblast markers αSMA, vimentin, collagen-1, FSP-1, PDGFRβ, and periostin, but negative for the endothelial cell marker VE-cadherin (Figure S1) (*39–42*) and low frequency of PDGFRβ-positive cells were negative for FSP-1, and some FSP-1-positive cells were negative for PDGFRβ (Figure S1). Next, we determined the immune identity of the resident hVCFs by barcoding cardiac fibroblasts with previously validated heavy metal immune cell lineage- specific antibodies and by separating cell masses by time of flight (Figure 1, Panel A). Cisplantin, a cell viability stain, was included in the panel of antibodies selected for sub-categorization of primary human cardiac fibroblast cells. Signals integrated per cell and both marker intensity and distribution were visualized using dimensionality reduction and clustering algorithms. First, the cell populations were cleaned by removing the beads and gated for live, nucleated single-cell cell populations (Figure S2). Dimensionality reduction performed using viSNE analysis and T-distributed Stochastic Neighbor Embedding (tSNE) algorithms showed specific hVCF clusters expressing αSMA^+^ and Vimentin^+^ and similar levels of CD4^+^, a hallmark helper-T-cell marker among other immune cell markers constituting 2.8% ± 0.15% of the total analyzed cell populations (Figure 1, Panel B and Figure S3). Further, this population expressed other lymphoid cell markers, such as CCR6, and CD183 but not CD3 due to antibody degradation (Figure 1, Panel C), confirming the lymphoid lineage of the newly identified subpopulation of cardiac fibroblast cells. Consistent to the viSNE algorithm, another higher resolution clustering algorithm using uniform manifold approximation and projection (UMAP) non-linear dimensionality reduction using other parameters also showed distinct cell population expressing mesenchymal and other known immune cell specific markers thereby validating the previous finding (Figure S4).

**Figure 1:**
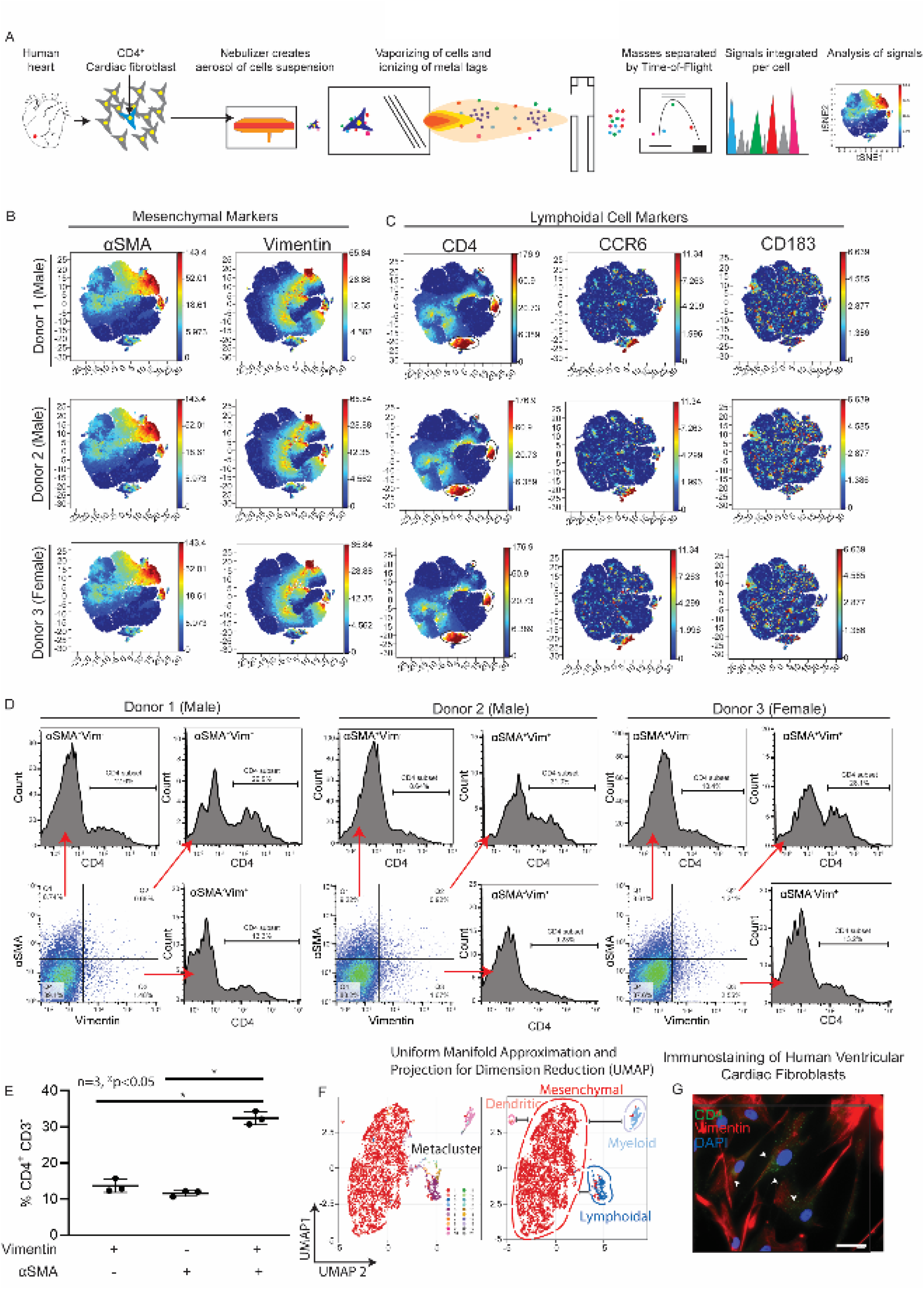
Identification of a subpopulation of resident hVCFs expressing mesenchymal and lymphoid cell markers with multidimensional single-cell mass cytometry. Immunophenotyping of hVCFs derived from human subjects (2 males (ID# 62122, ID# 1281202) and 1 female (ID#534282) was performed by staining 3 x 10^6^ cells with heavy metal-tagged antibodies (Table S3), followed by mass cytometry to identify single-cell expression. (A) Schematic summary of mass cytometry analysis. hVCFs were immunostained with epitope-specific antibodies conjugated to transition element isotope reporters of different masses. The cells were nebulized into single-cell droplets, followed by acquisition of an elemental mass spectrum. Conventional flow cytometry methods were used to analyze integrated elemental reporter signals for each cell. (B) viSNE graphs of manually gated live, nucleated, hVCF cell clusters expressing varying marker intensities and distribution for a representative donor. Resident hVCFs were marked by Vimentin expression, and activated cardiac fibroblasts were marked by αSMA expression. (C) The lymphoid lineage protein-expressing hVCFs were marked by the cluster of differentiation (CD) CD4, CCR6, and CD183 on hVCF. Protein expression levels are demonstrated on the secondary y-axis scale with blue showing no expression, green, the least expression, and red, the highest expression. Each dot represents the expression profile of a single cell. (D) Representative flow cytometry histograms showing CD4^+^ subset of αSMA^+^Vim^+^ hVCF’s for all the human donor cells detected using mass cytometry. (E) Quantification of CD4^+^ CD3^-^ cells gated from Vimentin^+^αSMA^-^, Vimentin^-^αSMA^+^ and Vimentin^+^ αSMA^+^ populations is represented as a scatter plot of Mean ± SD (n=3 biological replicates, 2 males and 1 female; \**P* <0.05, as determined using one-way ANOVA and Tukey’s post-hoc multiple comparison. (F) Meta-clustering of the populations based on the distances of neighbors (UMAP algorithm) to provide a global picture of cell populations. The colors stratify the similar and dissimilar cell populations showing heterogeneity. The dotted lines represent the 4 major populations (Lymphoid-, Myeloid-, Mesenchymal- and HLA-positive cells) that were identified in the primary human cardiac fibroblast. (G) Representative hVCF cells immunostained CD4, Vimentin and DAPI antibodies showing coexpression of Vimentin and CD4.

To determine the frequency of activated cardiac fibroblast cells marked by αSMA^+^ expressing CD4^+^ and resident cardiac fibroblast cells marked by Vimentin expressing CD4^+^ hVCF, cells expressing Vimentin^+^αSMA^+^ (Figure S5), Vimentin^+^αSMA^-^ (Figure S6) or Vimentin^-^αSMA^+^ were gated (Figure S7). Of the total population of hVCF cells expressing CD4^+^, 29.6% of the total cells counted were Vimentin^+^αSMA^+^CD4^+^, 11.91% ± 2.32% cells of the total cells were Vimentin^+^αSMA^-^CD4^+^ resident cardiac fibroblasts, and 10.53% ± 2% of the total cells were αSMA^+^Vimentin^-^CD4^+^ activated cardiac fibroblasts (Figure 1, Panel E). The total percentages of CD4^+^ subpopulations expressing either αSMA or Vimentin or both are summarized in Figure S8. However, cardiac fibroblast cells expressing the triple Vimentin^+^αSMA^+^CD4^+^ marker were about 3% of the total hVCF populations analyzed. Primarily, viSNE and UMAP algorithm distinguished four primary human ventricular cardiac fibroblast subsets, consisting of populations expressing markers of lymphoid cells, myeloid cells, mesenchymal cells, and dendritic cells (Figure 1D, all donors).

Moreover, resident cardiac fibroblasts expressing CD4 marked by Vimentin^+^ cells were confirmed by immunostaining (Figure 1, Panel G). To exclude the possibility of contaminating T-cells in primary cultures, the cells were cultured in fibroblast growth media without T-cell activators for at least two to three passages. Besides size-based exclusion, adherent cardiac fibroblasts were washed several times to ensure that any loosely adhered immune cells were removed. In the aggregate, our data support the existence of a previously unidentified resident cardiac fibroblast subpopulation that co-expresses mesenchymal vimentin^+^αSMA^+^ and lymphoid CD4^+^ markers in humans.

### Distribution and expression of spindle-shaped αSMA^+^ CD4^+^-coexpressing cells in the fibrotic right ventricle (RV) of patients with Pulmonary Hypertension (PH)

Following the identification of a CD4^+^ human ventricular cardiac fibroblast (hVCF) sub- population *in vitro* using mass cytometry, the physiological relevance of these cells to the genesis of cardiac fibrosis and PH was determined in autopsy specimens of patients with right ventricular fibrosis and pulmonary hypertension. Autopsy diagnoses of human RV tissue donors in the groups PH and no PH are provided in Table S4. Morphological characterization of RV tissue using Hematoxylin and Eosin (H&E), Sirius Red and CD4 or αSMA/CD4 immunostaining showed RV hypertrophy and fibrosis in both the PH and no PH groups. Variable amounts of cardiac myocyte hypertrophy and age-associated fibrosis in interstitial, perivascular, and subendocardial locations in the H&E-stained and Sirius red sections were noted in all the cases (Figure 2, Panel A). Interstitial fibrosis was observed within the posterior papillary muscle of the right ventricle. Endocardial surfaces of right ventricle showed hypertrophic changes with big “boxcar” nuclei. Donor A demonstrated hypertrophic cardiac myocytes with “boxcar” nuclei and mild interstitial and perivascular fibrosis, highlighted by Sirius Red. Donor B demonstrated hypertrophic cardiac myocytes with marked sub-endocardial and interstitial fibrosis with entrapped cardiac myocytes in the areas of fibrosis. Cardiac myocytes surrounding the area of dense fibrosis was notably hypertrophied. Sirius Red staining highlighted extensive subendocardial fibrosis and interstitial fibrosis. Donor C demonstrated hypertrophic cardiac myocytes with subendocardial and patchy interstitial fibrosis. Sirius Red highlighted the bands of interstitial and subendocardial fibrosis and appeared focally pericellular in areas. Donor D demonstrated fewer hypertrophic cardiac myocytes with scattered foci of interstitial fibrosis. Sirius Red staining of the interstitial fibrosis highlighted larger areas of fibrosis, and pericellular fibrosis surrounding individual cardiac myocytes. Finally, Donor E demonstrated predominately perivascular fibrosis, highlighted by the Sirius Red stain (Figure 2).

**Figure 2:**
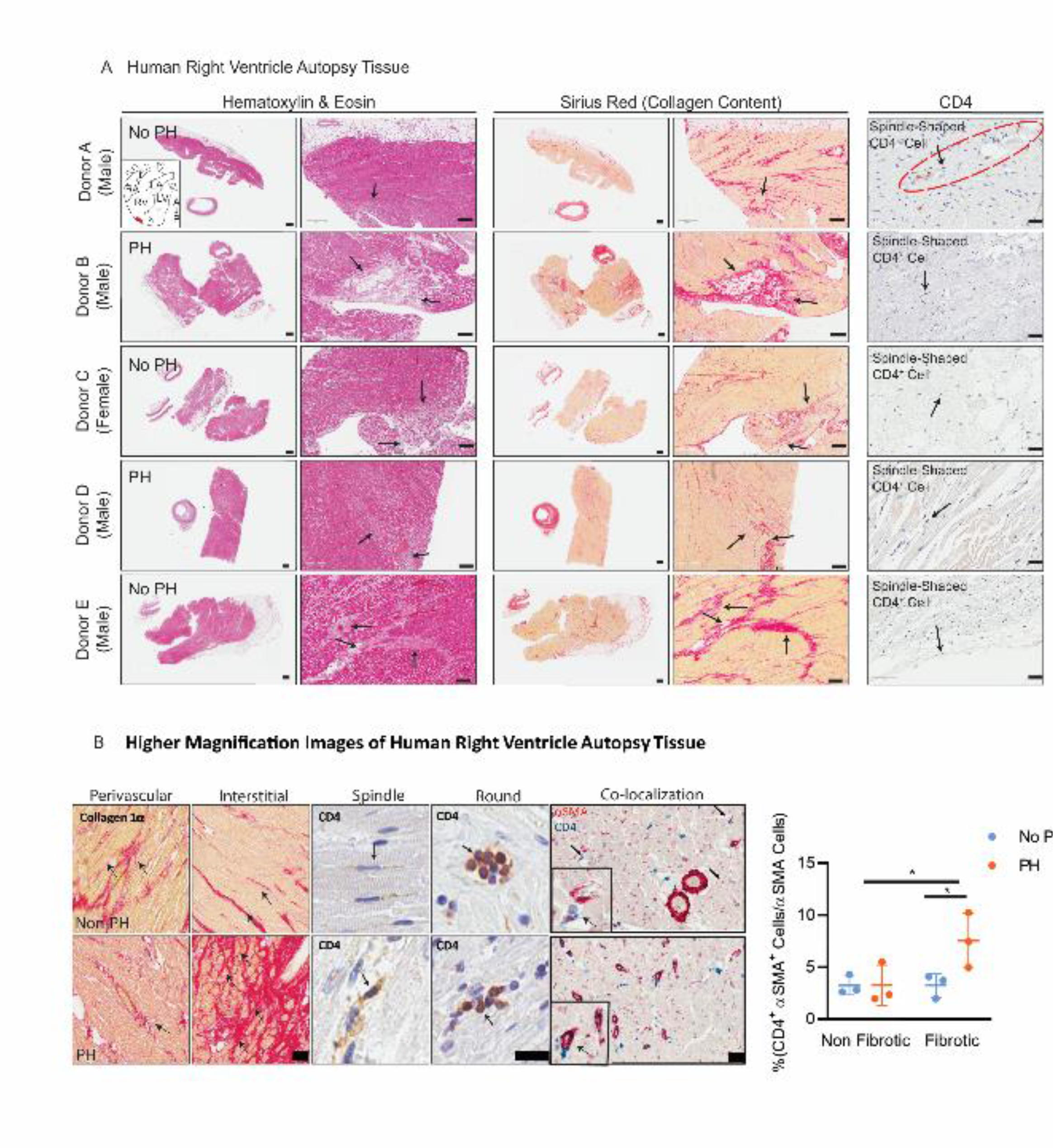
Distribution and expression of CD4 in the right ventricle in cardiac fibrosis in humans and in rat model of SUGEN/Hypoxia PH. (A) Human right ventricular tissues from 5 autopsied donors diagnosed with other conditions (Donor A, C and E) or PH (Donor B, D and F) were stained with standard hematoxylin & eosin (H&E) and Sirius Red. Donors A-E were immunostained for CD4 and counterstained with methyl green to identify nuclei. Donor F was used as a negative control for CD4 immunostaining. The black arrows point to fibrotic areas in H&E-, Sirius Red- and CD4- stained sections. CD4^+^ expressing spindle-shaped cells in the RV of donor tissue was determined; arrows point to CD4^+^ spindle-shaped cells. Quantification of dual-stained αSMA (red) and CD4 (blue) in the fibrotic and nonfibrotic regions of human RV tissue sections performed using immunohistochemistry. (B) Magnified images of Sirus red staining, CD4 staining (brown) and CD4/αSMA dual-staining in human RV. Arrows indicate the spindle-shaped cells expressing CD4. Quantification of CD4^+^αSMA^+^ cells in non-fibrotic and fibrotic regions in the human RV determined by a pathologist blinded to the groups. Scale bars, 75 µM. \**P*<0.05, one-way ANOVA and Tukey’s multiple comparison test. Quantification of CD4^+^/αSMA^+^ manually by a double-blinded observer from at least 15-20 sections/slide for n=6 samples (\**P*<0.05, one-way ANOVA and Tukey’s multiple comparison test; Nonfibrotic *vs*. Fibrotic and no PH *vs.* PH). Quantification of collagen I/III content in the human RV presented as a percentage of Sirius Red-positive region/total region is shown as scatter plots of Mean ± SD (n=3 donors, P=0.0897, one-tailed unpaired *t* test and Mann Whitney Analysis of Ranks; no PH *vs.* PH). Scale bars are 10 µM.

Cells with a spindle-shaped morphology and membranous CD4^+^ staining were present in the perivascular connective tissue in cases with minimal interstitial fibrosis and in the dense fibrous tissue seen in some cases (Figure 2, Panel A). Moreover, significant increases were found in spindle-shaped cardiac fibroblasts expressing both αSMA and CD4 in the fibrotic regions compared to the non-fibrotic regions of RV tissue in humans with a clinical diagnosis of right heart hypertrophy and dilation, compared to RVs from those without an autopsy diagnosis of PH (Figure 2, Panel B). Cells with a round shape and membranous CD4^+^ staining typical of traditional T-cells were predominantly identified inside capillaries (Figure 2, Panel B). The RV collagen content tended to increase in donors with a diagnosis of PH (Figure 2, Panel B). These results demonstrated the diversity within the population of cardiac fibroblast cells and suggested the existence of a CD4^+^ subpopulation of cardiac fibroblasts in the fibrotic regions of the human right ventricular diseased tissue.

### Distribution and expression of spindle-shaped αSMA^+^ CD4^+^-coexpressing cells in the fibrotic right ventricle in rats treated with SUGEN/hypoxia

After determining the distribution and expression of αSMA^+^CD4^+^ cells in the fibrotic regions of human right ventricular tissue, we sought to confirm the universality of the unique cardiac fibroblast sub-population in RV and LV using a rat SUGEN/hypoxia (SuHx) model of PH. Fischer rats, genetically prone to develop right heart failure and RV remodeling and PH were used as a model of cardiac fibrosis (29). Fischer rats were treated with a single intraperitoneal injection of the VEGF inhibitor SUGEN (25mg/kg) and then subjected to 3 weeks of hypoxia (Hx), followed by 5 weeks of normoxia (Nx). Control animals were injected with vehicle and housed in room air or Nx for 8 weeks. Collagen I/III expression increased significantly in both perivascular and interstitial regions in RV and LV, as determined by Sirius red staining and Masson trichrome staining (Figure 3, Panel B and Figure S10, Panel A). The SUGEN/Hypoxia rats had lower body weights compared to control rats (Figure S10, Panel B). Further, changes in cardiac fibroblast number and localization in response to SuHx treatment was determined using the fibroblast marker FSP-1 (Figure S10, Panel C). Tricuspid annular plane systolic excursion (TAPSE) was significantly decreased in SUGEN/Hypoxia-treated rats, providing echocardiographic evidence of PH (Figure 3, Panel A), however, the Fulton index measured from the RV and LV weights using the formula (RV/LV+S) was not significantly increased (Figure 3, Panel A). None of the other echocardiographic parameters measured, such as cardiac output, ejection fraction, and mean pulmonary arterial pressure, was significantly different from controls. Representative M-mode TAPSE tracings and pulmonary outflow traces are shown in Figure S11. Next, the number of cardiac fibroblasts co-expressing αSMA^+^CD4^+^ cells were quantified in the fibrotic regions of the RV of Nx and SuHx rats using confocal microscopy (Figure 3, Panel C).

**Figure 3:**
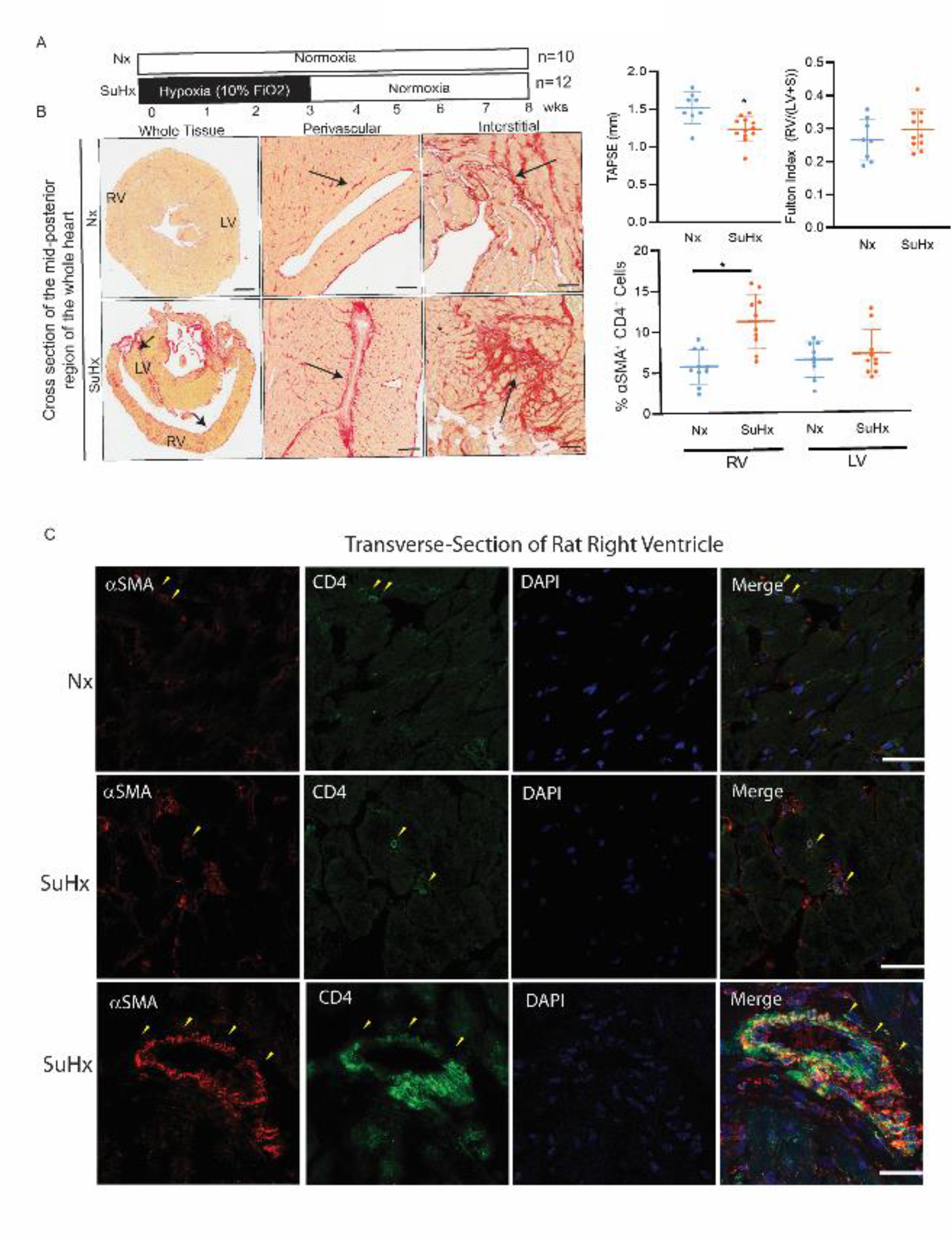
Histological analysis of the right ventricle in the setting of the SUGEN/hypoxia rat model. (A) Schematic representation of SUGEN/hypoxia for simulating PH in rats. (A) Male Fischer rats were given a single bolus of SUGEN (20mg/kg) and exposed to hypoxia (10% FiO_2_) for 3 weeks, followed by exposure to normoxia (Nx) for an additional 5 weeks (SuHx). In parallel, control animals were exposed to Nx (room air) for 8 weeks. The Nx and SuHx groups were comprised by n=10 and n=12 rats, respectively. Quantification of TAPSE trace measured using M-mode echocardiography is presented for Nx and SuHx rats. Fulton index determined from the weight of the RV and LV+ septum is indicated. Values are mean ± SD (n=10 rats) \**P*<0.05, unpaired t-test or one-way ANOVA and Tukey’s multiple comparison test (RV *vs.* LV, Nx *vs.* SuHx). Scale bars are 10 µM. (B) Representative images of collagen content in the of the perivascular or interstitial RV and LV indicated by the Sirius Red-positive regions in the transverse region of Nx and SuHx rats (10 µm thickness), whole heart section cut transversely at the mid-ventral region and perivascular and interstitial regions of both Nx and SuHx animals. Arrows suggest collagen I rich regions. (C) High magnification of the perivascular region of Nx and SuHx RV tissue expressing αSMA^+^ (red), CD4^+^ (green), and DAPI (blue)

Based on the markers, three cell populations: T-cells (αSMA^-^ CD4^+^), cardiac fibroblasts (αSMA^+^CD4^-^), and cardiac fibroblasts expressing CD4 (αSMA^+^CD4^+^) were identified in fibrotic regions of rat RV (Figure 3, Panel C) similar to the human tissue. We noted two morphologies of conventional CD4^+^cells, round αSMA^+^CD4^+^ and spindle-shaped αSMA^+^CD4^+^, in the RV regions, that were significantly increased in SuHx compared to Nx rats (Figure 3, Panel C). The number of αSMA^+^CD4^+^ cells tended to be higher in the perivascular fibrotic regions of the RV of SuHx rats compared to control (Figure 3, Panel C).

### Proliferation and differentiation of primary human cardiac fibroblasts in response to the pro-inflammatory cytokine, IL-1β

. After establishing the identity and localization of the unique subpopulation of hVCF cells in healthy cells and diseased human and rat tissues, we determined the regulation of this population by recombinant IL-1β based on the findings that IL-1β levels are elevated in the stressed human myocardium and animal models (*44–47*). We postulated that the proinflammatory cytokine IL-1β contributes to the induction of CD4 expression in resident cardiac fibroblasts. Heat-map analysis from mass cytometry data provided an overview of type and intensities of expression of different immune markers (Figure S12, Panel A). In addition, heatmaps suggested that CD4 expression increased with IL- 1β treatment specifically for male donors (Figure S12, Panel B). The non-redundancy scores suggested that CD4 was expressed at levels similar to those of vimentin and αSMA, markers of cardiac fibroblasts (Figure S12, Panel C). The percentages of HLA-DR^+^ dendritic cell, CD4^+^ lymphocyte, and CD68^+^ monocyte populations shifted with IL-1β treatment (Figure S12, Panel D). αSMA^-^ and CD4^-^expressing cardiac fibroblast populations increased with IL-1β treatment, as seen in the normalized density distribution plots (Figure S12, Panel E).

To determine whether IL-1β mediated the fibrotic responses by stimulating cardiac fibroblast proliferation, hVCFs were treated with vehicle or recombinant IL-1β for 24 h, followed by BrdU and MTT assays and Ki67 staining. IL-1β treatment resulted in a statistically significant increase in cardiac fibroblast proliferation in a dose-dependent manner (Figure 4, Panel A). No difference in the proliferation rate of cardiac fibroblasts in response to IL-1β was observed with time (Figure 4, Panel B). IL-1β similarly affected the proliferation rates of primary hVCF cells isolated from males and females (Figure 4, Panel C). IL-1β treatment tended to stimulate the deposition of Collagen Iα, as determined from total collagen in the lysates (Figure 4, Panel D) and confocal microscopy (Figure S13); however, these effects were not statistically different. To determine whether IL-1β differentiated cardiac fibroblasts and induced phenotypic changes, human hVCFs were treated with IL-1β for longer periods (96 hours), and this resulted in the induction of significant morphological changes in hVCF cells by day 4, including increased detachment and rounding of cells (Figure 4, Panel E). Round cells with non-homogeneous DAPI-stained nuclei appeared with 96 h of IL-1β treatment (Figure 4, Panel E). The round cells, but not the surrounding cells, were positive for both αSMA and CD4^+^ T cells (Figure 4, Panel E). The quantification of round cells showed a significant increase in these cells for all male and female donors with IL-1β treatment for 96 h (Figure 4, Panel E). The round cells were viable and not apoptotic, as determined using trypan blue staining. We next characterized the subcellular phenotypic switching of hVCFs with IL-1β after 96 h using transmission electron microscopy. Transmission electron microscopy showed more prominent endoplasmic reticulum and Golgi apparatus, suggesting activation and hypersecretory transformation of cardiac fibroblasts in the presence of IL-1β (Figure S14). Increased intracellular and budding extracellular microvesicles was seen in the IL-1β-treated cells. We next sought to expand the number of cardiac fibroblasts with T-cell features by growing the cells in T-cell expansion media for 10 days with added IL-1β (Figure S14). We noted the formation of cell clusters in all the human donors, suggesting that IL-1β induced phenotypic changes in cardiac fibroblasts from day 7 onwards (Figure S14). Taken together, shorter incubation with IL-1β.increased proliferation and longer incubation resulted in differentiation of hVCF into a secretory cell and distinct cell clustering.

**Figure 4:**
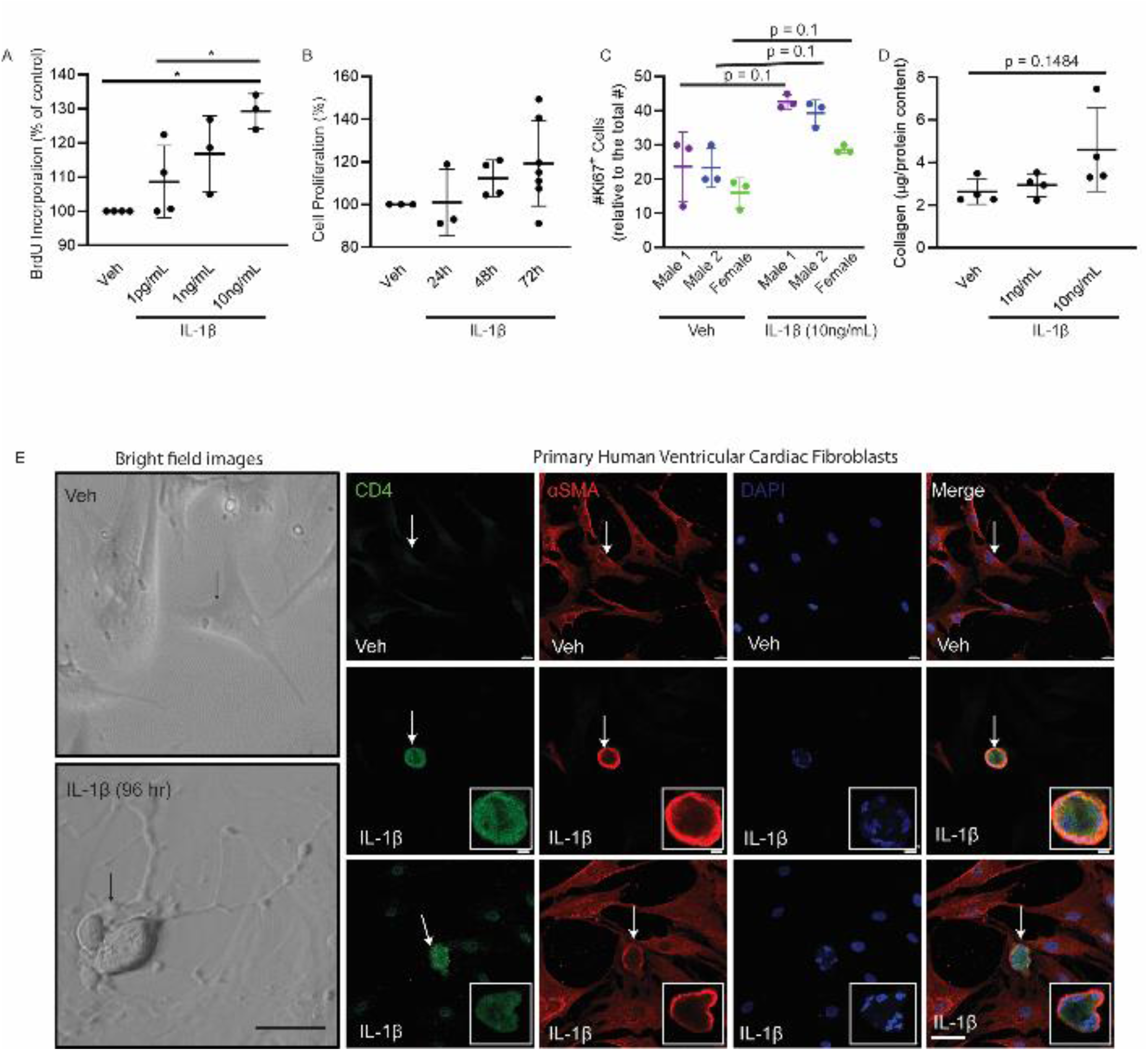
IL-1β-mediated proliferation, transdifferentiation and activation of primary human ventricular cardiac fibroblasts (hVCF) into CD4-expressing cells *in vitro*. Primary hVCF cells (2.0 x 10^6^ cells/mL) from donors (2 males (ID# 62122, ID# 1281202) and 1 female (ID#534282) were cultured in Fibroblast Basal Media (FBM) for 4 days with 10ng/ml of IL-1β or in T-cell media with CD3/CD28 T-cell activator for 13 days. (Α) hVCF proliferation in response to IL-1β exposure was assessed by BrdU incorporation. Data are Mean ± SD for 3 biological replicates. \**P* <0.05, one-way ANOVA with multiple comparisons. (B) Cell proliferation response of hVCF to IL-1β after 24 h, 48 h and 72 h of incubation was assessed using MTT assay. Data are Mean ± SD for 3 biological replicates (n=6 technical replicates). (C) Cell proliferation response to IL-1β determined by Ki67 staining of nuclei. Data are Mean ± SD for 4 biological replicates, *P*=0.1, one-way Kruskal-Wallis ANOVA with Dunn’s multiple comparisons. (D) Collagen content in response to IL-1β doses (1ng/mL and 10ng/mL) measured in the cell lysates using the Sircol assay. The data are represented as Mean ± SD values from individual subjects (n=4) biological replicates, no significance (ns), Kruskal-Wallis one-way ANOVA with Dunn’s multiple comparisons. (E) Left Panel: Bright-field images of live hVCF cells indicating the shifts in cellular morphology from a spindle-shaped fibroblast to a round cell with a large nucleus in response to 96 h of treatment with IL-1β (10ng/mL). Arrows indicate a change in morphology of hVCF with IL-1β treatment, but not with vehicle treatment. Right Panel. Immunostaining of fixed hVCF with lymphoid CD4 T-cell marker (green), mesenchymal αSMA (red) and DAPI (blue) markers to characterize the transformed cells. The arrow indicates the emergence of round, multinucleated αSMA^+^ CD4^+^ expressing cells. Representative magnified images of multinucleated notch-shaped giant cells immunostained with αSMA^+^ CD4^+^. Quantification of the bright-field images of differentiated cells counted per field (n=3 biological replicates). The data are represented as Mean ± SD values from individual subjects; (n=3) biological replicates, \**P*<0.05 and Kruskal-Wallis one-way ANOVA with Dunn’s multiple comparisons.

### Expression of extracellular matrix genes and inflammatory genes in response to inflammatory cytokine

One of the key functional features of immune cells is their ability to secrete cytokines, chemokines and immunomodulatory proteins. To determine whether hVCF expressed both the mesenchymal cell and inflammation associated genes with IL-1β, we used a qPCR approach of gene expression quantitation using a validated taqman array plate preset with primers specific to extracellular matrix genes or inflammatory genes. A comprehensive list of both extracellular matrix genes and inflammatory genes modulated by IL-1β is presented as a heatmap (Figure 5, Panel A and Panel B) and as a supplementary table (Table S6 and Table S7). Inspite, of the heterogeneity among the donor cells, clear distinction was made between untreated cells and cells treated with IL-1β for both the arrays (Figure 5, Panel A and Panel B). Notably, the *CD44* gene, highly expressed during T-cell development, is upregulated by 3-fold in IL-1β-treated cardiac fibroblast cells (Table S7). Inflammatory gene expression data analyzed using string techniques predict interaction between the *IL-1R1* gene and the inflammatory regulator Nuclear Factor kappa-light-chain-enhancer of activated B cells (*NFkB1)*(Figure 5, Panel C and Panel D). In addition, Annexin A1 (*ANXA1),* which is modulated with IL-1β, is predicted to interact with multiple binding partners, such as Platelet Activating Factor Receptor (*PTAFR),* Leukotriene B4 receptor 1*(LTB4R)* and Histamine Receptor *H1 (HRH1)*. Extracellular matrix gene expression data showed a cluster of Integrin Subunit Alpha 1 (*ITGA)* modulated by IL-1β interacting with binding partners and CD44 interacting with Matrix metalloproteinases (*MMP14)* and Versican *(VCAN). MMP1* is predicted to interact with *MMP11* and Alpha 2 Macroglobulin *(A2M)* (Figure 5, Panel C). Further, pathway analysis using CYTOSCAPE suggested that immune responsive genes with false discovery rate (FDR) of 2.85 x 10^-13^ belonged to gene ontology terms “cellular response to cytokine stimulus”, “cytokine-mediated signaling pathway”,” inflammatory response”, “defense response and immune system process” (Figure 5, Panel C), while extracellular matrix-associated genes with the FDR of 2.13 x 10^-17^ belonged to gene ontology term “extracellular matrix organization” (Figure 5, Panel D). We further validated the gene expression of *CCR2, IL-1R, IL-8* and *TGF-β* (Figure 5, Panel E) and CCL2, IL-6, IL-1R, MyD88, ColI-1, Col III, and MMP-9 (Figure S15, Panel A) in primary adult rat ventricular fibroblasts isolated using Langendorff’s method and expression of IL-2, IL-8, IL-10, ICAM, and TGF-β in primary human ventricular fibroblasts using qPCR (Figure S15, Panel A).

**Figure 5:**
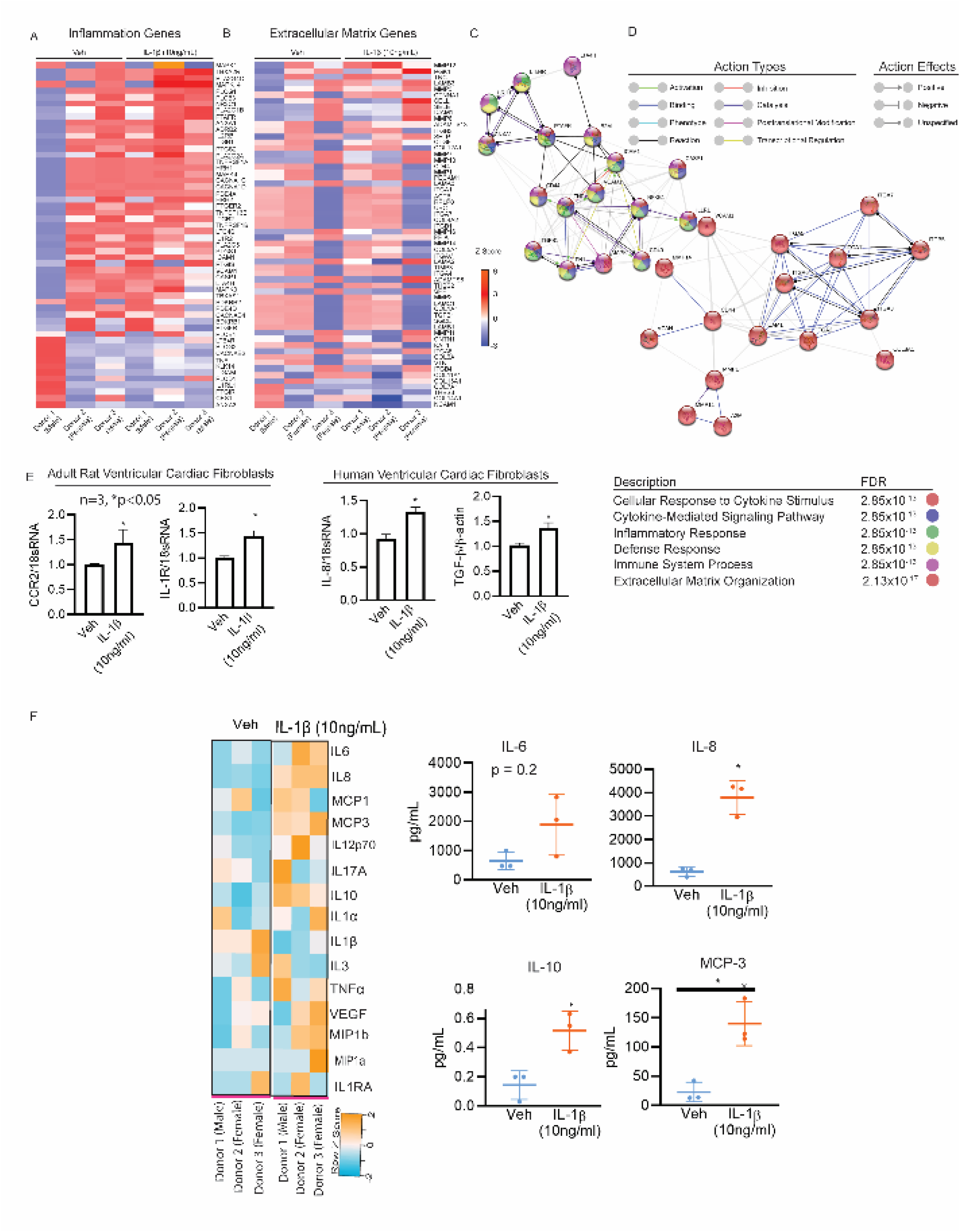
IL-1β modulates inflammatory and extracellular secretome in hVCF: Genomic profiles. Cardiac fibroblasts were treated with IL-1β for 96h, total mRNA was extracted, and differential regulation was measured. Heatmaps and Z-scores representing differential gene expression between vehicle- and IL-1β-treated cells affecting (A) inflammation and (B) extracellular matrix remodeling using R package (heatmap.2). Red denotes high expression, and blue denotes low expression. String pathway analysis using CYTOSCAPE of genes related to (C) inflammation and (D) extracellular matrix remodeling regulated by IL-1β. Gene networks represent undirected weighted partial correlation network constructed as a force directed graph. The network nodes represent individual genes that are differentially expressed. Edges connect the modes, and edge weights are colored based on relative activity. (E) Assay of genes involved in inflammation and extracellular matrix, such as CCR2 and IL- 1R, was performed in primary adult rat ventricular cardiac fibroblasts. Further assay of genes involved in inflammation (IL-8, IL-10, and TGFα) was performed in primary hVCF treated with Veh or IL-1β (10ng/mL) after 24h using quantitative PCR (qPCR). The graph represents Mean ± SD values (n=3, \**P*<0.05 by unpaired Student’s *t* test with Mann Whitney post hoc test).

Functional gene ontology (GO) enrichment analysis of the secreted protein using the g:profiler web server showed humoral immune response, immune effector process, leukocyte-mediated immunity, vesicle-mediated transport, platelet degranulation, neutrophil activation and lymphocyte-mediated immunity as highly significant for a search of biological processes (BP) (Figure S16, Panel B). A search for cellular component (CC) enrichment showed that the secreted proteins were predicted to be a part of extracellular exosome, collagen-containing extracellular matrix, extracellular matrix, secretory granule lumen and cytoplasmic vesicle lumen (Figure S16, Panel B).

### Induction of immunomodulatory proteins in response to inflammatory cytokine

Next, we traced the secreted proteomic signatures (immunomodulatory cytokines, chemokines, proteins and metabolites) in the conditional media, using quantitative cytokine and chemokine analysis, proteomics, and metabolomics. IL-1β-dependent-cytokine and chemokine secretion was analyzed using Milliplex MAP Human Cytokine/Chemokine Magnetic Beads in the presence of cardiac fibroblasts from three human donors. We noted that 24 h treatment with IL-1β significantly increased secretion of MCP- 3, IL-8 and IL-10, tended to increase secretion of IL-6, IL-12p70, TNF-β, and VEGF, but tended to decrease IL-1β levels, although these results were not statistically significant (Figure S15, Panel C).

Unlabeled proteomics using LC MS/MS to identify secreted proteins showed a total of 525 and 588 proteins secreted in the conditioned media of cardiac fibroblasts treated with Veh or IL-1β (10ng/mL) (Figure S16, Panel A and Tables S8-S11). Out of the annotated 394 proteins, the Venn diagram demonstrated that 120 proteins were unique to IL-1β and that 181 proteins were common to both Veh and IL-1β (Figure S16, Panel B). Conditioned media replicates for the Veh and IL-1β groups were internally consistent, as represented in Figure S16, Panel C. Gene ontology as analyzed with PANTHER showed terms synonymous with extracellular matrix regulation and inflammation in response to IL-1β. Tables S6 to Table S9 summarize the functions of designated proteins in terms of biological processes, molecular function and immune subset population identified uniquely in the conditioned media of IL-1β-treated hVCF. Gene Ontology (GO)-based annotation of proteins uniquely found in the conditioned media from cardiac fibroblasts treated with IL-1β (Tables S6-S9) showed that the proteins shown were involved in the activation of both adaptive and innate immune responses (GO: 0042102, Positive regulation of T-cell proliferation; GO: 0006959, Humoral immune response; GO: 0002443, Leukocyte-mediated immunity; GO: 0002252, Immune effector process; GO: 0002576, Platelet degranulation; GO: 0006956, Complement activation; GO: 0002274, Myeloid leukocyte activation and GO: 0002446, Neutrophil-mediated immunity) and recruitment of inflammatory and reparative immune cells (CXCL1, CCL2, CXCL3, CXCL5,CXCL8 apolipoprotein A1, IL-6, complement factor H related (CFHR1), and VCAM1). Specifically, proteins involved in the activation of innate and adaptive immune responses, leukocyte activation, and leukocyte migration were displayed in the biological process analysis using PANTHER (Figure S16, Panel E).

### Inflammatory cytokine IL-1β induced secretion of immune response-associated metabolites

Metabolic reprograming and metabolic flexibility is required for phenotype switching in response to environmental milieu. We therefore analyzed the metabolites in the conditioned media, following treatment with IL-1β, to determine the overall changes in the metabolic status of cardiac fibroblasts, using nontargeted metabolite profiling of the conditioned cell culture media. Principal component analysis based on overall differences between the metabolites indicates separation of vehicle group and IL-1β-treated groups (Figure 6, Panel A). The volcano plot distinguished metabolites with negative log with a false discovery rate higher than 1 modulated by IL-1β treatment (Figure 6, Panel B). Interestingly, the metabolic profile for the female donor of the cardiac fibroblast cells with and without IL-1β treatment was different compared to the male donors (Figure 6, Panel C). The top 25 differentially upregulated and downregulated metabolites corresponding to metabolic shifts from inactive cell to metabolically active cells are represented on the heatmap (Figure 6, Panel D). Pathway analysis of the differentially expressed metabolites indicated these metabolites involved in cardiolipin biosynthesis, a key inner mitochondrial membrane protein and mitochondrial oxidant producer, were significantly upregulated in IL-1β exposed cardiac fibroblast cells (Figure 6, Panel E). Typical CD4^+^ metabolite signatures of activated T cells involving methionine metabolic pathways and pyrimidine metabolism were also upregulated. Methyl histidine metabolism, phenylacetate metabolism, and *de novo* triacyglycerol biosynthesis were also upregulated (Figure 6, Panel E). Interestingly, metabolites serving as a link between glycolysis and citric acid cycle, including thiamine pyrophosphate, were differentially secreted in response to IL-1β stimulation. Specifically, metabolites associated with ATP generation, carbohydrate metabolism, and the production of amino acids, nucleic acids, and fatty acids, such as spermidine, spermine and atrolactic acid, were upregulated, while others were downregulated, such as sedoheptulose-1-7-phosphate, oxaloacetate, N6-acetyl-L-lysine, succinate, 1,3 diphosphate glycerate, and citrate-isocitrate (Figure 6, Panel F and Table S12-S13). In sum, our data suggests that cardiac fibroblasts are activated by IL-1β and transform into a immune cell in the presence of IL-1β by upregulating inflammatory and extracellular matrix genes and by secreting cytokines, chemokines, and immunomodulatory proteins and metabolites involved in both innate and adaptive immune response critical in both pathogenesis and resolution of injury.

**Figure 6:**
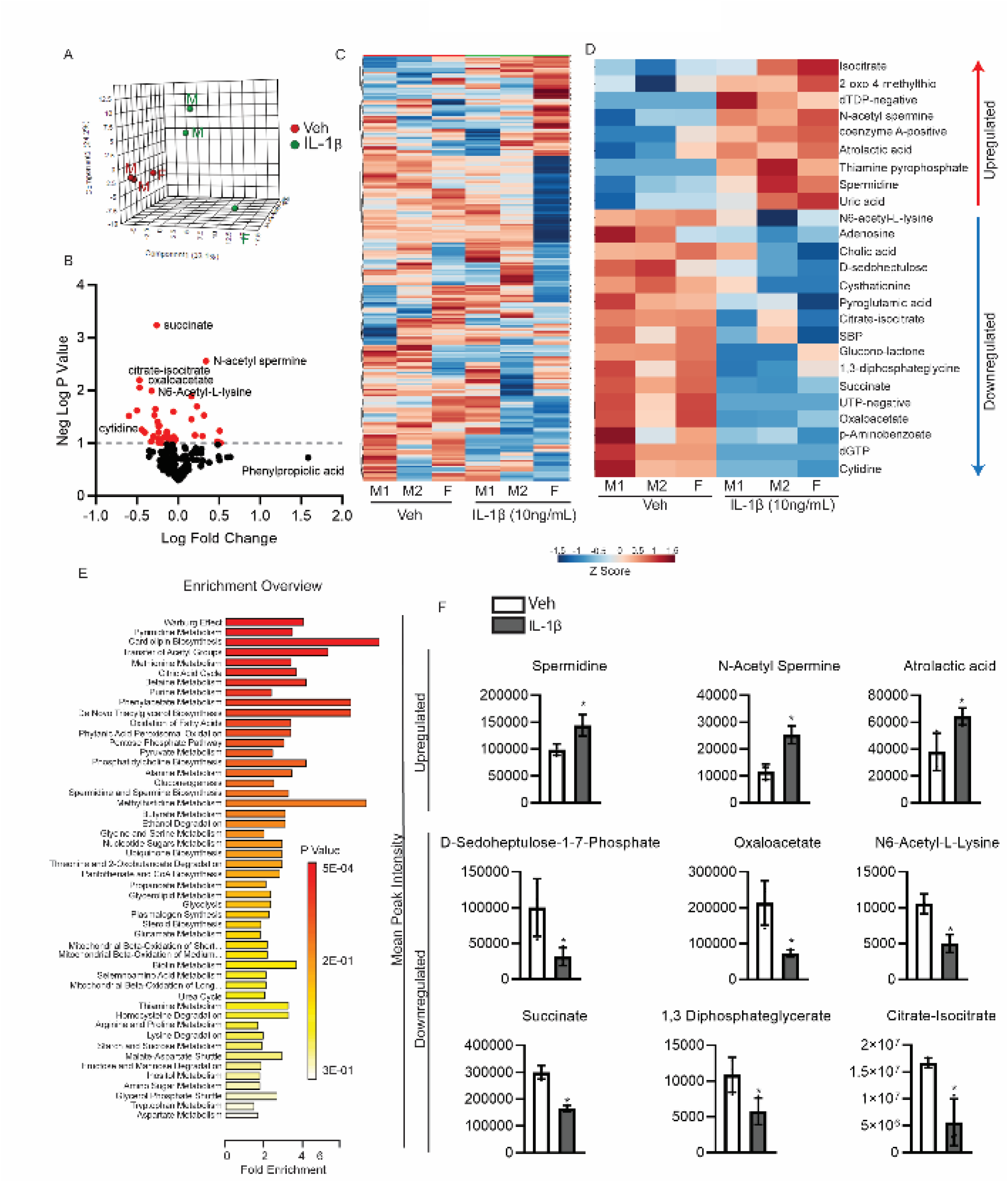
IL-1β modulates inflammatory and extracellular secretome in hVCF: Metabolomic profiles. (A) Cardiac fibroblasts were treated with IL-1β for 96h and principal component analysis of a cluster of metabolomic profiles relative to control. Multivariable metabolite profile of individuals (ID# 62122; male, ID# 1281202; male and ID#534282; female) was reduced to a single dot and represented based on consensus clustered alignment. Principal component 1 (PC1) on the x-axis versus principal component 2 (PC2) on the y-axis. (B) Volcano plot showing global expression of secreted metabolites in the conditioned media positively and negatively modulated upon treatment with IL-1β ng/mL for 24 hr. Each dot represents a differentially expressed metabolite. The horizontal line marks a P value of 0.01, and the red dots represent significant metabolites. (C) Heatmap columns comparing the expression of secreted metabolites identified by unsupervised hierarchical consensus clustering. Each row represents individual metabolites identified in the conditioned media of vehicle- and IL-1β-treated cells. The displayed color codes represent Z scores (SD above or below the mean); n=3 biological replicates. (D) Top 25 metabolites showing a significant expression of upregulated or downregulated metabolites (*P <0.05) between Veh- and IL-1β-treated cell conditioned media) are represented as a heatmap; n=3 biological replicates. (E) Pathway enrichment analysis of the metabolites differentially expressed with IL-1β treatment. Length of the bar corresponds to fold enrichment, and the color suggests the significance of the pathway based on the *P* value. (F) Quantification of the metabolites is represented as Mean ± SD from 3 different human donors. (\**P* <0.05, two-tailed *t* test, Mann Whitney post-hoc test).

### Transcriptome profile of IL-1β-induced CD4^+^ fibroblast cell population

To further characterize the identity of this unique cell population and based on the box-plot evidence of increased CD4^+^ population with IL-1β (Figure 7, Panel A), we isolated this population by FACS sorting and mapped the transcriptomic signatures using RNA sequencing. hVCF cells were flow sorted into CD4^+^ cardiac fibroblasts and CD4^-^ cardiac fibroblasts for both Veh and IL-1β groups after 10 days treatment in T-cell differentiation media, and then the genetic footprint of the newly identified CD4^+^-expressing cardiac fibroblasts traced using next-generation deep total RNA sequencing (Figure 7 and Figure S17). Unique transcriptomic signatures of the IL-1β CD4^+^ hVCF population (Figure 7, Panel B) and the differential gene expression between Veh and IL-1β groups were observed corresponding to metabolic changes, cell development and differentiation and T-cell signatures (Figure 7, Panel B).

**Figure 7:**
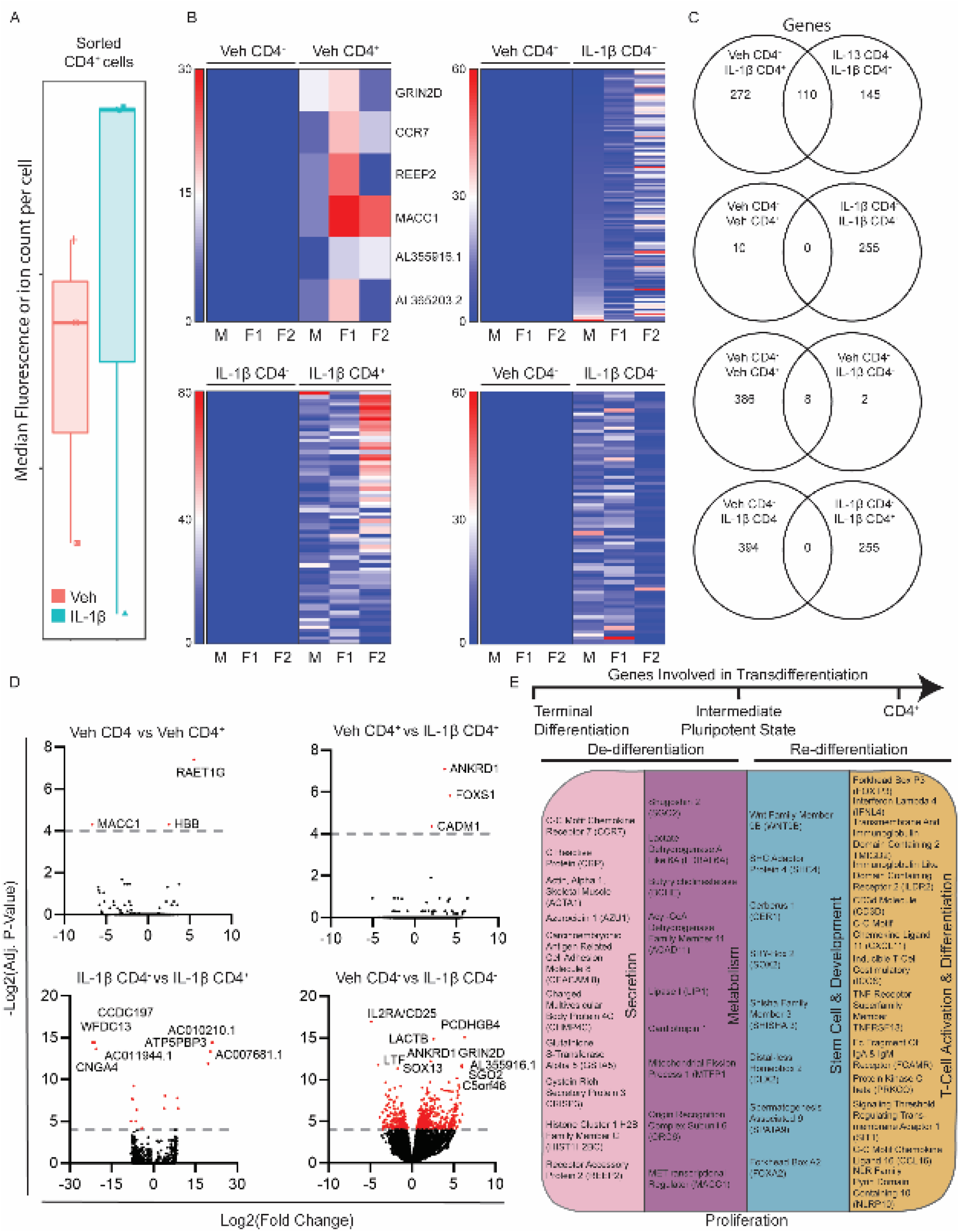
Genetic characterization of differentiated CD4^+^cardiac fibroblast cells. Primary hVCFs from 3 human donors (males and female) were differentiated for 10 days in T-cell media with or without IL-1β (10ng/mL) and flow sorted into CD4^+^ and CD4^-^ populations, followed by next-generation RNA sequencing using the illumina platform. (A) Box plot of shifts in CD4^+^ human cardiac fibroblast population in response to IL-1β. (B) Heatmaps showing the expression pattern of genes uniquely identified in Veh CD4^+^, Veh CD4^-^, IL-1β CD4^+^ and IL-1β CD4^-^ populations. (C) Venn diagram showing the number of common genes among the Veh and IL-1β among the unique gene identified for all the populations. (D) Volcano plot showing the number of significantly upregulated and downregulated genes for each of the comparisons. False Discovery Rate (FDR) (adjusted P value <0.01). (E) Summary of gene ontology hits using database for annotation, visualization, and integrated discovery (DAVID) online platform showing the enrichment of genes involved in secretion, pluripotency, development, metabolism and T-cell activation and differentiation. Benjamini Hochberg score of false discovery rate.

Heatmaps based on nonhierarchical clustering showed differences in transcriptomic signatures for the IL-1β CD4^+^ and IL-1β CD4^-^ populations (Figure 7, Panel B). However, only 6 genes were unique to Veh CD4^+^ compared to the Veh CD4^-^ group (Figure 7, Panel B). Venn diagrams further showed no common genes shared by the newly identified Veh CD4^+^ and IL1β CD4^+^ populations (Figure 7, Panel C). On the other hand, 255 genes were unique to the IL-1β group (Figure 7, Panel C). The number of significant genes common to all four groups were represented by the volcano plot (Figure 7, Panel D). Notably, 22 genes were significantly down-regulated, and 31 genes were significantly upregulated in the IL-1β CD4^+^ population. The CD4^+^ cardiac fibroblast population differed from the parent population by possessing a stem cell-like activated state, as evidenced by the presence of genes associated with pluripotency and reprogramming (*WNT9B, SHC4, CER1, SOX2, SHISHA3, DLX2, SPATA9 and FOXA2)*. Moreover, CD4^+^ stem cell-like cells were secretory, in contrast to the CD4^-^ cells, as evidenced by their expression of microvesicle- and exosome-associated genes (*CCR7, CRP, ACTA1, AZU1, CAECAM 8, CHMP4C, GSTA5, CRISP3, HIST1H2BC and REEP2*). In addition, these stem cell-like secretory cells were metabolically active (*SGO2, LDBAL6A, BCHE, ACAD11, LIP1, CT-1, MTFP1,ORC6 and MACC1*) and expressed characteristics of lymphoidal cells (*FOXP3, CD3D, IFNL4, TMIGD2, ILDR2, CXCL11, ICOS, TNFRSF18, FCAMR, PRKCQ, SIT1, CCL16 and NLRP10*), as annotated based on their biological function using Database for Annotation, Visualization, and Integrated Discovery (DAVID) (Figure 7, Panel E). Innate immunity-linked genes (2’-5’-oligoadenylate Synthase Like (OASL), absent in melanoma 2 (AIM2)) and adaptive immunity-linked genes (CD1c molecule, CD207 molecule), co-stimulatory signal during T-cell activation (inducible T-cell costimulatory (ICOS)), T-cell chemotaxis (CXCL11), TNF receptor superfamily member (TNFRSF18) involved in leukocyte migration, B-cell proliferation genes (GRB2-binding adaptor protein) and genes associated with phagocytosis (carcinoembryonic antigen-related cell adhesion molecule 4 (CEACAM4) were unique to the IL-1β CD4^+^ group (Figure 7, Panel E). These experiments confirmed that a subset of terminally differentiated resident cardiac fibroblasts may convert into lymphoid-like cells in the presence of the inflammatory cytokine, IL-1β.

### Transdifferentiation of cardiac fibroblasts into lymphoid-like cells is mediated by IL-1R-pMAPK and IL-1R-NFkβ pathways

IL-1β binds to the Interleukin-1 receptor (IL-1R) to execute its downstream expression and activity. We found that IL-1R gene expression increased significantly in response to 24 hour treatment with IL-1β (10 ng/mL) (Figure 8, Panel A). We also observed a dramatic increases in IL-1R puncta on micrographs taken using confocal microscopy and Z-stacking of images followed by 3D projection, both in the nucleus and cytoplasm. In addition, we observed secretion of IL-1R outside the cells, visualized using a 3D rendering of the confocal images obtained from immunostained IL-1β-treated hVCF (Figure 8, Panel B). In addition, IL-1R protein expression and foci formation were significantly reduced with IL- 1Ra (1 µg/mL). Differentiation of cardiac fibroblasts in T-cell media for 14 days showed that the cell clusters typical of IL-1β treatment were significantly reduced with IL-1RA (Figure 8, Panel C). We next assessed the activation status of IL-1β-driven downstream signaling pathways involved in proliferation and differentiation, i.e., p38 MAPK and NF-κB p65. We showed that treatment with IL-1β increases prodifferentiation p38 MAPK phosphorylation at Thr180/Tyr182 residues (Figure 8, Panel D). This effect was inhibited by incubation with IL-1RA, but not rescued by IL-1β. Similar results were noted with NF-κB p65 phosphorylation at Ser 536 residue in 2 out of 3 instances, although these results did not reach statistical significance. In sum, IL-1β promotes differentiation of mesenchymal cardiac fibroblast to cells structurally and functionally similar to T-helper cells through the IL-1R/p38 MAPK pathway assessed using human cell, tissue, rat disease models and sensitive recent methodologies (Figure 8, Panel E and Figure S18).

**Figure 8:**
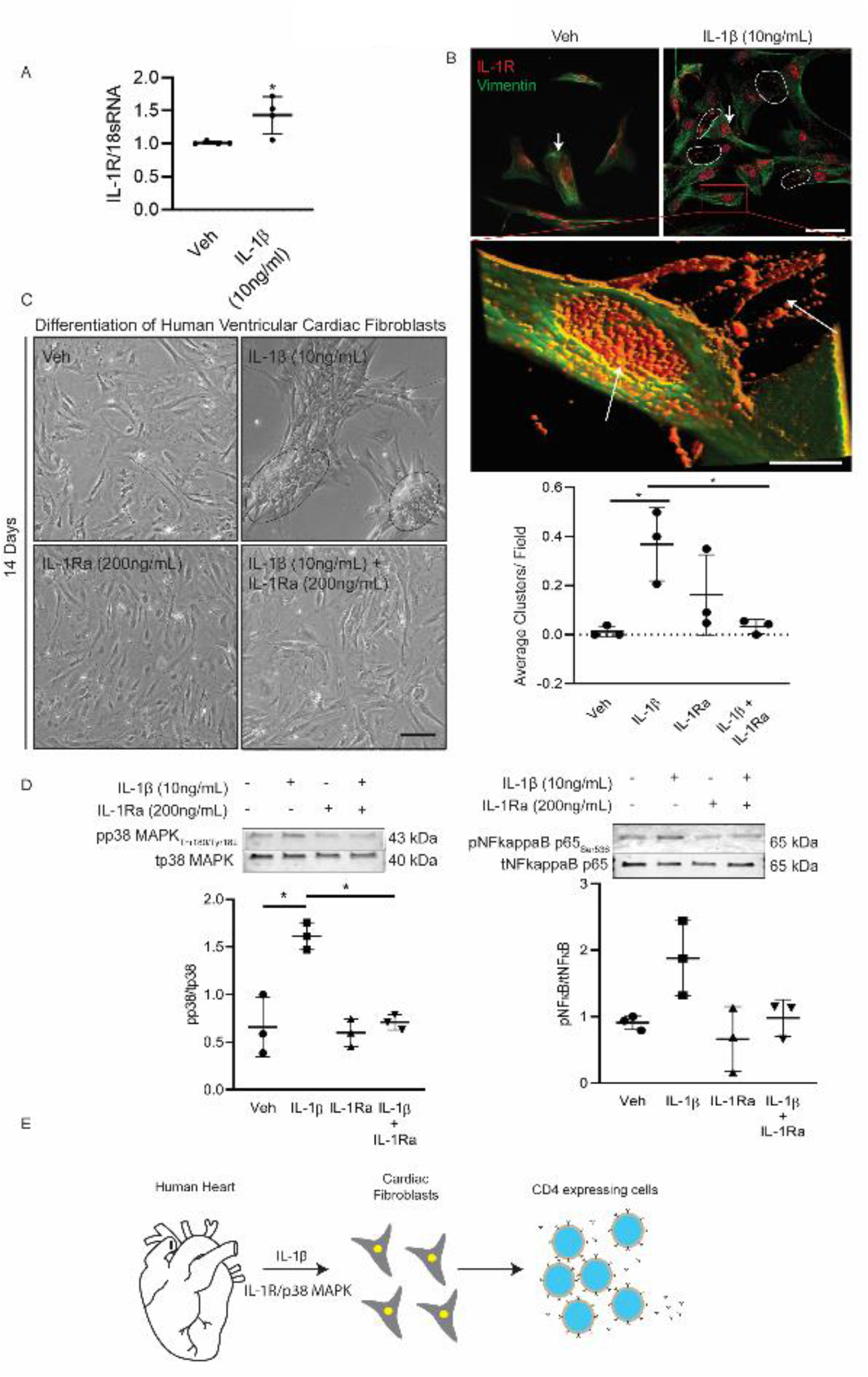
IL-1β-mediated transdifferentiation of primary human ventricular cardiac fibroblasts (hVCF) into cells with CD4 T-cell markers occurs through the IL-1R-pMAPK and IL-1R-NFkB pathways. Primary hVCFs from 3 human donors (males and female) were treated with Veh, IL-1β (10 ng/mL), IL-1RA (200 ng/mL) and IL-1β +IL-1RA for 24 h. (A) Quantification of IL-1R gene expression determined using qPCR was performed in primary adult rat cardiac fibroblasts (Left). The data represent Mean ± SD values from 3 biological replicates. \**P*<0.05 unpaired two-tailed Student’s -*t* test. (B) Representative images of IL-1R expression and distribution across the nucleus and cytoplasm of hVCF treated with Veh or IL-1β. The white dotted lines indicate the bright red foci of IL-1R outside hVCFs after treatment with IL-1β. Z stack images of IL-1β-treated hVCFs were projected in 3D using image J. Arrows indicate overexpression and extracellular release of IL-1R. (C) Bright-field images of hVCF differentiation in T-cell media after 14 days of treatment with Veh, IL-1β ng/mL, IL-1RA (200 ng/mL) or IL-1β +IL-1RA. Quantification of rosette clusters per field performed by a double-blinded observer on randomized images. The black lines indicate the differentiated cell clusters formed in response to IL-1β after 14 days. (D) Phosphorylation of pMAPK_Thr180/Tyr182_ and pNFkBp65_Ser536_ upon 24 h stimulation of hVCFs with Veh, IL-1β (10 ng/mL), IL-1RA (200 ng/mL) or IL-1β + IL-1RA, as determined by immunoblotting. The data are presented as scatter plot of Mean ± SD values from individual donors and technical replicates (n=3, \**P* <0.05 Kruskal-Wallis one-way ANOVA with Dunn’s multiple comparisons). (E) We propose that resident cardiac fibroblasts undergo staged polarization/phenotypic switching in response to proinflammatory signals, such as IL-1β, to facilitate repair and cell survival. This results in transdifferentiation to a secretory cell with a lymphoid cell-like phenotype that participates in the amplification and regulation of the inflammatory response through secretion of immunomodulatory proteins and metabolites.

## Discussion

Myocardial injury is characterized by complex interactions involving mesenchymal cardiac fibroblast and lymphoidal immune cells that collectively form the backbone of regenerative repair in the heart (*48, 49*). Despite the significance of cardiac fibroblast in cardiac diseases, very little is known about these cells, their origin, activation states, transitory states and their role in inflammation and repair. Partly, due to the lack of sensitive techniques that could resolve the complexities at a single cell levels. Here, we have used advance cell profiling techniques such as mass cytometry and powerful algorithm to identify a previously unknown sub-population of cardiac fibroblasts with dual mesenchymal and lymphoidal features. Further employing an array of complementary methods in isolated human cells, tissues and rat model of SUGEN/Hypoxia, we show that this unique population is localized in fibrotic regions and structurally and functionally mimick the mesenchymal and lymphoidal cells evidenced from a prototypical fibroblast-immune secretome (genome, proteome, metabolome) analysis. Blocking of the IL-1β receptor using an IL-1R antagonist *invitro* reverses IL-1β-mediated clustering of cardiac fibroblast cells and p-p38 MAPK phosphorylation suggesting a receptor mediated downstream activity. Lastly, characterization of the IL1-β induced CD4^+^ population using deep RNA sequencing suggests that these unique population possess signatures of development, metabolism, T-cell markers, stem cell markers. These findings, taken together, are consistent with a novel role for transdifferentiated resident fibroblasts in carrying out a “first response” to pressure-related cardiac injury through immune modulation of fibrosis adding to previous finding of matrix deposition after injury ((3), *44, 50*).

Previous studies have shown that cardiac fibroblasts are also capable of assuming an inflammatory role whereby they secrete immune cell-recruiting cytokines and chemokines (*53*) and enact phagocytosis (*54*) in the setting of myocardial injury. However, these cells were not characterized. The lack of detection of these cells so far may be simply related to not looking for this CD4^+^ cardiac fibroblast population in injured myocardium. Immune dysregulation involving CD4^+^ helper T-cell type I (Th1)/Th17 immunity is one of the key attribute of PH. Modulating CD4^+^ Treg cells reduces endothelial injury and prevents PH in T-cell-deficient rats (*55, 56*). Mature CD4^+^- depleted mice are protected from left ventricular fibrosis, but not cardiac hypertrophy in transverse aortic constriction (TAC)-induced left ventricular failure (27). However, the mechanisms by which CD4^+^ Treg cells potentiate or alleviate fibrosis are unclear. It is speculated that CD4^+^ Treg cells recruited from the bone marrow secrete extracellular matrix similar to cardiac fibroblasts which promotes thickening and reduces compliance of the ventricular wall, leading to systolic and diastolic dysfunction. Similarly, resident myofibroblasts may be activated or recruited in response to damage-associated molecular patterns (DAMPS) from the damaged myocytes to accelerate the process of extracellular deposition and healing; however, this theory is debated (26). Yano et al. suggest that resident cardiac fibroblasts, but not bone marrow-derived cells, add to the myofibroblast pool in the setting of myocardial repair (*57*).

The surge in the numbers of myofibroblasts following myocardial injury is unaccounted for, and the role of resident cardiac fibroblasts in myofibroblast accumulation is not known (*58*).

Recent attempts using periostin and TCF-21 mouse models have been made to identify and trace the lineage of the sudden expansion and activation of fibroblast cell populations after myocardial infarction (*59, 60*). Results employing these models demonstrate that circulating bone marrow-derived cells, CD14^+^ fibrocytes, smooth muscle cells, and endothelial cells transform into cardiac myofibroblasts through endothelial to mesenchymal transitioning, which may constitute a source of myofibroblasts and deposition of excess of extracellular matrix (*20, 61–63*). Both CD4^+^T cells and cardiac fibroblasts are identified independently in fibrotic regions secreting cytokines, chemokines and extracellular matrix deposition and remodeling. In response to proinflammatory stimuli, such as IL-1β, our data suggest that resident cardiac fibroblasts respond by genetic reprogramming and phenotypic switching into CD4^+^ cells in order to recruit further immune cells and initiate repair. T-cell macrophage cell fusion is common in the setting of myocardial injury, resulting in the formation of giant cells. We show the appearance of notch-shaped nonhomogeneous giant cells after treatment of cardiac fibroblasts with IL-1β for 4 days, confirming yet another feature of T-cells. Our RNA- seq data further demonstrate that CD4^+^ cardiac fibroblasts exhibit features of mesenchymal cells, T-cells, pluripotent cells and metabolism in IL-1β- treated, but not untreated, cells, suggesting that terminally differentiated cardiac fibroblasts may dedifferentiate into a stem cell-like metabolically active state and then redifferentiate into lymphoidal cells in response to IL-1β, further opening up a new possibility of regenerating damaged cells and cell survival (*64*). In fact, the epigenetic and metabolic reprogramming seen in the CD4^+^ population is reminiscent of the induction of lymphocyte-dependent adaptive immune responses and trained immunity (*65*). The pathways involved in cardiac fibroblast activation, proliferation, and differentiation in response to IL-1β are currently unknown. One possible mechanism is that IL-1β binds to the IL-1R1 receptor and thereby recruits MyD88 adaptor protein, which, in turn, results in the activation of the central inflammatory mediator, NFkβ, and the synthesis of IL-1, IL-6 and TNF-α (*38*). The enlistment of the IL1R1/MyD88/NFkβ pathway in the manner shown thus implies a role of this sequence of events in the genesis of cardiac fibrosis and pulmonary hypertension (*38*). Deciphering the preferential pathways linking the activation of the IL-1R1 receptor with transdifferentiation under varied pathological conditions will be the focus of future research.

Most of the studies related to cardiac fibroblasts function in cardiac disease states is performed in mice and rats. We acknowledge that while most of the work is performed in cultured cells, these are primary cells obtained from humans through optimized protocols. The findings were confirmed in autopsy human RV tissue from patients with Pulmonary Hypertension and cardiac fibrosis. While, we tried isolating cardiac fibroblasts from the rat RV and LV, sufficient numbers were not obtained from fibrotic tissue to repeat all the characterization. And based on prior studies, it is possible that the cells have lost their nascent properties. Studies elucidating the functional role of transdifferentiated myofibroblasts *in vivo* in different tissues, along with the analysis of lineage, fate, epigenetic changes, and regulation of IL-1R downstream signaling mediators and promotors at gene, protein, metabolism levels, as well as interaction with surrounding cells, is beyond the scope of this work. We further acknowledge that the regulation of cell phenotypic switching in response to tissue perturbations, such as those depicted here, is likely to be exceedingly complex, and involve multiple cell types and cytokine pathways and would require several transgenic animal models and significant resources. Moreover, compensatory mechanisms affecting the transdifferentiation of resident cardiac cells into immune-like phenotypes may involve the orchestration of multiple cell types, growth factors, tissue repair agents, metabolite consumption and secretome release. While we are not excluding the involvement of other cytokines and chemokines in the transdifferentiation of the cardiac fibroblasts into lymphoid lineages *in vivo*, our work indicates that IL-1β alone is sufficient to transdifferentiate 2-3% of the cardiac fibroblast population into CD4^+^ cells, possibly through dedifferentiation into stem cell-like cells and then redifferentiation. Lastly, we have not addressed the possibility that the process of transdifferentiation may, in fact, be bidirectional, with phenotypic switching from cardiac fibroblasts to lymphoid cells coupled with phenotypic switching from lymphoid cells back to cardiac fibroblasts. Nonetheless, we have characterized through the use of complementary *in vitro,* and *in vivo* methods the presence and biological significance of these unique CD4^+^- expressing cells in the adult heart (Figure S18).

In conclusion, we have identified a unique population of cells which appears to bridge the characteristic biology of fibroblasts and lymphoid cells during tissue repair. Recognizing the potential role of fibroblast plasticity in orchestrating the conversion, enlistment, and activation of an array of inflammatory cell types to the stressed myocardium, therapeutic methods designed to modulate cardiac fibroblast plasticity-related, inflammation-induced right ventricular fibrosis may emerge in the future. Since right ventricular failure is the most common cause of mortality in PH, modulation of fibroblast plasticity raises the possibility of new approaches to therapy by regulating phenotype switching, promoting the reparative phenotypes in injured regions and preventing the transition to maladaptive remodeling through de-differentiation and re-differentiation of adult cells.

## Materials and Methods

### (i) Human

#### Human Tissue Material Characteristics

De-identified transverse right ventricular tissue sections of 10μM thickness were obtained post-mortem from patients with a diagnosis of PH. Control tissue samples were obtained from patients who died of unrelated diseases. Samples were obtained from the Yale Tissue Repository Service, New Haven, Connecticut. Microscopically, all of the PH myocardial tissue of right ventricles showed multiple foci of fibrosis and scar consistent with chronic ischemic damage. Interstitial fibrosis was observed within the posterior papillary muscle of the left ventricle. Endocardial surfaces of both ventricles, especially right ventricle, showed hypertrophic changes with notable big box car nuclei. The cardiovascular autopsy reports for all cases (A, B, C, D, E, and F) are provided in Table S4. The IRB approvals were obtained from the Providence VA Medical Center IRB committee.

#### Immunohistology of Human Right Ventricular Tissue

Transverse sections (10μM thick) of formalin-fixed, paraformaldehyde-embedded (FFPE) tissues from the apex of right ventricle from individuals diagnosed with PH or controls were stained with hematoxylin & eosin (H&E). Unstained human right ventricular tissue sections were subjected to immunohistochemistry detecting CD4 (Cat # ab133616, clone EPR6855). The FFPE sections were de-paraffinized in xylene, dehydrated in 100% ethanol, 95% ethanol, 70% ethanol, 50% ethanol and rehydrated in water. Antigen retrieval steps to unmask the antigens were performed using Tris-EDTA buffer (100mM Tris base, 1mM EDTA, 0.05% Tween-20, and pH adjusted to 9.0) and steamed at 100°C for 20 min. The endogenous peroxidase activity was blocked using 0.3% peroxide solution for 30 min. The slides were incubated in 2.5% normal serum for 1h at room temperature. CD4 primary antibody was used at a concentration of 1:500 and incubated at 4°C overnight. Slides were washed, incubated with secondary antibody for 30 min, and visualized using 3, 3’ diaminobenzidine peroxidase substrate system. Morphometry was performed on both the round and spindle shaped cells expressing CD4. The nucleus was counterstained with methyl green. Pathological assessment was done in the random order on the entire right ventricular tissue sections by a blinded observer. Nuclei were quantified using orbit image analysis software (v3.15). The specificity of the CD4 antibodies were validated on OCT embedded rat spleen sections by immunostaining.

### (ii) Animal

#### SUGEN/Hypoxia Rat Model of Cardiac Fibrosis and Primary Rat Ventricular Cardiac Fibroblasts

This investigation conformed with the Guide for the Care and Use of Laboratory Animals published by the US National Institutes of Health (NIH Publication NO. 85-23, revised 1996) and ARRIVE guidelines. The animal experiments were approved by the Institutional Animal Care and Use Committee (IACUC) of the Providence VA Medical Center. Twenty male Fischer (CDF) rats (F344/DuCrl) weighing 220-250g were obtained from Charles River Laboratories. The animals were divided into two groups: Normoxia (Nx; n=10) and SUGEN/Hypoxia (SuHx; n=10). Rats were administered a single intraperitoneal injection of the VEGFR inhibitor, SUGEN 5416 (Cayman Chemical; 20mg/kg). The animals in the SUGEN/Hypoxia group were then exposed to 3 weeks of normobaric hypoxia (10% FiO_2_, Biospherix, Ltd, Parish, NY), followed by 5 weeks of normoxia (*29*) . The animals in the control group received a single intraperitoneal injection of the vehicle and were exposed to normoxia for the entire duration of the study (8 weeks in total). Fulton’s index, a weight ratio of the RV divided by the sum of left ventricle and septum was measured to determine the extent of right ventricular hypertrophy. Cardiac fibrosis was determined by Sirius red staining and Masson trichrome staining in addition to Eosin and Hematoxylin staining. The images were analyzed using Orbit Image Analysis V3.15 software which can be trained to measure the entire section. Adult rat ventricular fibroblasts were isolated using Langendorff’s free method as previously described (*66*).

#### Echocardiography Measurements

The rats were anesthetized using isoflurane inhalation and subjected to transthoracic echocardiography using Vevo 2100 with a MS250 transducer at 13-24 MHz at baseline and after 3 weeks of hypoxia to measure RV fractional area, RV free wall thickness, pulmonary acceleration time (PAT), pulmonary ejection fraction and tricuspid annular plane systolic excursion via parasternal short-axis view via mid-papillary level. Pulse wave Doppler echo was used to measure pulmonary blood outflow at the levels of aortic valve to measure PAT and PET. TAPSE was measured by aligning an M-mode cursor s to obtain an apical four-chamber view and aligning it as close to the apex of the heart as possible. The remaining parameters, such as Stroke volume (SV), fractional shortening (FS), ejection fraction (EF), and cardiac output (CO) were measured from the left ventricle (LV).The cardiovascular parameters were analyzed on Vevo Lab 3.2.0 using 3 different heart cycles averaged. Rats were euthanized by an overdose of CO_2_, which was continued for one minute after cessation of breathing.

#### Immunohistology of Rat Right Ventricular Tissue

Animals were euthanized and the RV and LV dissected and fixed with 30% formalin. The Optimal Cutting Temperature (OCT) blocks with the samples were sectioned at the mid-ventricular heart region to 5μM thickness by a cryostat before the immunostaining procedure. Samples were then washed in PBS (2x), blocked with 5% normal serum, 1% BSA, and 0.3 M glycine in PBS and then incubated with rabbit monoclonal anti- CD4 and mouse monoclonal anti- αSMA (Table S2) followed by incubation with secondary antibodies. The slides were then washed with PBS (3x), mounted with Vectashield Antifade Mount Medium with DAPI, and visualized using Zeiss LSM 800 Airyscan confocal microscope. Quantification of the CD4 and αSMA regions were performed using orbit image analysis. The individual points represent the average fluorescence intensity of areas captured from 5-10 fields per section for a total of 2 sections. Fibrosis was assessed on 4% paraformaldehyde fixed transverse section of the mid-heart region using standard Masson Trichrome and Sirius Red Staining captured using Aperio Scan Scope. The individual data points represented an average of 10-15 positively stained areas.

### (iii) *in vitro* studies

#### Primary Human Ventricular Cardiac Fibroblast Culture and Stimulation

Human ventricular cardiac fibroblasts (hVCF) were commercially obtained from Lonza (Walkersville, MD) (Cat# CC-2904) with the following lot numbers (Lot# 67771, 62122, 1281202, 534282, TL210281) and a purity of >99%. According to the company records, the cells were isolated from heart explant during transplantation. Cells were obtained and propagated from the ventricle of 3 male and 2 female subjects (n = 5). The donors were between 63-73y of age and were apparently normal. Clinical characteristics of the human subjects are presented in Table S1. The hVCF cells (passage 3-8) were seeded at the density of 2 x 10^5^ cells/ 75cm^2^ flasks in medium supplemented with FGM^TM^-3 Fibroblast Growth Medium-3 BulletKit^TM^ (Cat# CC-4526), 0.1% gentamicin/ amphotericin-B (Gibco, Thermo Fisher Scientific, Waltham, MA, USA) and incubated in a 37°C and 5% CO_2_ incubator.

#### Characterization of the Primary Human Ventricular Cardiac Fibroblast Culture

The hVCF were characterized by morphology, contact inhibition on confluency(*67*), adherence to the culture plate^47^ and immunostaining for collagen Iα, vimentin, fibroblast specific protein (FSP-1), platelet derived growth factor receptor-β (PDGFRβ), α-smooth muscle actin (αSMA) and periostin (Figure S1). The cells were fixed with 4% paraformaldehyde for 10 min at room temperature and washed with PBS. Next, cells were permeabilized with 0.1% Triton-100 and blocked with 5% BSA for 2h at room temperature (*40, 68*). List of antibodies is provided in Table S2. The primary antibodies were diluted (1:100) in the blocking buffer and incubated with the cells at 4°C overnight. The cells were then washed 3x PBS and incubated with species-specific Alexa Fluor 488 conjugated goat anti-rabbit or Alexa Fluor 594 conjugated goat anti-mouse secondary antibodies (1:300; Invitrogen). The cells were then washed 3x PBS and mounted on a 1.5 mm coverslip with a Prolong™ Gold Antifade Mountant with DAPI.

#### Mass cytometry analysis

hVCF (3 x 10^6^) cells treated with vehicle or IL-1β (10ng/mL, when indicated) were subjected to staining with mass cytometry panel of 26 metal conjugated antibodies allowing identification of myeloid, lymphoid, B cells, natural killer cells and dendritic cells (Table S3). The panel design, cell labelling, data processing and analysis is given below.

(i) **Panel Design** .Our cytometry dataset consisted of 200,000 cells analyzed using 28 validated markers and manually gated cell populations with a total number of 3 million cells. Our panel design was based on the identification of immune functions of cardiac fibroblast cells upon stimulation with IL- 1β. We designed the panel to identify the immune cells surface markers and to classify the unidentified cardiac fibroblast population based on the literature evidence of immune cell lineages. Our second criteria was to use previously validated antibodies to avoid the signal overlap issues. Therefore our panel consisting of antibodies that are previously published by the group at Yale aimed at investigating T-cells, NK cells, B cells, monocytes and dendritic cells(*69*).
(ii) **Cell Labelling.** Cell labelling with validated antibodies and mass cytometry assay were performed as previously described and run on CyTOF 1.0 mass cytometer (Fluidigm San Francisco, CA,USA) (*70*),(*69*). Briefly, the cells were incubated with cisplatin, 25μM, Enzo Life Sciences, Farmingdale, NY. 3 x 10^6^ cells were washed with rinsed with 1x PBS and resuspended in 50ul of 1X PBS and 1% BSA containing metal tagged antibody cocktail for 30min at room temperature (Table). The cells were washed twice with PBS and then fixed with 1.6% paraformaldehyde. Cells were washed again with PBS and 1% BSA and incubated overnight at -20°C. Cells were stained with an iridium DNA intercalator (Fluidigm) for 20min at room temperature. Cells were washed again in a series with PBS and water before re-suspending in 1X EQTM Four Element Calibration Beads (Fluidigm) and collected in a CyTOF 1.0 mass cytometer (Fluidigm). Events were normalized as described previously(*70*).
(iii) **Data processing and analysis.** The fcs files generated by the mass cytometry were pre-gated on viable cisplatin and nucleated cells (iridium+). Single, viable, nucleated cells were selected by gating using CYTOFClean R command on the fcs files generated from the CYTOF. Data were analyzed using Cytobank 7.3.0 (Santa Clara, CA, USA) (*71*). The raw median intensity values corresponding to the expression of the immune lineage markers were transformed to hyperbolic arc sine (arcsinh) with a cofactor of 5 for all the datasets. viSNE, FLOWSOM (*72*) and hierarchical clustering were performed using published algorithms (*71, 73*). We used FlowSOM clustering algorithms to annotate the clusters into six major immune lineages on the basis of known lineage markers: (i) CD4^+^ T cells, (ii) B cells, (iii) NK cells, (iv) dendritic cells, (v) lymphocytes (vi) myeloid cells and (v) unidentified population of cells. We used the transformed median expression of each lineage marker for all the cells in the particular cluster to quantify the protein expression (*70*). The raw median intensity values corresponding to the expression of the immune lineage markers were transformed to hyperbolic arc sine (arcsinh) with a cofactor of 5 for all the datasets. Next, to specifically examine the CD4^+^ T cell population in greater details, we manually gated the highly CD4^+^ expressing T-cell population and produced t-SNE maps for the CD4^+^ T cell population. We used the median expression of each lineage marker for all the cells in the particular cluster(*70*). In addition we performed the uniform manifold approximation and projection (UMAP), a non-linear dimensionality reduction technique based on manifold learning technique that preserves local neighborhood relationships and global structure. The code for running UMAP is uploaded as a separated document.

#### Cardiac Fibroblast Culture in ImmunoCult^TM^ –XF T-Cell Expansion Media

Human left ventricular cardiac fibroblast cells were isolated as described above. Cells were cultured in FGM^TM^-3 Fibroblast Growth Medium-3 BulletKit^TM^ (Cat# CC-4526), 0.1% gentamicin/ amphotericin-B (Gibco, Thermo Fisher Scientific, Waltham, MA, USA) for 96h with or without IL-1β at a concentration of 10ng/mL before replacing the media with ImmunoCult^TM^ –XF T-cell expansion media (Cat # 10981) supplemented with IL-1β or no treatment. ImmunoCult^TM^ Human CD3/CD28 T-Cell Activator was added after 72h of culture in ImmunoCult^TM^ –XF T-cell expansion media. Cells were cultured in T-cell expansion media with T Cell Activator in the presence or absence of IL-1β for 14 days.

#### Morphometry Analysis of Cell Clustering

Morphometry was performed on the cell clusters per bright field image at 100μM magnification in a randomized and blinded manner based on one of three criteria. 1) layering of cells forming a rosette; 2) weaving or lattice pattern; 3) formation of a 3D structure that can be visually identified by its white color and clearly defined boundaries.

#### CD4 Antibody Validation Flow Cytometry

The rat blood (1mL) was provided by the Choudhary group at the Providence VA Medical Center. Blood was aliquoted into FACS tubes at 50uL/tube, and stained with either CD4 PE or CD3 APC or the isotypes diluted in FACS buffer. Samples were vortexed, covered with aluminum foil, and incubated at room temperature on gentle shaking for 45 minutes. BD Pharm Lyse ^TM^ (2-8mL) was then added to each sample to lyse the RBC, vortexed, covered with aluminum foil, and incubated for another 15 minutes on low shaking. Cells were then washed with 2mL of FACS buffer per sample, and centrifuged at 1300 rpm for 10 minutes at room temperature. Supernatant was decanted, and cells were re-suspended in 250-500uL of FACS buffer and covered with aluminum foil until run on BD LSR II flow cytometer.

#### TaqMan Human Extracellular Matrix Array and Inflammation Array

Total RNA was extracted from cardiac fibroblast cells using Qiagen RNA isolation kit and converted to cDNA using high capacity reverse transcriptase (Applied Biosystems, Thermo Fisher Scientific ^TM^, Waltham, MA, USA). Transcripts associated with inflammation and extracellular remodeling were analyzed using the TaqMan Human Inflammation Array 96 well, Applied Biosystems, Thermo Fisher Scientific, Cat #: 4414074, Waltham, MA, USA) following the manufacturer’s instructions.

#### Real Time Quantitative Polymerase Chain Reaction (PCR) Analysis

Based on the TaqMan array, the expression level of the selected inflammation (*GAPDH, IL1R, IL-6, CCR2, CCL2, IL-10, TGF β)* and extracellular matrix genes (*COL-I, COL-III, MMP-9*) were evaluated using quantitative RT-PCR (qPCR) analysis. Total RNA was isolated from cardiac fibroblasts using RNeasy Mini Kit (Qiagen Inc., Valencia, CA). First-strand cDNA was synthesized from total RNA (10ng/μl) using high capacity cDNA kit (Applied Biosystems) according to the manufacturer’s instructions. The SYBR green PCR reactions were performed on the cDNA samples using the SYBR green universal master mix. The fold differences in mRNA expression were calculated by normalization of the cycle threshold [C(t)] value of a target gene to reference gene (β-actin and 18sRNA) using the Livak method. The primers for genes are provided in Table S5.

#### Transmission Electron Microscopy

The hVCF cells seeded on glass coverslips were washed 3x and fixed in 1% glutaraldehyde in 0.05M cacodylate buffer. Samples were then post-fixed in 1% osmium tetroxide and 1% uranyl acetate. Samples were dehydrated, embedded, and examined using Philips 410 Transmission Electron Microscope.

#### Immunoblotting

Immunoblotting procedure was performed on primary hVCF cells as previously described by the group (*74*). Briefly, 1 x10^6^ cells were collected from petriplates and lysed in radio immunoprecipitation assay buffer containing phosphatase inhibitor cocktail. The tubes were spun at 5,000g for 5 min and supernatant transferred to new tubes. The protein content in the lysate were measured using BCA protein assay and normalized to yield 100 μg/mL. 1x Laemelli sample buffer (containing 0.1% β-mercaptoethanol) was added and the samples heated for 10 min at 90°C. Protein samples were separated based on molecular weight by SDS-PAGE and transferred to polyvinylidene fluoride membranes followed by blocking in 5% Milk or 5% BSA for 1 h. The membranes were incubated with primary antibodies (Table S2) in blocking buffer overnight. The membranes were then washed 3 times with TBST and incubated with secondary antibodies for 2 h in blocking buffer. The phosphorylated proteins were detected by primary antibodies, which were recognized by horseradish peroxidase–conjugated or fluorophore (IRDye 800CW and IRDye 680RD)-conjugated, species-specific secondary antibodies. The fluorescence signals were captured and quantified using LiCOR Odyssey Imaging Systems.

#### Sample Preparation for Proteomic Analysis

hCVF (2 x 10^6^) were seeded and grown to 80% confluence and treated with IL-1β (10ng/ml) for 24h. Cell culture media were then collected and subjected to proteomic analysis. Conditioned media were concentrated using ProteoSpin TM column (Norgen Biotek Corp, Canada). Concentrated samples were lysed with buffer (8 M urea, 1 mM sodium orthovanadate, 20 mM HEPES, 2.5mM sodium pyrophosphate, 1 mM β-glycerophosphate, pH 8.0), followed by sonication (QSonica, LLC, Model no. Q55), and cleared by centrifugation (*75*). The LC-MS/MS was performed on a fully automated proteomic technology platform that includes an Agilent 1200 Series Quaternary HPLC system (Agilent Technologies, Santa Clara, CA) connected to a Q Exactive Plus mass spectrometer (Thermo Fisher Scientific, Waltham, MA). The MS/MS spectra were acquired at a resolution of 17,500, with a targeted value of 2 ×10^4^ ions or maximum integration time of 200 ms. The ion selection abundance threshold was set at 8.0 ×10^2^ with charge state exclusion of unassigned and z =1, or 6-8 ions and dynamic exclusion time of 30 seconds. Peptide spectrum matching of MS/MS spectra of each file was searched against the human database (UniProt) using the Sequest algorithm within Proteome Discoverer v 2.3 software (Thermo Fisher Scientific, San Jose, CA).

#### Cytokine and Chemokine Detection

Cytokine and chemokine assays were performed by the Forsyth Multiplex Core (Cambridge, MA, USA) using the Human Cytokine/Chemokine Magnetic Bead Panel (Milliplex, Millipore Sigma, Burlington, MA, USA) and Bio-Plex®200 plate reader following manufacturers’ specifications. Data were analyzed using Bio-Plex Manager software v6.0.

#### Metabolomics

Metabolomics of conditioned media was performed at the Beth Israel Deaconess Medical Center Mass Spectrometry Core (*75*). hVCF (2 x 10^6^) were seeded and grown to confluence and treated with IL-1β (10ng/ml) for 24h. Conditioned media (1ml) from the vehicle or IL-1β treated hVCF cells were used for the metabolomics analysis. 500μL of chilled -80°C 80% methanol was added to 15ml tubes containing 1ml of conditioned media and evaporated using Speedvac to pellet the metabolites. SRM with polarity switching with a QTRAP 5500 mass spectrometer (AB/SCIEX) was used to assay 300 polar molecules, based on the previously published protocol(*75*).

#### Illumina RNA Seq Analysis

hVCF Total RNA was extracted from flow sorted Veh CD4^+^ Veh CD4^-^ cells and IL-1β CD4^+^ and IL1β CD4^-^ cells using Qiagen RNeasy Mini Kit (Cat# 74134) and the samples submitted to Genewiz for RNA Seq analysis. Initial sample QC analysis was performed followed by RNA library preparation with polyA selection and sequenced on illumina Hi Seq in 2 x 150 bp paired end configuration. The raw data was obtained in FASTQ format and sequence reads were trimmed to remove possible adapter sequences and poor quality nucleotides using Trimmomatic v.0.36. The trimmed reads were mapped to Homo sapiens GRCh38 reference genome available on ENSEMBL using the STAR aligner v.2.5.2b. The unique gene hit counts were calculated using feature counts from Subread package v.1.5.2. And the gene hit counts were used for differential expression analysis. The gene expressions were compared using DESeq2 and Wald test was used to generate the p values and log2 fold changes. The genes with an adjusted p-value <0.05 and absolute log2 fold change >1 were considered as differentially expressed gene. Gene ontology on unique set of genes for each population was performed using Database for Annotation, Visualization and Integrated Discovery (DAVID) (https://david.ncifcrf.gov/) v6.8.

### Statistical Analysis

The normal or Gaussian distribution for small sample sizes was determined using Shapiro-Wilk’s test. For statistical comparisons between 2 groups, 2-tailed unpaired *t* test was used for unequal variances. When the sample sizes were small and not normally distributed, non-parametric tests 2-tailed Mann-Whitney U test was performed. Comparisons of >2 groups were made with 1- or 2- way ANOVA with either Tukey’s post-hoc test as stated in the figure legends. Kruskal-Wallis tests followed by Dunn test comparison was used for continuous variables that did not show normal distribution. The values \**P*<0.01and \**P*<0.05 was considered significant. The analyses were performed in GraphPad Prism version 8.0. Missing data are common in both proteomics and metabolomics due to heterogeneous responses to treatment or if the abundances of the protein or metabolite is below the detection limit of the mass spectrometer or a reduction in the electrospray performance. We used the Xia et al approach to disregard the variables containing missing data for our statistical analysis and focusing on the metabolites where the information was present for all the three human donors in both vehicle and IL-1β groups (*76*). In some cases where there was a single missing value in one of the replicates, imputation using mean values were performed to replace the missing values. To control for multiplicity of the data, we employed the false discovery rate or Bonferroni correction as indicated.

## Author contributions

Conceptualization: JHS, PD, RJG; Methodology: JHS, PD, FSP, AZ, HG, SC; Investigation: JHS, FSP, AZ, HG, SC; Visualization: JHS, RJG, PD and SR; Funding acquisition: JHS, SR; Project administration: JHS; Supervision: JHS, SR, RJG; Writing (original draft): JHS, RJG; Writing (review & editing): JHS, RJG, PD, SR, SS.

## Competing interest statement

Dr. Sadayappan provided consulting and collaborative research studies to the Leducq Foundation (CURE-PLAN), Red Saree Inc., Greater Cincinnati Tamil Sangam, AstraZeneca, MyoKardia, Merck and Amgen. No other competing interests are reported.

## Acknowledgments

We thank Dr. Ruth Montgomery, PhD, and Ms. Shelly Ren from the Yale CyTOF core facility and Dr. John M Asara, PhD, Director of the Mass Spectrometry core, Harvard Medical School, for helpful discussions. We thank Siraj Presswala for his assistance with the UMAP analysis using PYTHON and R tools and Daniel Federick from Fluidigm for help with the tSNE analysis using CYTOBANK. We also acknowledge Nagib Ahsan and Lelia Noble from the COBRE Center for Cancer Research Development, Proteomics Core Facility, Rhode Island Hospital, Providence, RI, for their help with proteomic sample processing and analysis. The authors also acknowledge support from Dr. Fenghai Duan of advance CTR (U54GM115677) for help with the statistical analysis of proteomics and metabolomics data. We thank Yang Zhou for his help with the Fulton index animal measurements. This work was performed using the facilities and resources of CPVB COBRE and Providence Veterans Affairs Medical Center. The study was funded in part by grant from the National Institutes of Health P20 GM103652 Pilot Award (JHS), National Institutes of Health R01HL130230 (SR), Advance CTR (U54GM115677), R56/R01 HL139680 (RJG, SS), R01 HL130356 (SS), R01 HL105826 (SS), R01 AR078001 (SS) and R01 HL143490 (SS), Rhode Island Foundation Grant (20190594) (JHS) and TEAM UTRA grant from Brown University (JHS).

## Data and materials availability

All data are available in the main text or the supplementary materials. The raw data files for mass cytometry will be available on FlowRepository. RNA-seq data will be deposited in gene expression omnibus and proteomics data will be deposited in PRIDE and metabolomics data will be deposited in Metabolomics workbench.

## Notes

### Competing Interest Statement

The authors have declared no competing interest.

### Summary of Updates

The revisions include presenting the figures more clearly and improving the manuscript text.

