## Supplementary figure files for "Identification of CD4^+^ Sub-population of Resident Cardiac Fibroblasts Linked to Myocardial Fibrosis"

### Figure S1

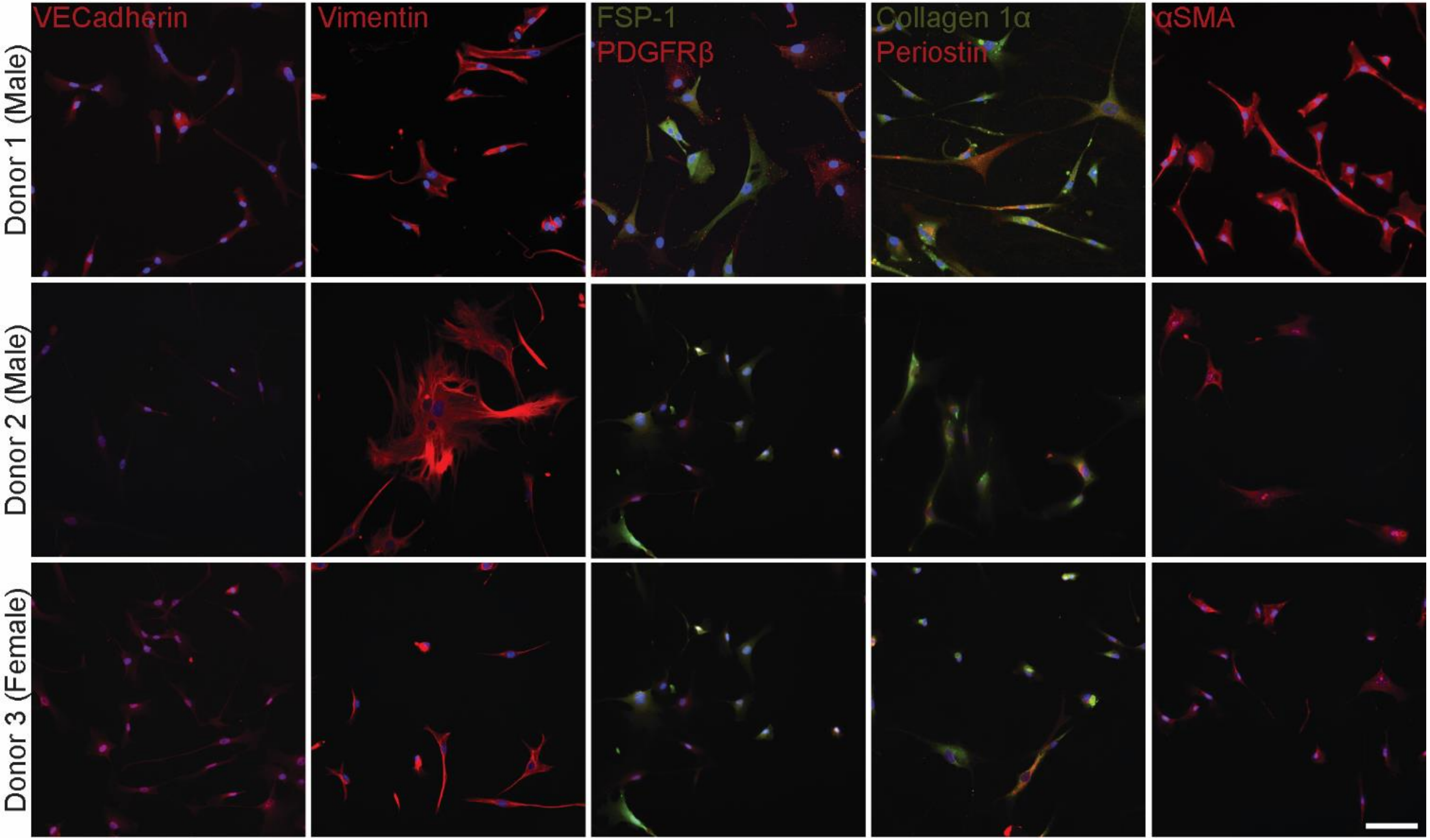
VECadherin
Vimentin
FSP-1
PDGFRβ
Collagen 1α
Periostin
αSMA
Donor 1 (Male)
Donor 2 (Male)
Donor 3 (Female)

### Figure S2

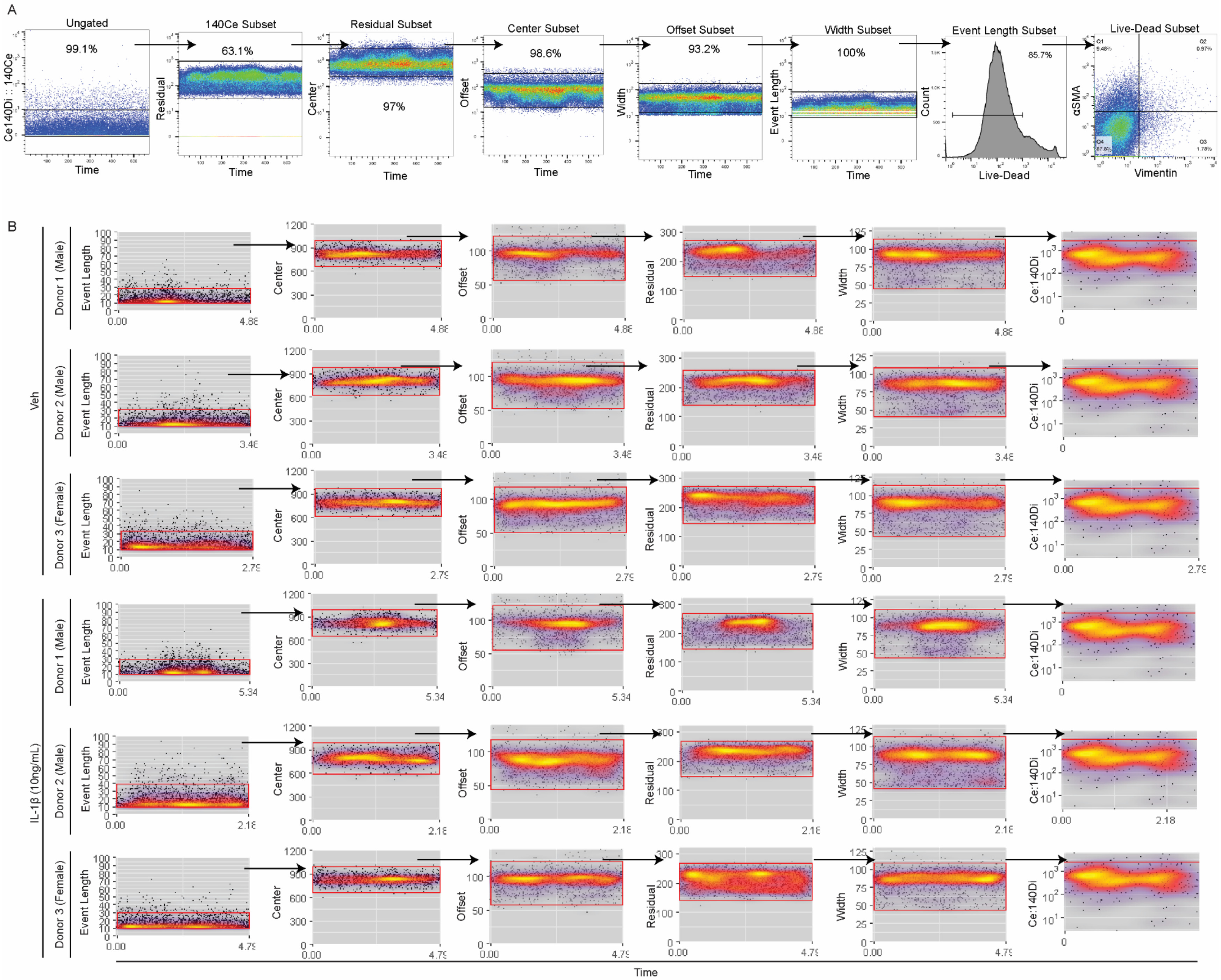
A
Ungated
99.1%
Ce140Di :: 140Ce
Time
140Ce Subset
63.1%
Residual
Residual Subset
Center
97%
Center Subset
98.6%
Offset
Offset Subset
93.2%
Width
Width Subset
100%
Event Length
Event Length Subset
85.7%
Count
Live-Dead
Live-Dead Subset
αSMA
Vimentin
B
Veh
IL-1β (10ng/mL)
Donor 1 (Male)
Donor 2 (Male)
Donor 3 (Female)
Event Length
Center
Offset
Residual
Width
Ce:140Di
Time

### Figure S3

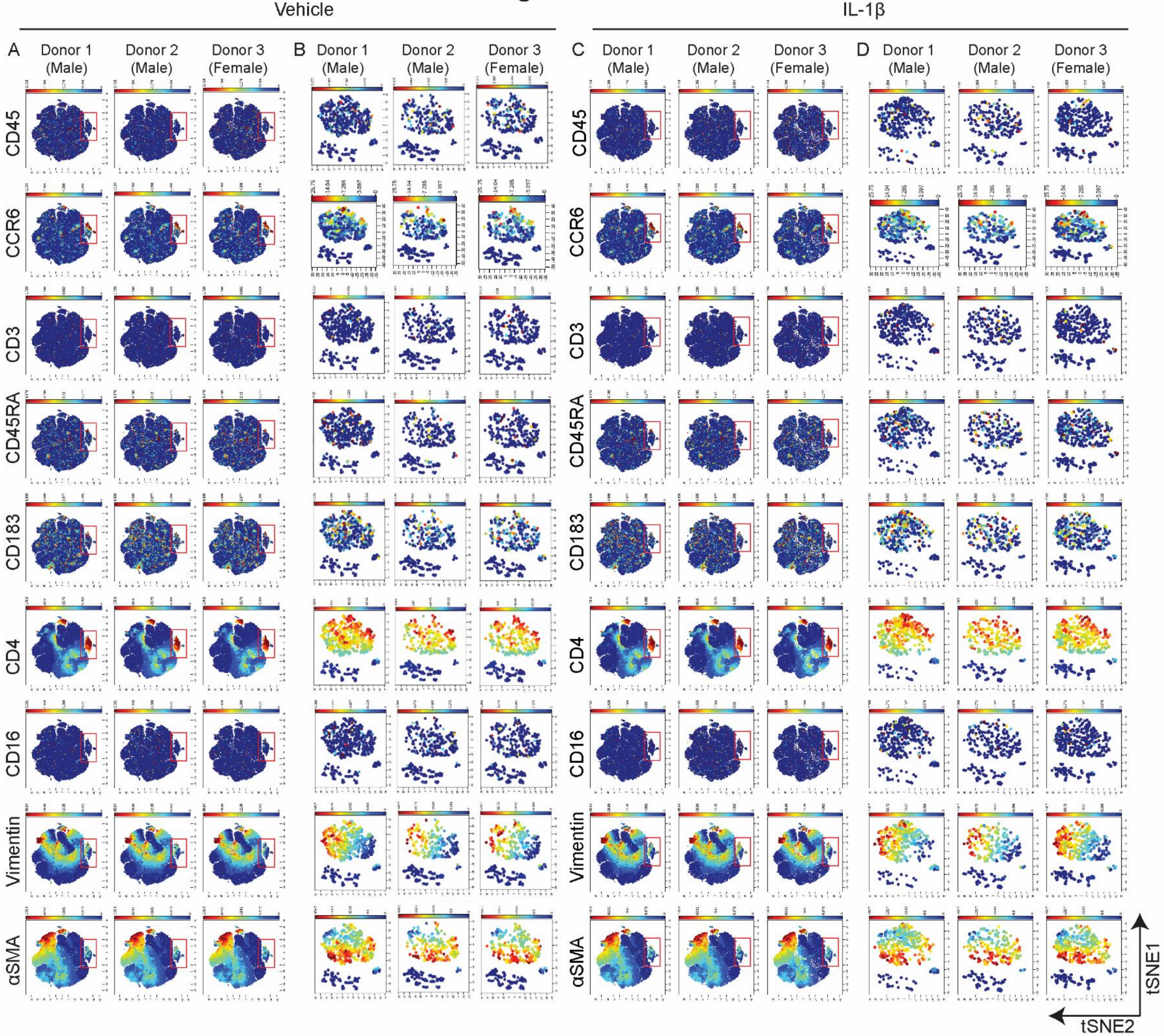

Vehicle
IL-1β
A
B
C
D
Donor 1 (Male)
Donor 2 (Male)
Donor 3 (Female)
CD45
CCR6
CD3
CD45RA
CD183
CD4
CD16
Vimentin
αSMA
tSNE1
tSNE2

### Figure S4

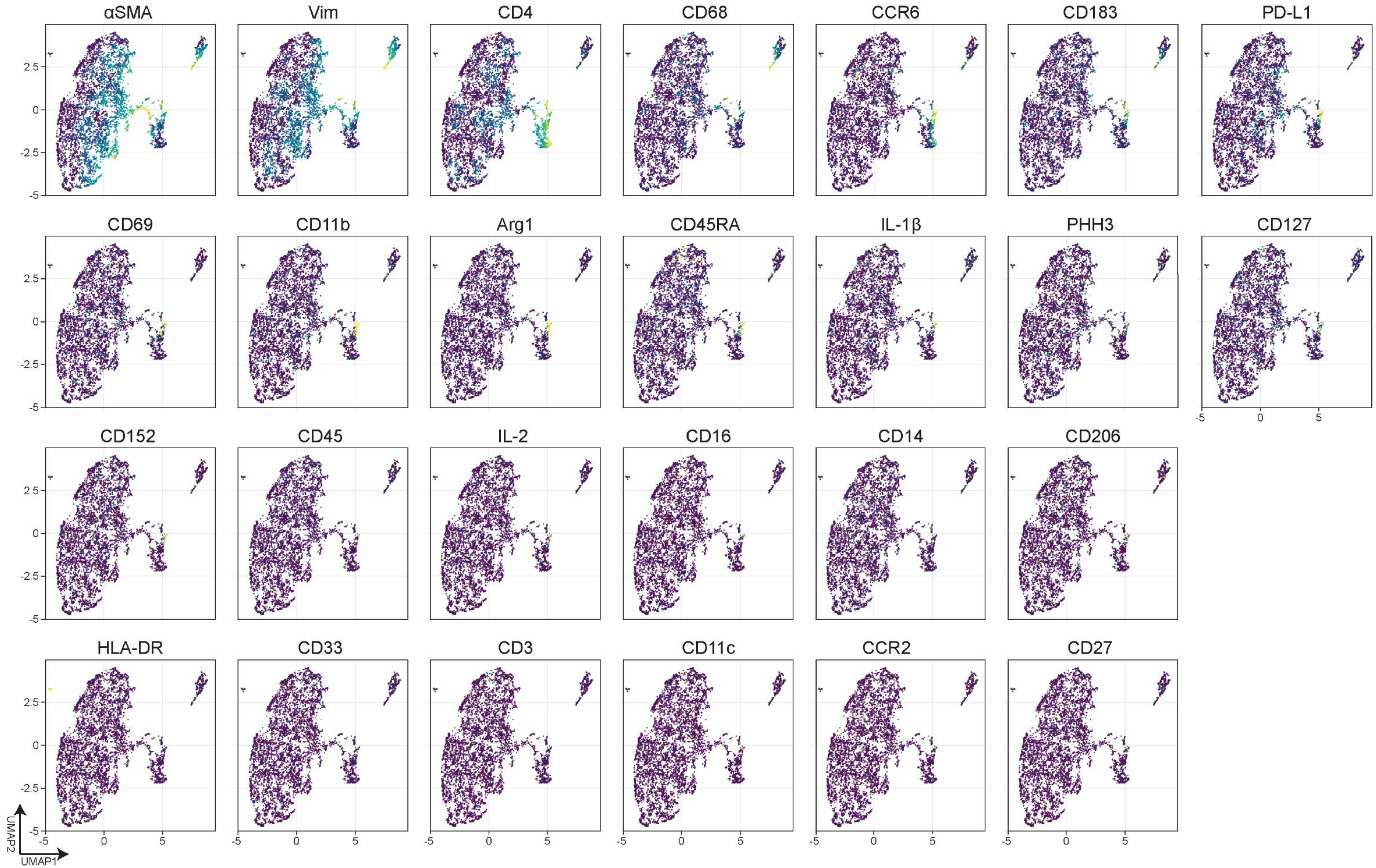
αSMA
Vim
CD4
CD68
CCR6
CD183
PD-L1
CD69
CD11b
Arg1
CD45RA
IL-1β
PHH3
CD127
CD152
CD45
IL-2
CD16
CD14
CD206
HLA-DR
CD33
CD3
CD11c
CCR2
CD27
2.5
0
-2.5
-5
0
5
UMAP2
UMAP1

### Figure S5

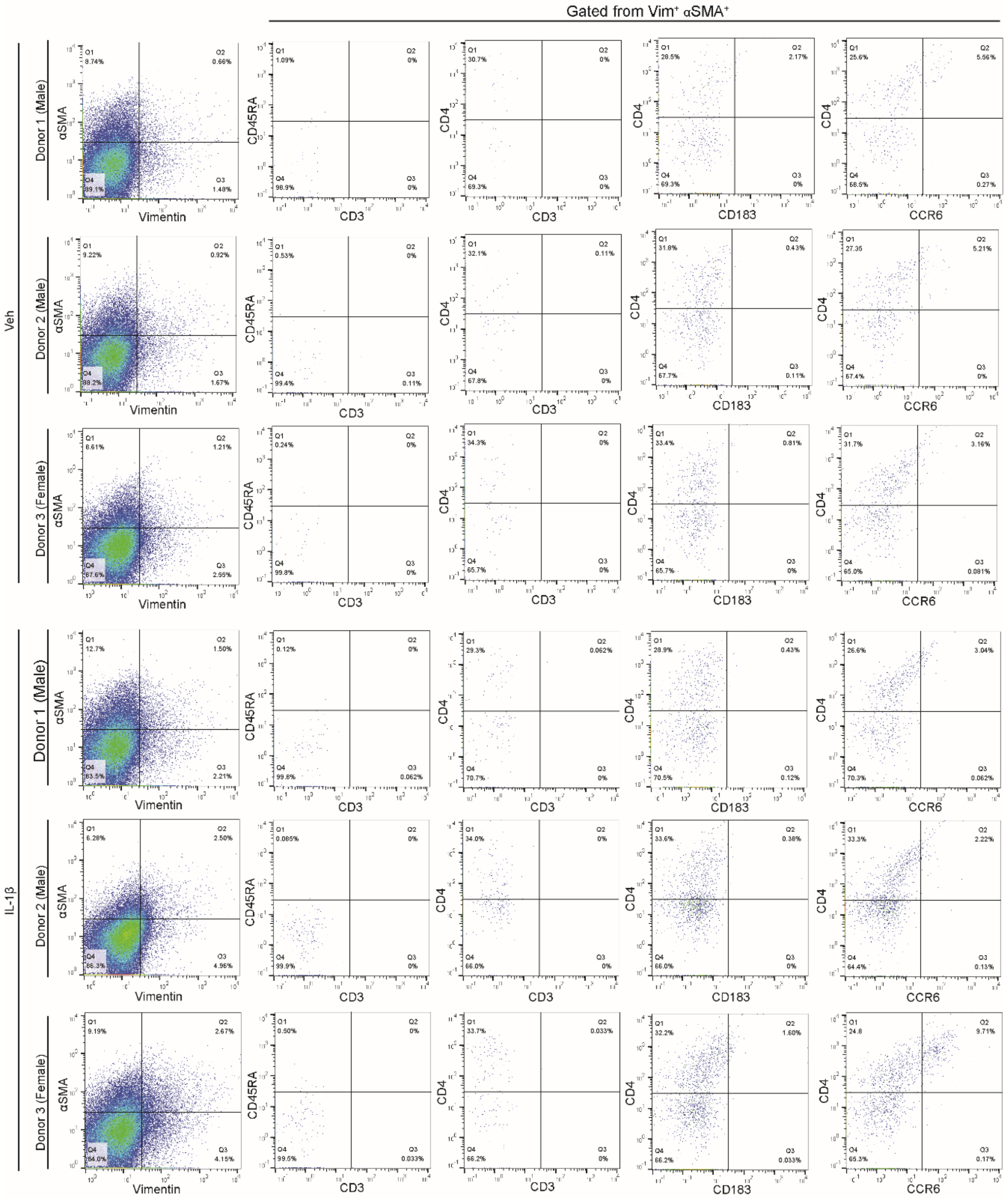
Gated from Vim⁺ αSMA⁺
Veh
IL-1β
Donor 1 (Male)
Donor 2 (Male)
Donor 3 (Female)
αSMA
Vimentin
CD45RA
CD3
CD4
CD183
CCR6

### Figure S6

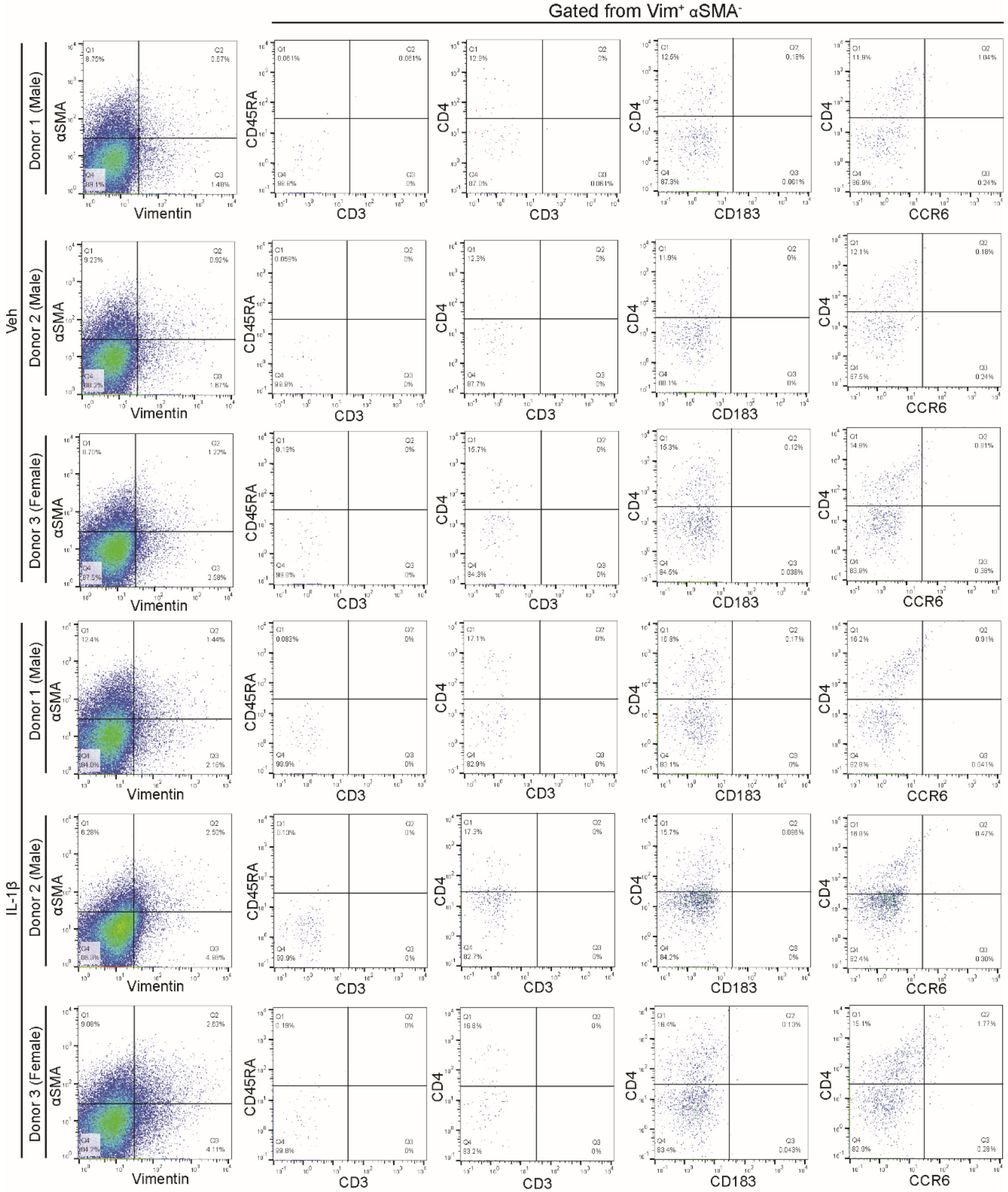
Gated from Vim⁺ αSMA⁻
Veh
IL-1β
Donor 1 (Male)
Donor 2 (Male)
Donor 3 (Female)
αSMA
Vimentin
CD45RA
CD3
CD4
CD183
CCR6

### Figure S7

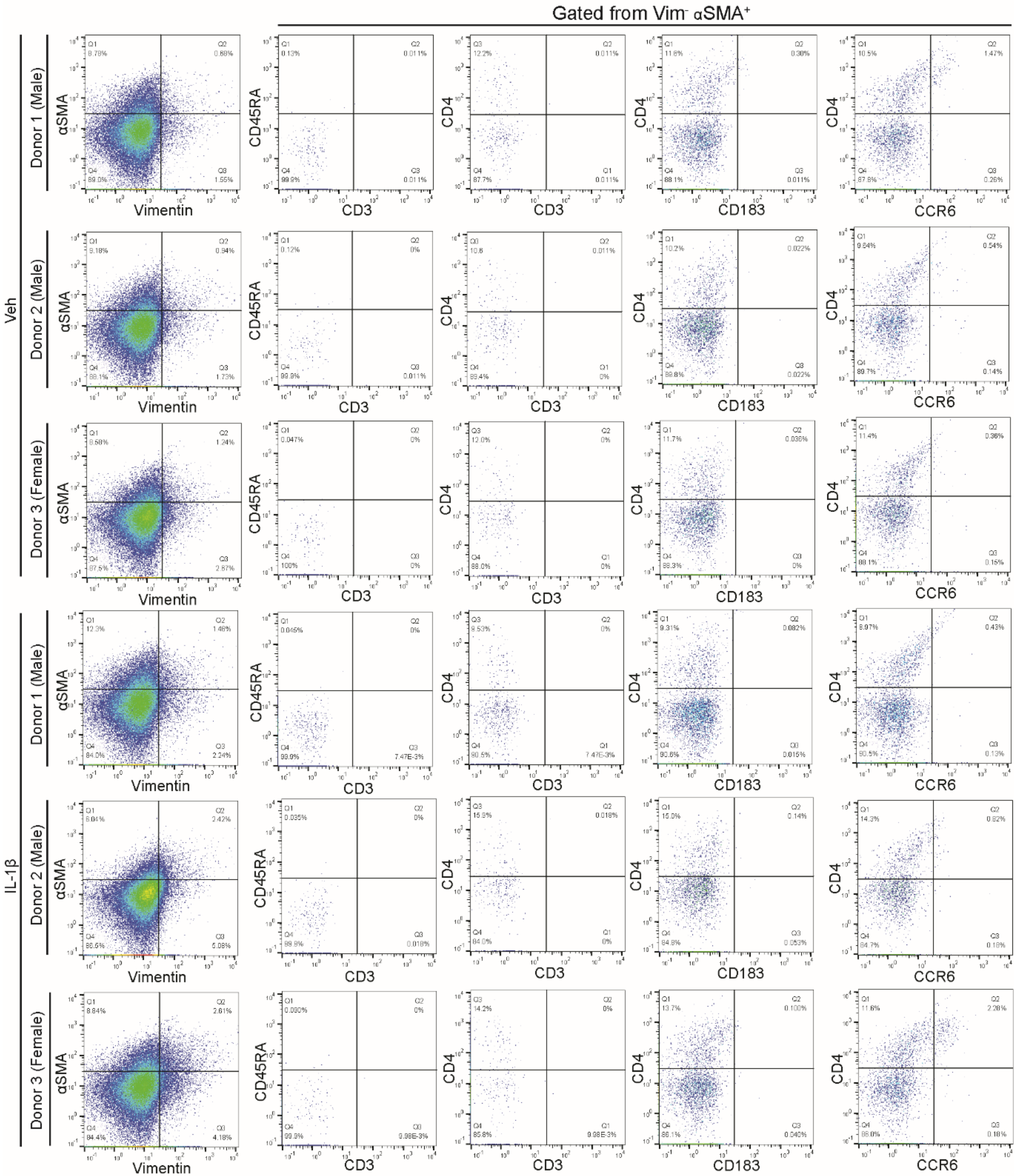

Gated from Vim⁻ αSMA⁺
Veh
IL-1β
Donor 1 (Male)
Donor 2 (Male)
Donor 3 (Female)
αSMA
Vimentin
CD45RA
CD3
CD4
CD183
CCR6

### Figure S8

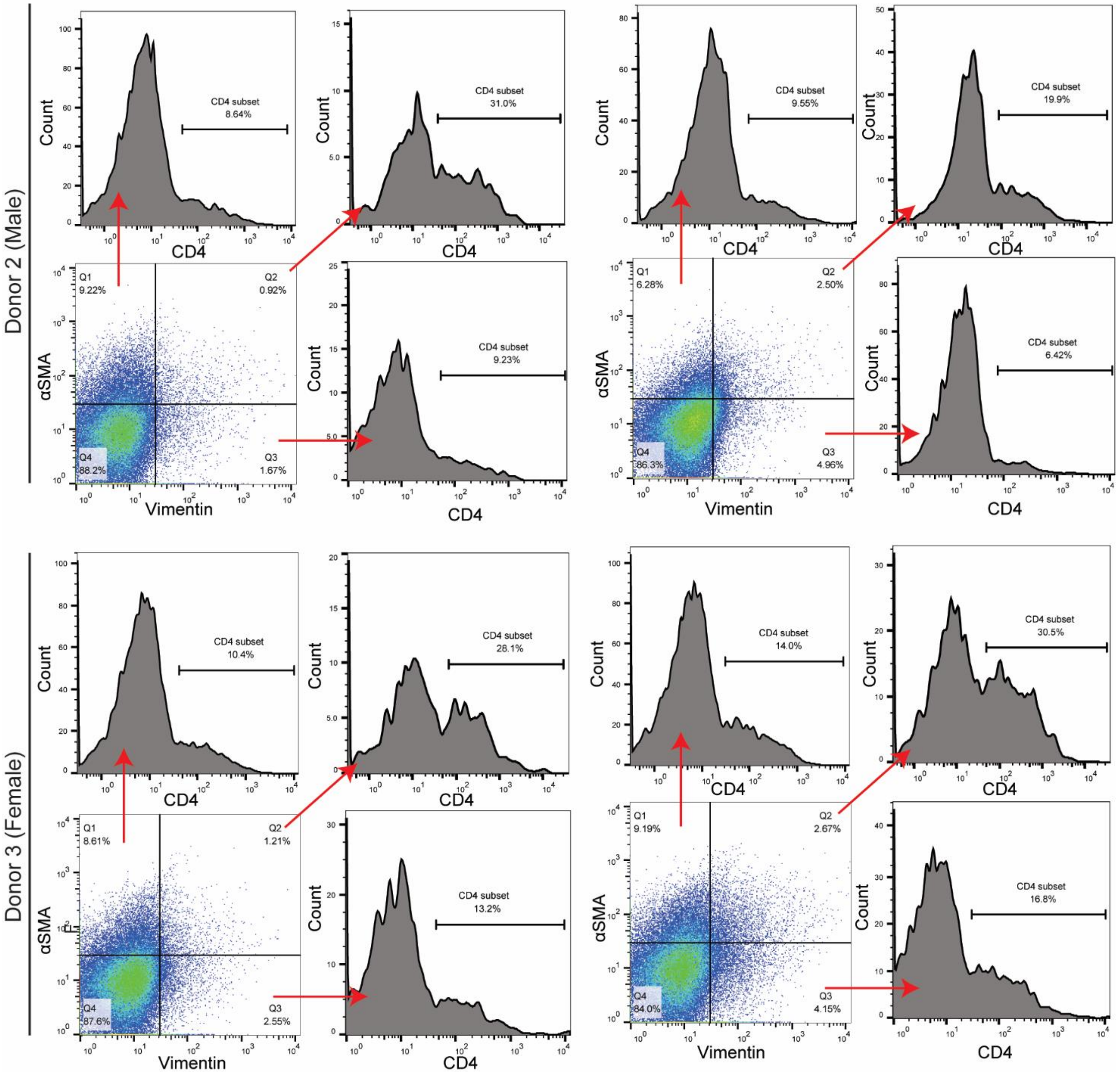
Donor 2 (Male)
Donor 3 (Female)
Count
CD4
αSMA
Vimentin
CD4 subset
8.64%
31.0%
9.23%
9.55%
19.9%
6.42%
10.4%
28.1%
13.2%
14.0%
30.5%
16.8%
Q1
9.22%
Q2
0.92%
Q3
1.67%
Q4
88.2%
Q1
6.28%
Q2
2.50%
Q3
4.96%
Q4
86.3%
Q1
8.61%
Q2
1.21%
Q3
2.55%
Q4
87.6%
Q1
9.19%
Q2
2.67%
Q3
4.15%
Q4
84.0%

### Figure S9

A

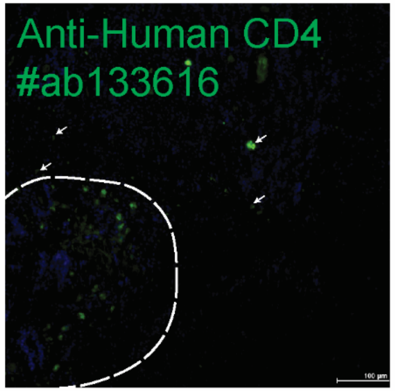
Anti-Human CD4
#ab133616

B

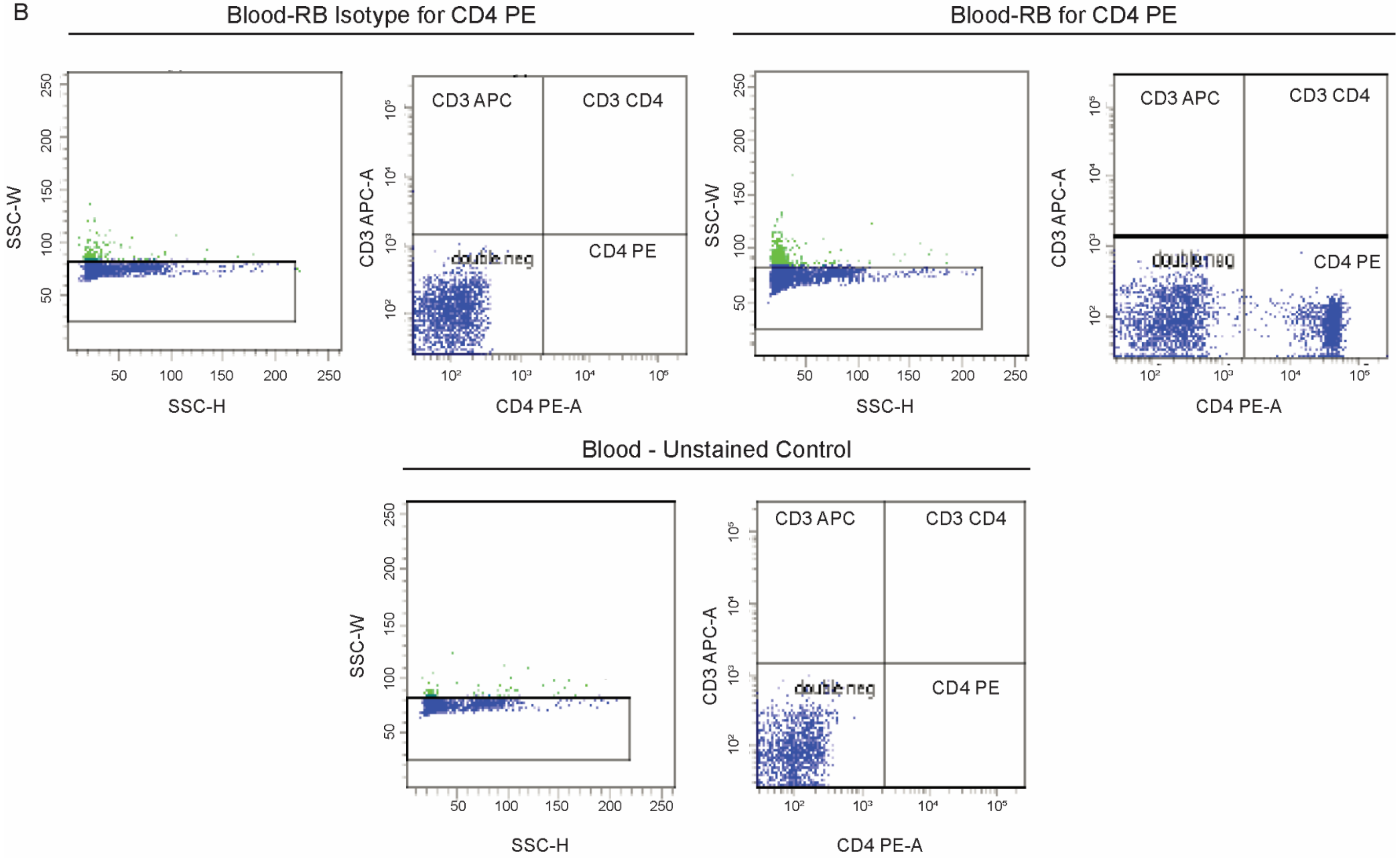
Blood-RB Isotype for CD4 PE
Blood-RB for CD4 PE
Blood - Unstained Control
SSC-W
SSC-H
CD3 APC-A
CD4 PE-A
CD3 APC
CD3 CD4
double neg
CD4 PE

### Figure S10

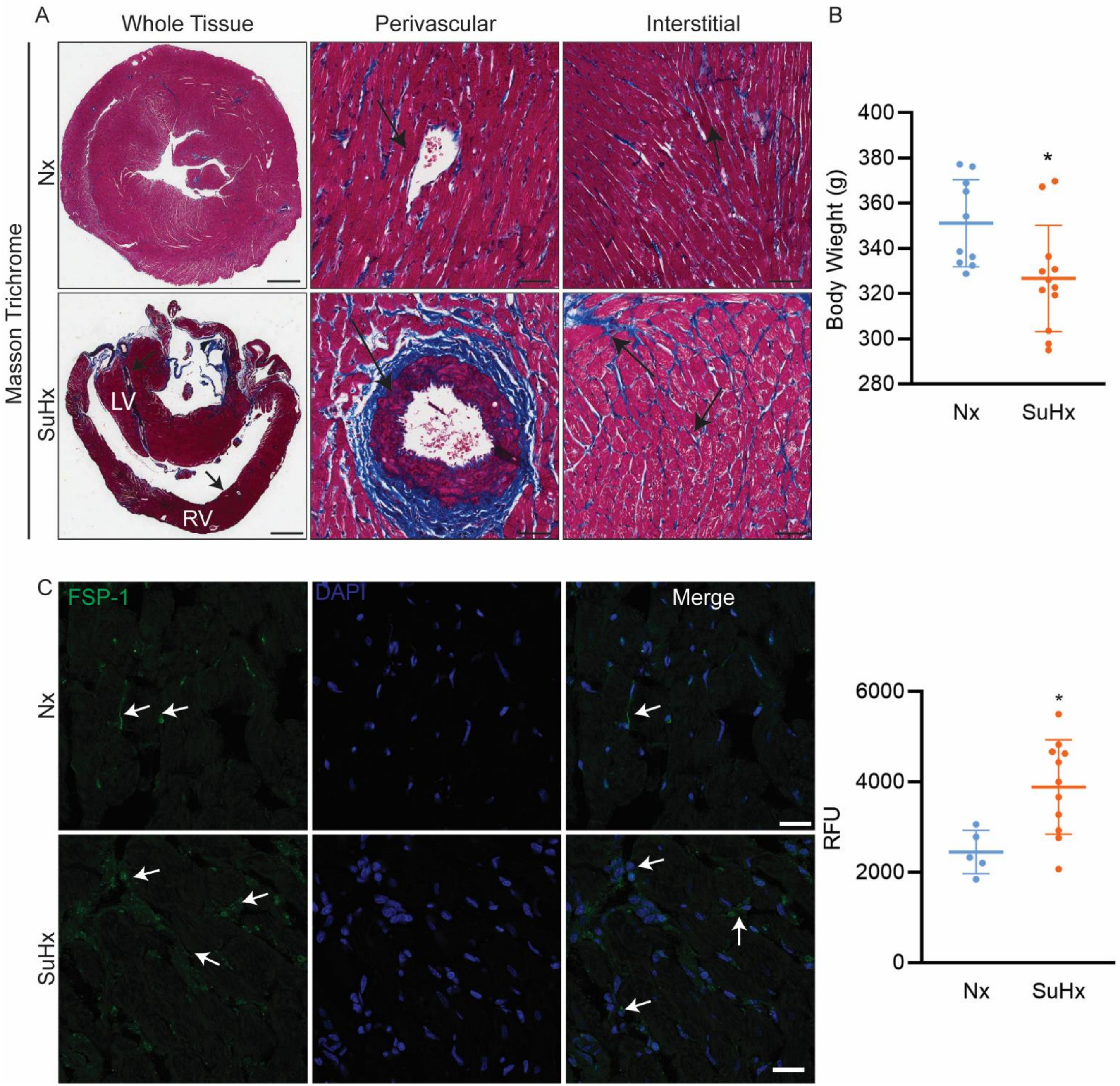
A
Whole Tissue
Perivascular
Interstitial
Masson Trichrome
Nx
SuHx
LV
RV
B
Body Wieght (g)
400
380
360
340
320
300
280
*
Nx
SuHx
C
FSP-1
DAPI
Merge
Nx
SuHx
RFU
6000
4000
2000
0
*
Nx
SuHx

### Figure S11

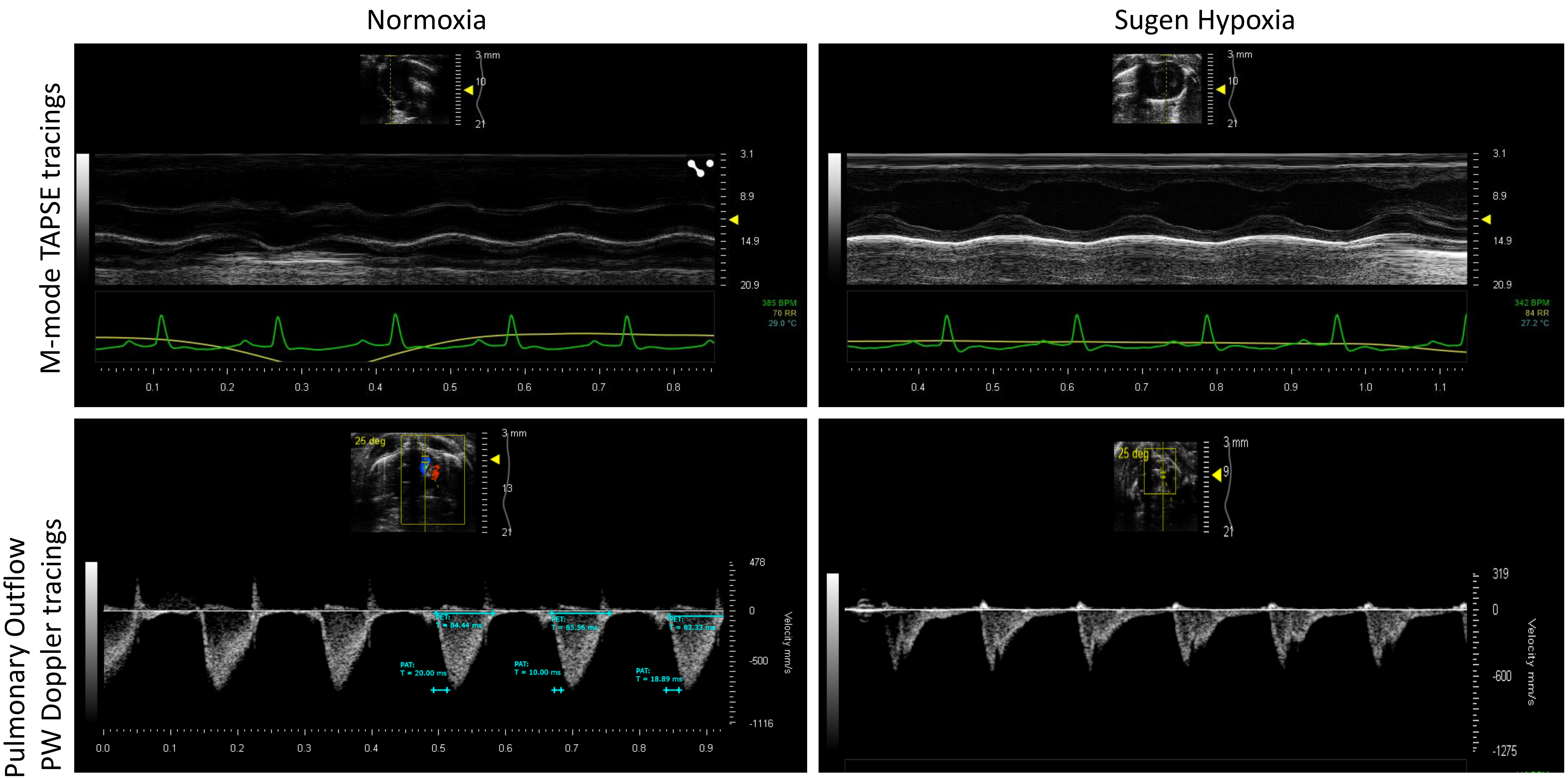
Normoxia
Sugen Hypoxia
M-mode TAPSE tracings
Pulmonary Outflow
PW Doppler tracings
385 BPM
70 RR
29.0 °C
342 BPM
84 RR
27.2 °C
PET:
T = 84.44 ms
PET:
T = 85.56 ms
PET:
T = 83.33 ms
PAT:
T = 20.00 ms
PAT:
T = 10.00 ms
PAT:
T = 18.89 ms
Velocity mm/s
25 deg

### Figure S12

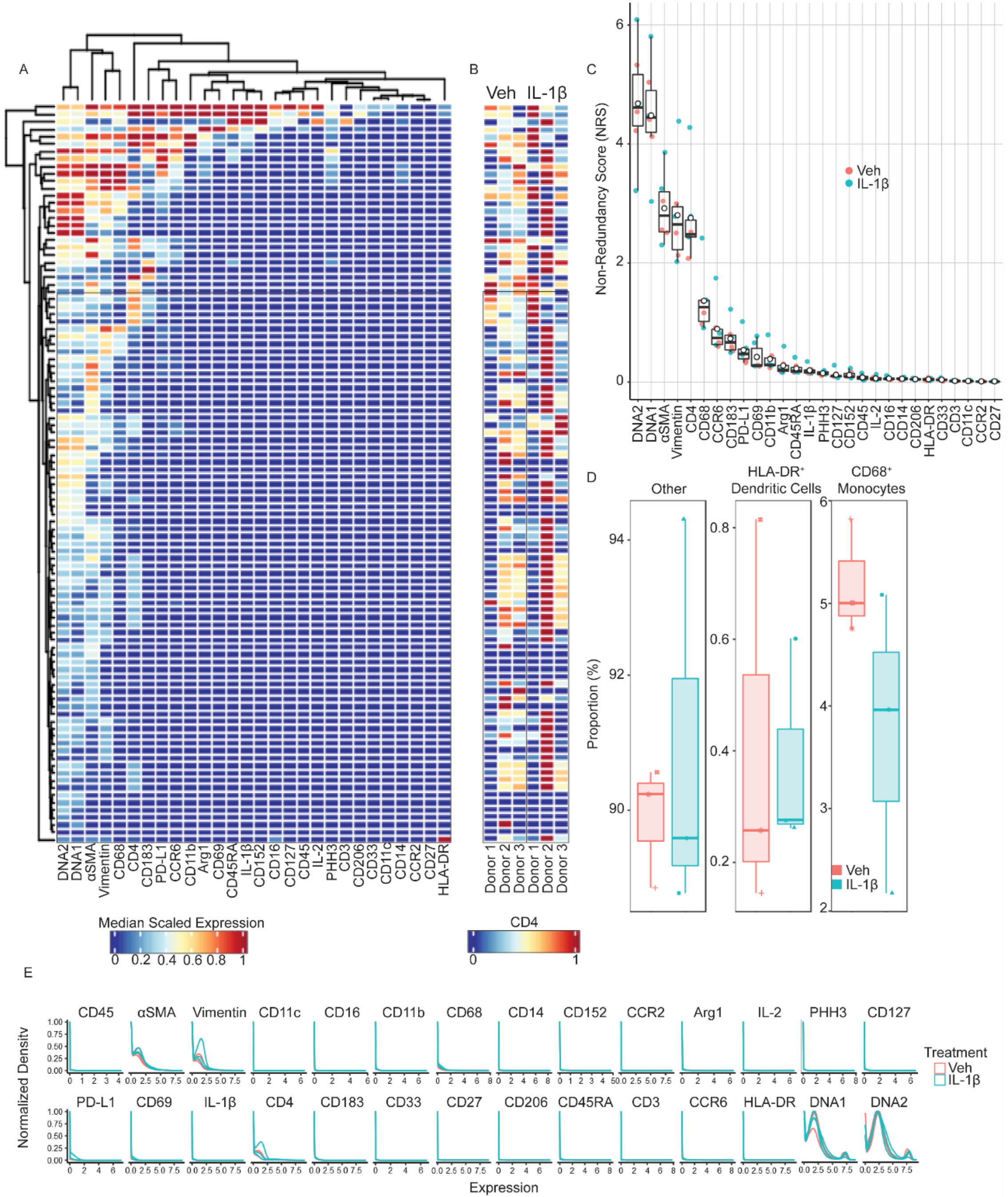

A
B
Veh IL-1β
C
Non-Redundancy Score (NRS)
Veh
IL-1β
D
Other
HLA-DR⁺ Dendritic Cells
CD68⁺ Monocytes
Proportion (%)
Median Scaled Expression
CD4
E
Normalized Density
Expression
Treatment
Veh
IL-1β

Figure S13

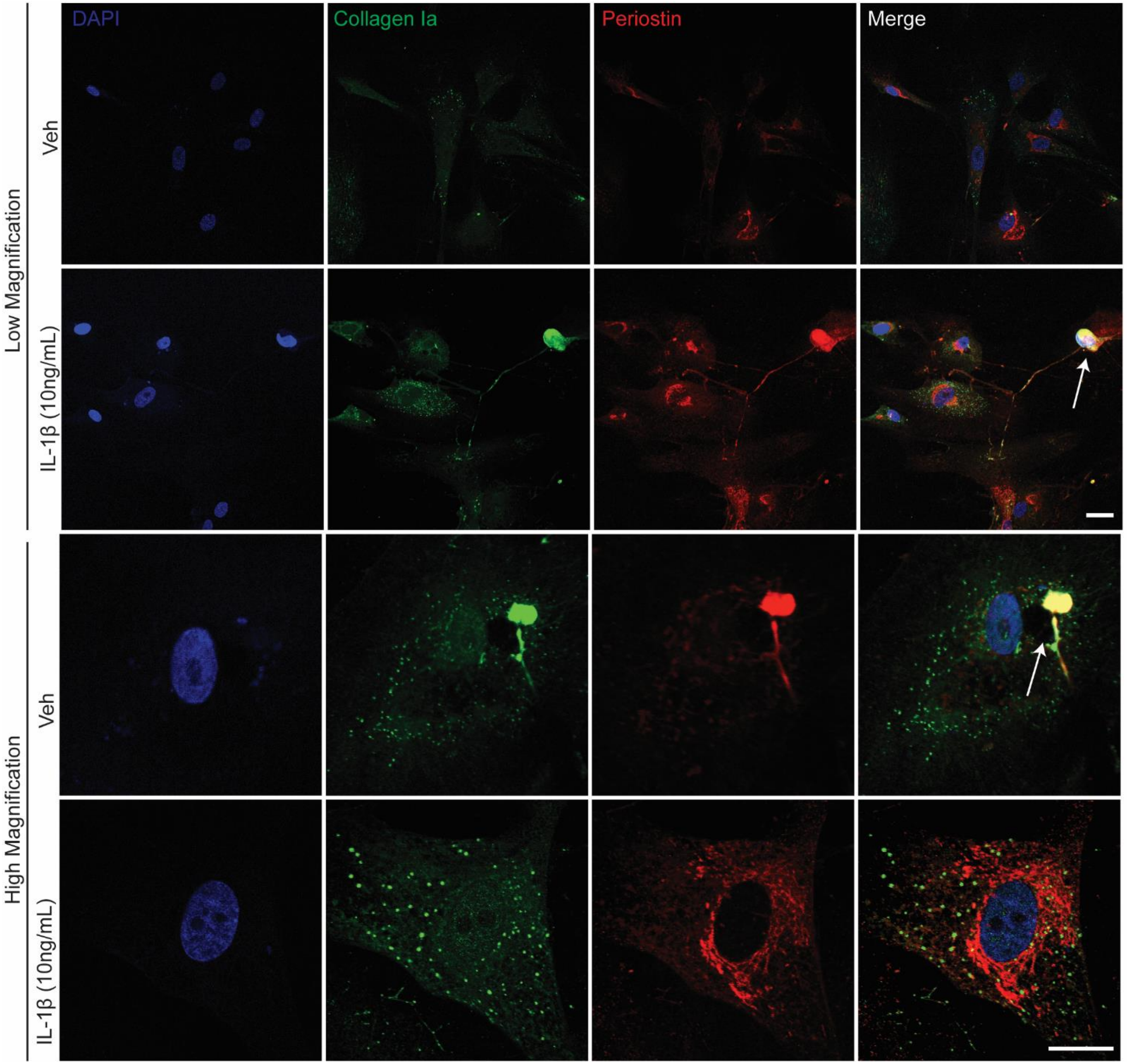
DAPI
Collagen Ia
Periostin
Merge
Low Magnification
Veh
IL-1β (10ng/mL)
High Magnification
Veh
IL-1β (10ng/mL)

### Figure S14

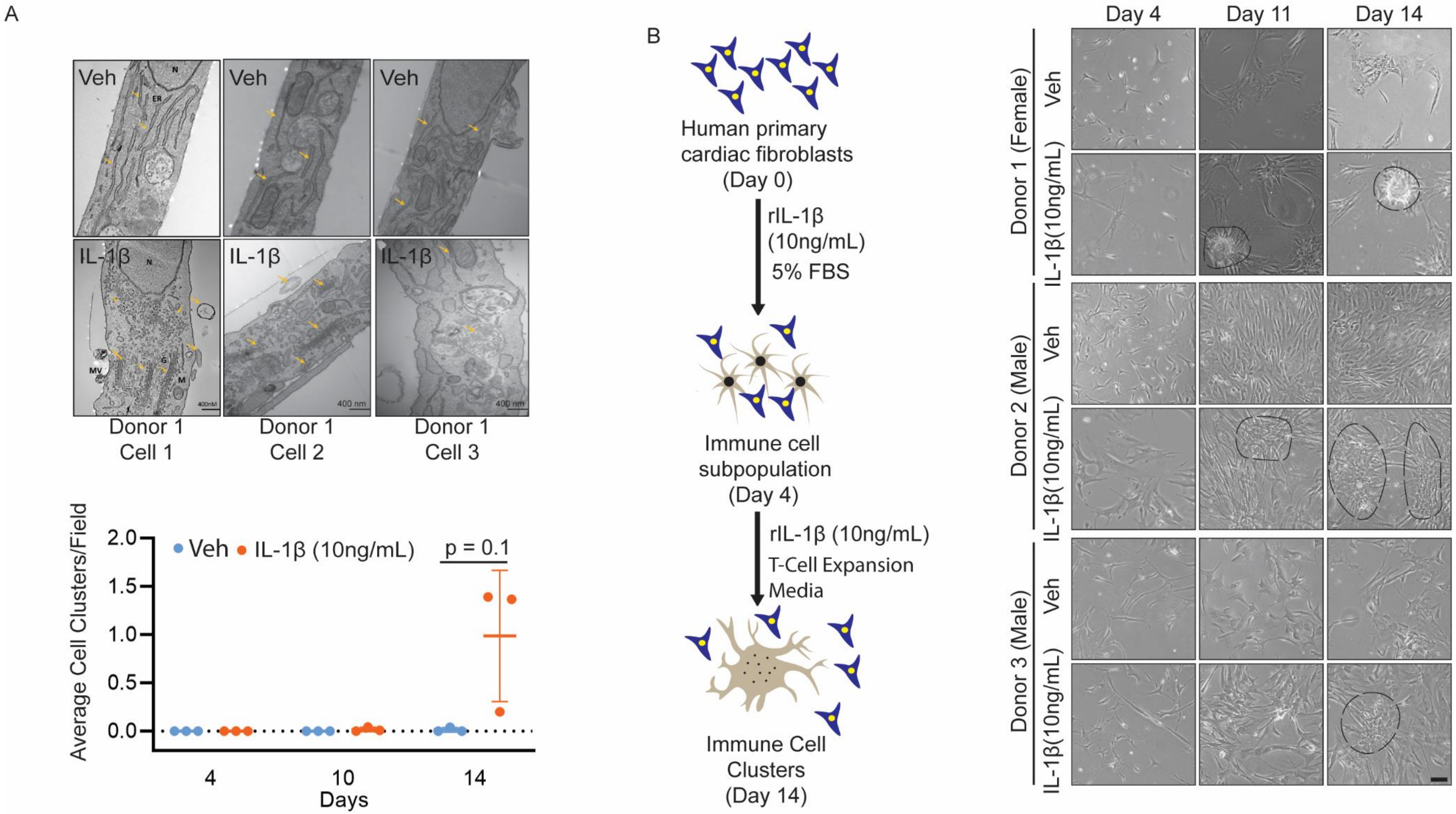

A
Veh
Veh
Veh
IL-1β
IL-1β
IL-1β
Donor 1
Cell 1
Donor 1
Cell 2
Donor 1
Cell 3
Average Cell Clusters/Field
2.0
1.5
1.0
0.5
0.0
Veh
IL-1β (10ng/mL)
p = 0.1
4
10
14
Days
B
Human primary cardiac fibroblasts (Day 0)
rIL-1β (10ng/mL)
5% FBS
Immune cell subpopulation (Day 4)
rIL-1β (10ng/mL)
T-Cell Expansion Media
Immune Cell Clusters (Day 14)
Day 4
Day 11
Day 14
Donor 1 (Female)
Donor 2 (Male)
Donor 3 (Male)
Veh
IL-1β(10ng/mL)

### Figure S15

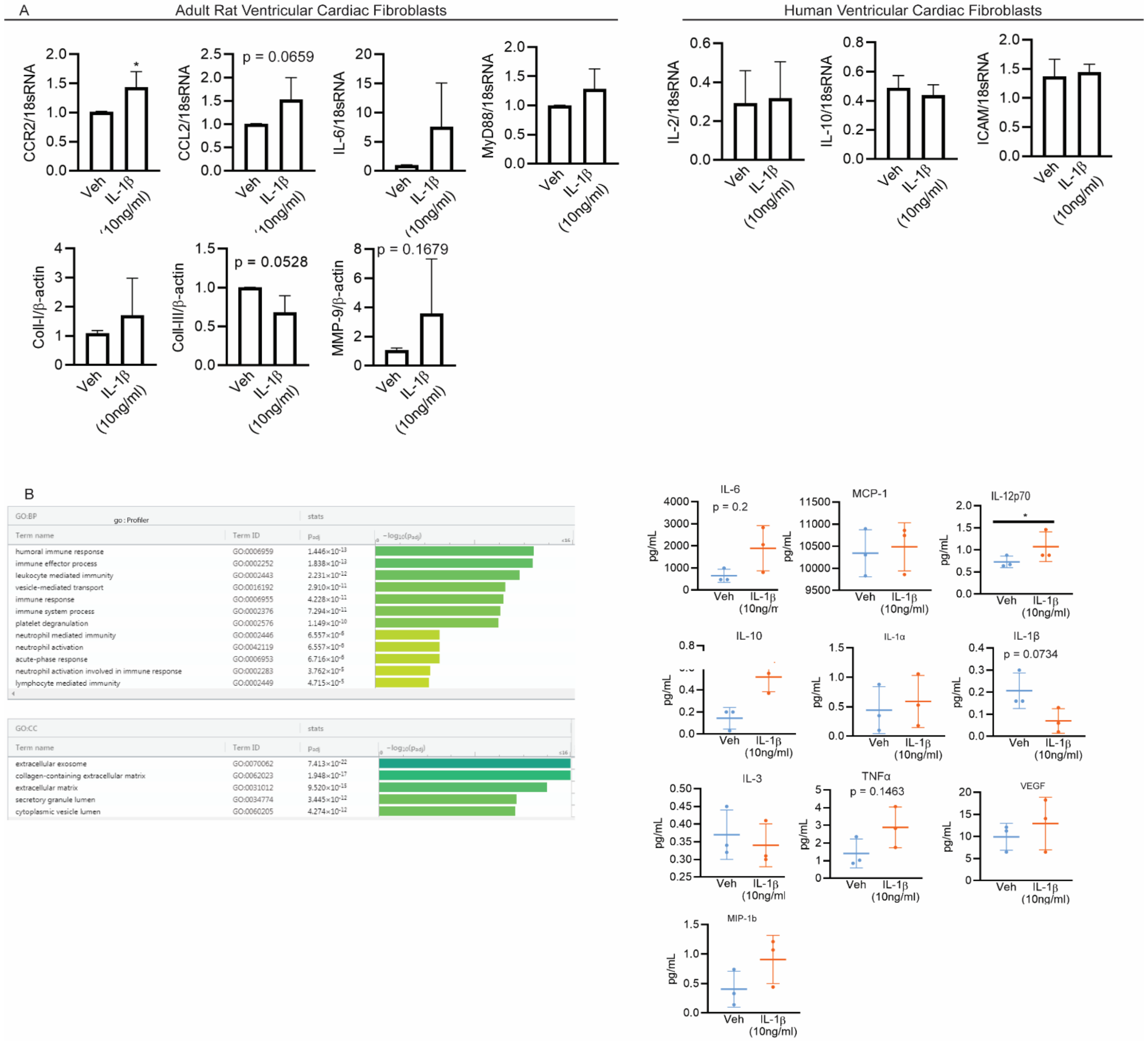

A
Adult Rat Ventricular Cardiac Fibroblasts
Human Ventricular Cardiac Fibroblasts
CCR2/18sRNA
CCL2/18sRNA
p = 0.0659
IL-6/18sRNA
MyD88/18sRNA
IL-2/18sRNA
IL-10/18sRNA
ICAM/18sRNA
Coll-I/β-actin
Coll-III/β-actin
p = 0.0528
MMP-9/β-actin
p = 0.1679
Veh
IL-1β (10ng/ml)
B
GO:BP
go : Profiler
stats
Term name
Term ID
$p_{adj}$
$-\log_{10}(p_{adj})$
humoral immune response GO:0006959 $1.446\times10^{-13}$
immune effector process GO:0002252 $1.838\times10^{-13}$
leukocyte mediated immunity GO:0002443 $2.231\times10^{-12}$
vesicle-mediated transport GO:0016192 $2.910\times10^{-11}$
immune response GO:0006955 $4.228\times10^{-11}$
immune system process GO:0002376 $7.294\times10^{-11}$
platelet degranulation GO:0002576 $1.149\times10^{-10}$
neutrophil mediated immunity GO:0002446 $6.557\times10^{-6}$
neutrophil activation GO:0042119 $6.557\times10^{-6}$
acute-phase response GO:0006953 $6.716\times10^{-6}$
neutrophil activation involved in immune response GO:0002283 $3.762\times10^{-5}$
lymphocyte mediated immunity GO:0002449 $4.715\times10^{-5}$
GO:CC
extracellular exosome GO:0070062 $7.413\times10^{-22}$
collagen-containing extracellular matrix GO:0062023 $1.948\times10^{-17}$
extracellular matrix GO:0031012 $9.520\times10^{-15}$
secretory granule lumen GO:0034774 $3.445\times10^{-12}$
cytoplasmic vesicle lumen GO:0060205 $4.274\times10^{-12}$
IL-6
p = 0.2
MCP-1
IL-12p70
IL-10
IL-1α
IL-1β
p = 0.0734
IL-3
TNFα
p = 0.1463
VEGF
MIP-1b
pg/mL

### Figure S16

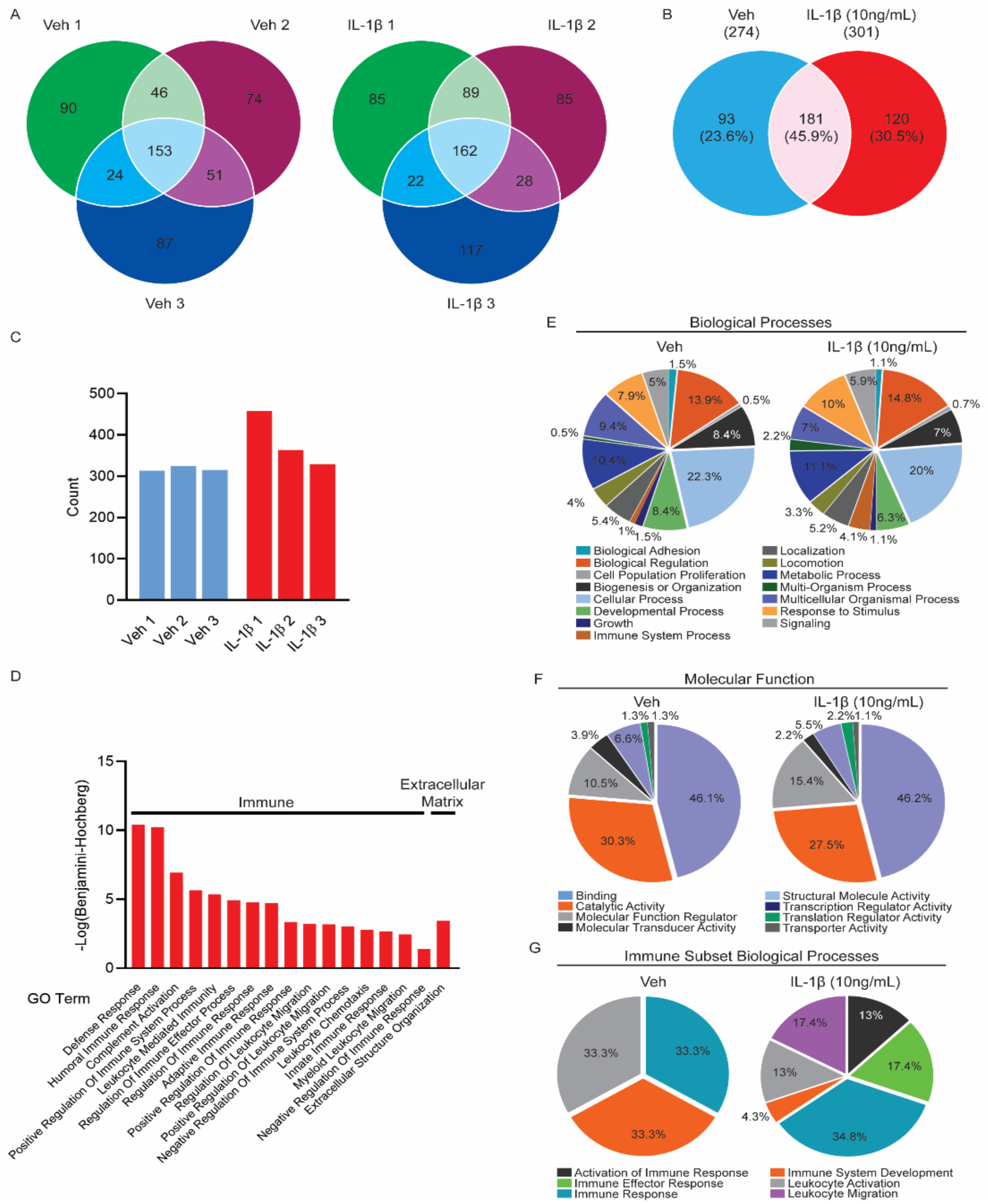
A
Veh 1
Veh 2
Veh 3
90
46
74
153
24
51
87
IL-1β 1
IL-1β 2
IL-1β 3
85
89
85
162
22
28
117
B
Veh
(274)
IL-1β (10ng/mL)
(301)
93
(23.6%)
181
(45.9%)
120
(30.5%)
C
Count
Veh 1
Veh 2
Veh 3
IL-1β 1
IL-1β 2
IL-1β 3
D
-Log(Benjamini-Hochberg)
Immune
Extracellular Matrix
GO Term
Defense Response
Humoral Immune Response
Complement Activation
Positive Regulation Of Immune System Process
Leukocyte Mediated Immunity
Regulation Of Immune Effector Process
Adaptive Immune Response
Regulation Of Immune Response
Positive Regulation Of Immune Response
Regulation Of Leukocyte Migration
Positive Regulation Of Leukocyte Migration
Negative Regulation Of Immune System Process
Leukocyte Chemotaxis
Innate Immune Response
Myeloid Leukocyte Migration
Negative Regulation Of Immune Response
Extracellular Structure Organization
E
Biological Processes
Veh
IL-1β (10ng/mL)
Biological Adhesion
Biological Regulation
Cell Population Proliferation
Biogenesis or Organization
Cellular Process
Developmental Process
Growth
Immune System Process
Localization
Locomotion
Metabolic Process
Multi-Organism Process
Multicellular Organismal Process
Response to Stimulus
Signaling
F
Molecular Function
Veh
IL-1β (10ng/mL)
Binding
Catalytic Activity
Molecular Function Regulator
Molecular Transducer Activity
Structural Molecule Activity
Transcription Regulator Activity
Translation Regulator Activity
Transporter Activity
G
Immune Subset Biological Processes
Veh
IL-1β (10ng/mL)
Activation of Immune Response
Immune Effector Response
Immune Response
Immune System Development
Leukocyte Activation
Leukocyte Migration

### Figure S17

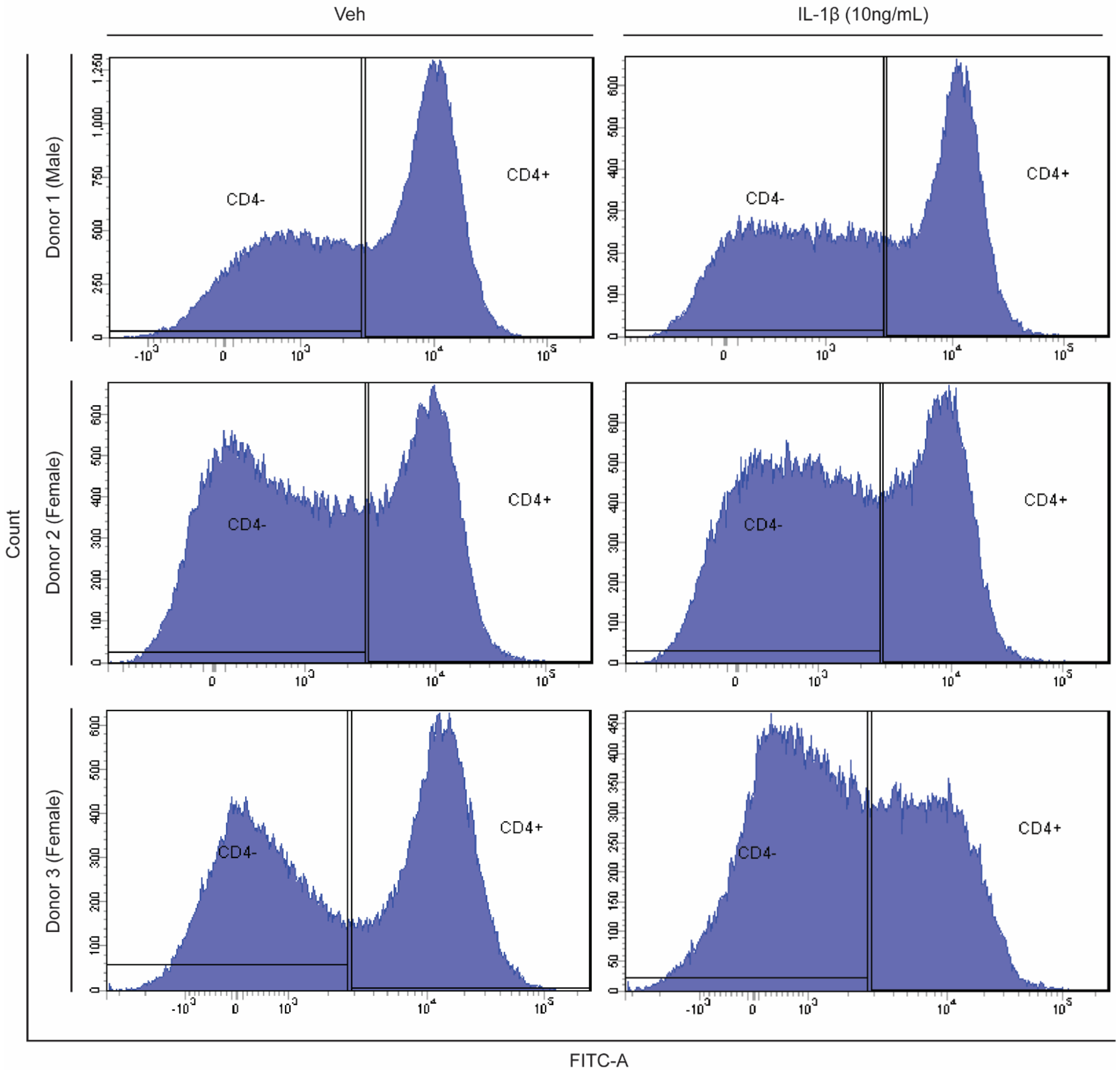
Veh
IL-1β (10ng/mL)
Donor 1 (Male)
Donor 2 (Female)
Donor 3 (Female)
Count
CD4-
CD4+
FITC-A

### Figure S18

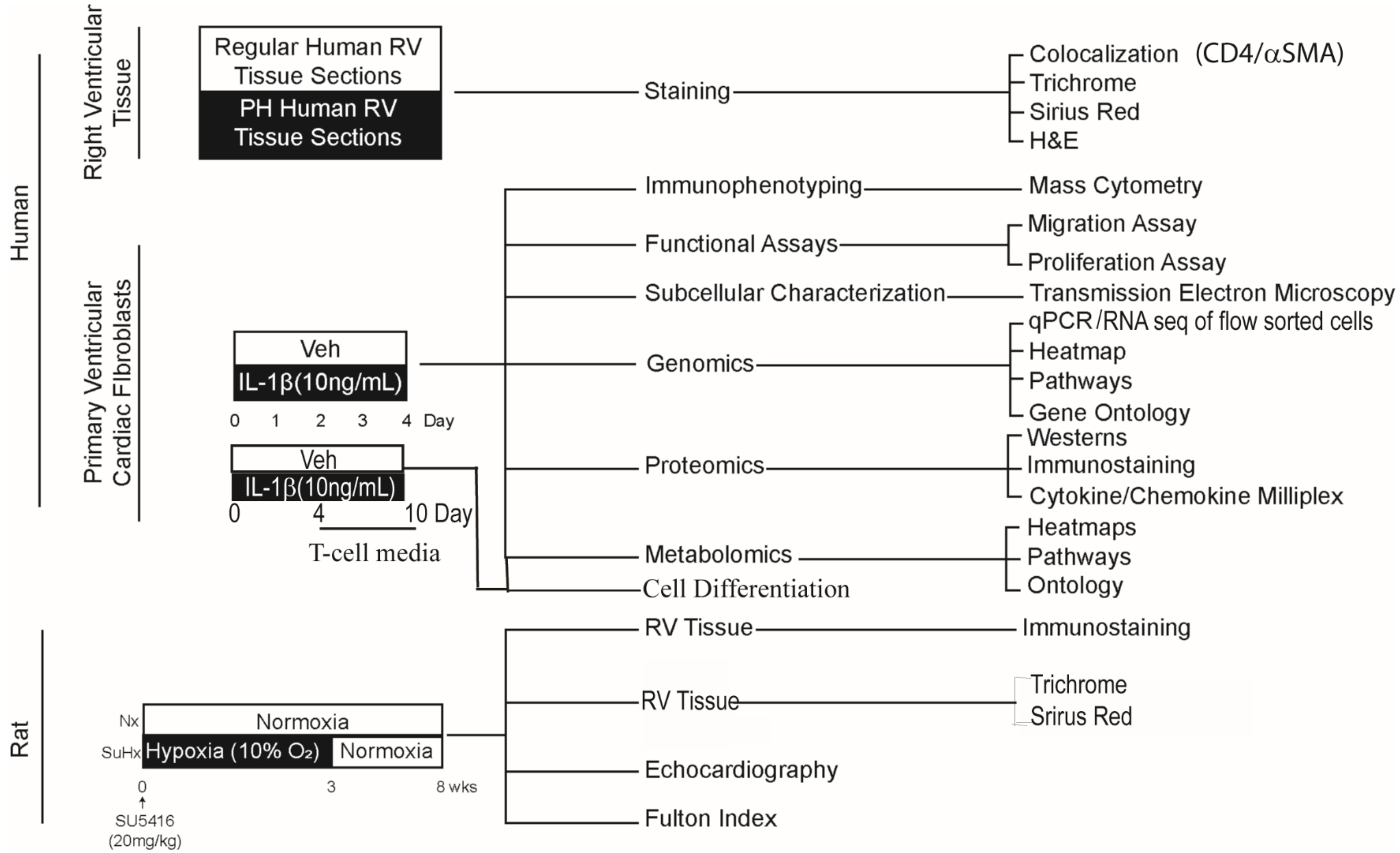

Human
Right Ventricular Tissue
Regular Human RV Tissue Sections
PH Human RV Tissue Sections
Staining
Colocalization (CD4/αSMA)
Trichrome
Sirius Red
H&E
Primary Ventricular Cardiac Fibroblasts
Immunophenotyping
Mass Cytometry
Functional Assays
Migration Assay
Proliferation Assay
Subcellular Characterization
Transmission Electron Microscopy
Veh
IL-1β(10ng/mL)
0 1 2 3 4 Day
Genomics
qPCR/RNA seq of flow sorted cells
Heatmap
Pathways
Gene Ontology
Veh
IL-1β(10ng/mL)
0 4 10 Day
T-cell media
Proteomics
Westerns
Immunostaining
Cytokine/Chemokine Milliplex
Metabolomics
Heatmaps
Pathways
Ontology
Cell Differentiation
Rat
RV Tissue
Immunostaining
Nx
Normoxia
SuHx
Hypoxia (10% O₂)
Normoxia
0 3 8 wks
SU5416 (20mg/kg)
RV Tissue
Trichrome
Srirus Red
Echocardiography
Fulton Index
