## Supplementary figure legends for "Identification of CD4^+^ Sub-population of Resident Cardiac Fibroblasts Linked to Myocardial Fibrosis"

**Figure S1. Characterization of primary human ventricular fibroblast using lineage specific markers and endothelial cell markers**. Immunostaining of passage 3 untreated 20,000 hVCF ((ID# 62122; male, ID# 1281202; male and ID#534282; female) with cardiac fibroblast specific markers; Collagen I (green), Periostin (red), Platelet Derived Growth Factor (PDGFRβ) (red), and Fibroblast Specific Protein (FSP-1) (green). Endothelial cell specific marker, Vascular Endothelial (VE) Cadherin (red) was used as negative control, Scale bar; 10 μm, 20 μm.

**Figure S2: Gating strategy of live, nucleated single cell for mass cytometry analysis.** (A) Gating strategy for the selection of single nucleated, viable cells used to generate viSNE graphs in Figure 1. Gating performed on cluster of events with cell staining for two isotopes of Iridium DNA intercalator and viability stain to remove debris and select intact cells. Further gating on event_length parameter was performed to remove cell aggregates to obtain "intact singlets". Next, "live cell" gating was employed to select cells with low staining with "thiol" reactive dye. The "event length" parameter refers to the time of detection of cell event and partially eliminates cell aggregates. From the ungated populations, 140Ce population was gated followed by gating on the residual subset, center subset, offset subset, width subset, event length subset, live-dead subset and gating of $Vim^+$ $\alpha SMA^+$ populations. The gating strategy using manual and cytofclean tool in R is presented for all the three human donors (ID# 62122; male, ID# 1281202; male and ID#534282; female) untreated or treated with IL-1β (10ng/mL).

**Figure S3: Expression of lymphoid markers on hVCF cell surface in response to interleukin-1-beta.** T-distributed Stochastic Neighbor Embedding (tSNE) plots representing the distribution and intensity of markers on primary human ventricular cardiac fibroblasts treated with Veh or IL-1β (10ng/mL) for 96 h and subjected to mass cytometry. (A) tSNE plots of the entire vehicle-treated population of cardiac fibroblast cells expressing CD16, CD4, CD183, CD45RA, CD3, CCR6, and CD45. (B) tSNE plots of the expanded population of cells from A marked by a red square. (C) tSNE plots of the entire IL-1β (10ng/mL) treated population of cardiac fibroblast cells expressing CD16, CD4, CD183, CD45RA, CD3, CCR6, and CD45. (D) tSNE plots of the expanded population of cells from C marked by a red square. The panels of each donor represent a total 2 x $10^6$ cardiac fibroblast cells profiled (n=3 males and a female).

**Figure S4**: **Uniform Manifold Approximation and Projection algorithm, a non-linear unsupervised learning model used to visualize the mass of all the markers by preserving the global structure**. UMAP, a non-linear unsupervised learning model used to visualize the mass of all the markers by preserving the global structure. The X and Y axis in the UMAP are dimensionless. UMAP which allows for the characterization of complex relationships between the marker by calculating the distances, neighbors and density performed on the mass cytometry data using PYTHON and R software tools. A summary of the donor population expressing all the cell surface markers is presented.

**Figure S5. Expression of lymphoid cells markers on cardiac fibroblast cells gated from Vimentin$^+$ αSMA$^+$ cells.** Flow cytometry panels showing the percentage of hVCF expressing lymphoid markers CD3$^+$ CD45RA$^+$, CD3$^+$ CD4$^+$, CD183$^+$ CD4$^+$, CCR6$^+$ CD4$^+$ cells in response to Veh or 48 h IL-1B (10ng/mL) treatment from all the three donors (ID# 62122; male, ID# 1281202; male and ID#534282; female) gated from Vimentin$^+$ αSMA$^+$ cells. The percentage of cells are indicated in quadrants.

**Figure S6. Expression of lymphoid cells markers on cardiac fibroblast cells gated from Vimentin$^+$ $\alpha$SMA$^-$ cells.** Flow cytometry panels showing the percentage of hVCF expressing lymphoid markers CD3$^+$ CD45RA$^+$, CD3$^+$ CD4$^+$, CD183$^+$ CD4$^+$, CCR6$^+$ CD4$^+$ cells in response to Veh or 48 h IL-1$\beta$ (10ng/mL) treatment from all the three donors (ID# 62122; male, ID# 1281202; male and ID#534282; female) gated from Vimentin$^+$ $\alpha$SMA$^-$ cells. The percentage of cells is indicated in quadrants.

**Figure S7. Expression of lymphoid cells markers on cardiac fibroblast cells gated from Vimentin$^-$ $\alpha$SMA$^+$ cells.** Flow cytometry panels showing the percentage of hVCF expressing lymphoid markers CD3$^+$ CD45RA$^+$, CD3$^+$ CD4$^+$, CD183$^+$ CD4$^+$, CCR6$^+$ CD4$^+$ cells in response to Veh or 48 h IL-1$\beta$ (10ng/mL) treatment from all the three donors (ID# 62122; male, ID# 1281202; male and ID#534282; female) gated from Vimentin$^-$ $\alpha$SMA$^+$ cells. The percentage of cells are indicated in quadrants.

**Figure S8 Percentages of CD4 hVCF expressing either Vimentin$^+$ or $\alpha$SMA$^+$ or both.** This supplementary figure is a part of Figure 2 showing histograms of populations of cells expressing Vimentin$^+$ or $\alpha$SMA$^+$ in response to Veh and IL-1$\beta$. Mass cytometry 2D plots of total frequencies of $\alpha$SMA and Vimentin expressing hVCF. The red arrow from quadrant 1 indicate the frequencies CD4$^+$ cells among the Vimentin$^-$ $\alpha$SMA$^+$ activated resident cardiac fibroblast population. The arrows from quadrant 3 indicate the frequencies CD4$^+$ cells among the Vimentin$^+$ $\alpha$SMA$^-$ quiescent resident cardiac fibroblast population. The arrows from quadrant 2 indicate the frequencies of CD4$^+$ cells among the Vimentin$^+$ $\alpha$SMA$^+$ population.

**Figure S9. Characterization and validation of anti-human CD4 or anti-rat CD4 or anti-mouse-CD4 FITC antibody.** (A) Representative image of validation of CD4 antibody from humans and rats used in the experiments reported in this manuscript. The white lines indicates the splenic follicular structure surrounded by the lymphocytes. The white arrows point to the T cells (B) Validation of the antibodies using flow cytometry 2D images of rat blood stained with CD4 (ABCAM #ab133616) antibody used against human cells or rat tissue or mice tissue using a negative control and isotype control. Scale bar is 10μM.

**Figure S10: Tricuspid annular plane systolic excursion (TAPSE) tracings and Pulmonary outflow PW tracings using echocardiography.** (A) TAPSE tracings in a rat model of Sugen hypoxia taken at baseline and after 8 weeks using echocardiography (B) Pulmonary outflow PW Doppler tracings of a rat model of Sugen hypoxia taken at baseline and after 8 weeks using echocardiography.

**Figure S11. Characterization of SUGEN/hypoxia rat model of pulmonary hypertension.** Male Fischer rats from Charles River were given a single bolus of SUGEN (20mg/kg) and exposed to hypoxia (10% $FiO_2$) for three weeks, followed by exposure to normoxia (Nx) for an additional five weeks (SuHx). In parallel, control animals were exposed to normoxia (room air) for 8 weeks. Each Nx and SuHx group had n=10 rats. (A) Representative images of collagen content as measured by percentage of Sirius red positive regions in the transverse region of the right ventricle of Nx and SuHx rats (10 μm thickness), whole heart section cut transversely at the mid-ventral region and perivascular and interstitial regions of both Nx and SuHx animals. The arrows suggest fibrosis in the tissue (B) Representative images of Masson trichrome staining whole heart section and perivascular and interstitial regions of both Nx and SuHx animals. The arrows suggests fibrosis in the tissue. (C) Body weight and of Nx and SuHx animals measured using echocardiography. Values are mean ± SD (n=10). *$P$<0.05 (D) Fibroblast specific protein (FSP-1) staining of cardiac fibroblast in rat RV of both Nx and SuHx groups. Scale bars, 50 μm for the picosirus red and masson trichrome staining images. Values are mean ± SD (n=10). The comparisons were made with unpaired t test. *$P<0.05$ vs Nx rats. Scale bars, Whole tissue transverse section 1mm, perivascular and interstitial region, 50 μM.

**Figure S12: IL-1β mediated expansion of CD4 expressing cells detected using single cell mass cytometry** (A) Heatmap of marker intensities of all the 26 markers that were used in the mass cytometry analysis of resident primary hVCF. (B) Single cells expression of CD4 in Veh and IL-1β for all the 3 donors. (C) Quantification of all the markers in Veh and IL-1β represented as a non-redundancy score. (D) Quantification of the HLA-DR, and myeloid and other cell populations treated with Veh and IL-1β based on the heatmap. (E) Normalized density expression based on the heatmap of all the markers in Veh and IL-1β treated cells measured using mass cytometry. n=3 biological replicates.

**Figure S13: Collagen Iα and Periostin expression in human cardiac fibroblast cells with IL-1β.** Top Panel: Low magnification images of Veh or 24 h IL-1β(10ng/mL) treated human cardiac fibroblast cells immunostained with Collagen 1α or Periostin. Bottom Panel: High magnification images of Veh or 24 h IL-1β (10ng/mL) treated human cardiac fibroblast cells immunostained with Collagen 1α or Perisotin. Scale bars 20μM and 100μM.

**Figure S14: Appearance of secretory vesicle and clusters of hVCF cells with IL-1β.** (A) Transmission electron microscopy micrograph of ultramicroscopic subcellular structures of the vehicle and IL-1β (10ng/ml) 24 h treated hVCF. N: Nucleus; ER: Endoplasmic Reticulum; MV: Secreted Microvesicle; G: Golgi body; M: Mitochondria. Scale bar, 400nm. n=3 technical replicates. (B) Schematic representation of cardiac fibroblast 14 days differentiation in T-cell media. hVCF were treated with rIL-1b (10ng/mL) in fibroblast media for 4 days.

Cell were then transferred to T-cell expansion media with T-cell activators (CD3, CD2, CD28) for 10 days. Bright field images of cardiac fibroblasts grown in T-cell media with T-cell activators with or without IL-1β (10ng/mL) for 13 days. Multicellular clusters marked by arrows were seen in the IL-1β treated wells but not in the vehicle treated wells. Quantification of the multi-cell rosettes clusters is represented as a scatter plot of Mean ± SD values from individual subjects (n=3, *$P$ <0.05 Kruskal-Wallis one-way ANOVA with Dunn's multiple comparisons).

**Figure S15: IL-1β Modulates inflammatory genes and extracellular genes in hVCF.** (A) Validation of gene involved in inflammation such as CCR2, CCL2, IL-6, TGFβ, MyD88 and IL-1RA were validated in primary adult rat ventricular cardiac fibroblasts. Further validation of genes involved in inflammation and extracellular matrix (CD4, IL-2, IL-8, MMP-9, IL-10, ICAM, Coll-I and Coll-III) was performed in primary hVCF treated with Veh or IL-1β (10ng/mL) after 24h using quantitative PCR (qPCR). (B) Quantification of cytokines and chemokines released in the condition media from Veh and IL-1β treated cardiac fibroblasts. The scatterplots represent Mean ± SD values (n=3 biological replicates, *$P$<0.05 by unpaired-student t test with Mann Whitney post hoc test. Gene ontology functional analysis using g:profiler shows the gene ontology terms for biological processes and cellular component categorizing the secreted proteins in the conditioned media identified using LC-MS/MS. The graph represents Mean ± SD values (n=3, *$P$<0.05 by unpaired-student t test with Mann Whitney post hoc test).

**Figure S16: Non-immune cardiac fibroblasts secrete immunomodulatory proteins in response to IL-1β treatment.** (A) Multivariable proteomic profile of proteins in technical replicates of cardiac fibroblasts (ID# 62122) conditioned media volumes (1 mL, 2 mL and 4 mL). Venn diagram represents proteins secreted in the conditioned media of primary human ventricular cardiac fibroblasts treated with Veh or 24 h IL-

1β (10 ng/mL) from three samples determined using Venny v2.0. (B) Venn diagram represents proteins that are unique to either the Veh or IL-1β or proteins common to both the conditions. (C) Average number of proteins detected in the conditioned media of cardiac fibroblasts treated with Veh or IL-1β for each of the replicate (D) Pathway enrichment analysis of the metabolites differentially expressed with IL-1β treatment in DAVID (https://david.ncifcrf.gov/). The length of the bar corresponds to significance of the pathway based on the –Log q value and the multiplicity of the components corrected using Benjamini-Hochberg. (E) Pie charts based on gene ontology showing the number and proportion of secreted proteins that belong to different biological processes in both Veh and IL-1β groups determine using PANTHER (http://www.pantherdb.org/) (F) Pie charts showing the number and proportion of secreted proteins that belong to different molecular function in both Veh and IL-1β groups. (G) Pie charts showing the number and proportion of secreted proteins that belong to different immune subsets in both Veh and IL-1β groups. Secretory protein belonging to the functional process, leukocyte migration and activation of immune response were distinctly upregulated with IL-1β.

**Figure S17:** Graphical representation of untreated or IL-1β treated human ventricular cardiac fibroblast sorted into $CD4^{+}$ and $CD4^{-}$ population for RNA seq analysis for all the three human donor cells.

**Figure S18: Summary of the methods and models used to identify the cardiac fibroblast subset expressing mesenchymal and lymphoidal markers.** We assessed rat PH model and human right ventricular cells and tissue through complementary methods, including immunophenotyping, genomics, proteomics, and metabolomics, in order to determine the mechanism by which resident cardiac fibroblasts assume an immune phenotype.
