## Supplementary table for "Identification of CD4^+^ Sub-population of Resident Cardiac Fibroblasts Linked to Myocardial Fibrosis"

**Supplementary Tables**

**Table S1:** Overview of characteristics of donors of primary human ventricular cardiac fibroblasts

| **Human Donor #ID** | **Age**  **(yr)** | **Sex** | **Isolation Location** | **Medications** | **Diabetes** | **Alcohol** | **Smoking** |
| --- | --- | --- | --- | --- | --- | --- | --- |
| 67771 | 67 | F | Left Ventricle | Xanax, Prozac, Neurontin, Diluad ID, Mobic, Morphine, Zyprexa, Oxycodone, Flomax, Levophed, Vancomycin, Clindamycin | No | Yes | 1 pack per day, 30+ years |
| 62122 | 73 | M | Left Ventricle | None | No | Yes | No |
| 1281202 | 67 | M | Left Ventricle | Neosenephrine, Solumedrol, Lasix, T4, Zosyn, Meropenem, Cleocin | No | No | No |
| 534282 | 63 | F | Left Ventricle | None | No | Yes | Yes |
| TL210281 | 73 | M | Left Ventricle | HTN, Lupus Medication | No | Yes | 1-2 packs per day on opiates per tox screen |

**Table S2:** Antibodies used for immunostaining and immunoblotting *in vitro* and *in vivo* experiments

| **Antibody (Ab)** | **Clone** | **Source** | **Identifier (Cat#)** |
| --- | --- | --- | --- |
| α-actin smooth muscle actin (mouse) | 1A4 | Sigma-Aldrich | A2547 |
| IL1 Receptor I/IL-1R-1 (rabbit) | N/A | Abcam | ab106278 |
| β-actin (mouse) | AC-15 | Sigma-Aldrich | A5441 |
| Vinculin (rabbit) | N/A | Santa Cruz Biotechnology | sc-5573 |
| Periostin (mouse) | 5B2.3 | Sigma-Aldrich | MABS1183 |
| S100A4/FSP1 (recombinant) | EPR2761(2) | Abcam | ab124805 |
| PDGFR-β (mouse) | D-6 | Santa Cruz Biotechnology | sc-374573 |
| Collagen I (rabbit) | N/A | Abcam | ab34710 |
| VE-cadherin (mouse) | F-8 | Santa Cruz Biotechnology | sc-9989 |
| CD4 (rabbit mAB) | EPR6855 | Abcam | ab133616 |
| CD68 (mouse) | KP1 | Abcam | ab955 |
| Vimentin (mouse) | AMF-17b | Developmental Studies Hybridoma Bank | AMF-17b |
| CD3-APC (rat) | REA223 | Miltenyi Biotec | 130-103-132 |
| CD4 (Domain 1)-PE (rat) | REA482 | Miltenyi Biotec | 130-107-668 |
| p-p38 MAPK (Thr180/Tyr182) (rabbit mAB) | D3f9 | Cell Signaling Technologies | 4511S |
| p38 MAPK (rabbit mAB) | D13E1 | Cell Signaling Technologies | 8690S |
| p-p65NFκB (Ser 536) (rabbit mAB) | 93H1 | Cell Signaling Technologies | 3033 |
| NFκB p-65 (rabbit mAB) | D14E12 | Cell Signaling Technologies | 8242 |
| CD4-FITC (mouse) | RMA-5 | Biolegend | 100510 |

**Table S3:** Heavy metal conjugated antibodies used for immunostaining and immunoblotting *in vitro* and *in vivo* experiments.

| **Ab #** | **Target** | **Metal tag Ab (amu)** | **Marker functions** | **Ab Clone** | **Catalog number** | **Company name** |
| --- | --- | --- | --- | --- | --- | --- |
| 1 | α-SMA | ^141^Pr | Myofibroblast marker | 1A4 | 3141017D | Fluidigm |
| 2 | Vimentin | ^143^Nd | Cardiac fibroblast marker | D21H3 | 3143027D | Fluidigm |
| 3 | CD11c | ^147^SM | Dendritic cell marker | Bu15 | 3147008B | Fluidigm |
| 4 | CD16 | ^148^Nd | Neutrophil | 3G8 | 3148004B | Fluidigm |
| 5 | CD11b | ^149^Sm | Macrophages and Microglia | EPR1344 | 3149028D | Fluidigm |
| 6 | PD-L1 | ^150^Nd | Expansion of CD4 and CD8-T cells | E1L3N | 3150031D | Fluidigm |
| 7 | CD69 | ^153^Eu | Activation marker on hematopoietic stem cells | FN50 | V00399 | Long Wood |
| 8 | IL-1β | ^154^Sm | Adaptive Immunity | H1b-27 | V05038 | Long Wood |
| 9 | CD4 | ^155^Gd | T-helper cell marker | RPA T4 | V04286 | Long Wood |
| 10 | CD183 | ^156^Gd | Effector T-cells | G025H7) | 3156004B | Fluidigm |
| 11 | CD33 | ^158^Gd | Myeloid marker | WM53 | 3158001B | Fluidigm |
| 12 | CD68 | ^159^Tb | Classical macrophage | KP1 | 3159035D | Fluidigm |
| 13 | CD14 | ^160^Gd | LPS receptor | M5E2 | 3160001B | Fluidigm |
| 14 | CTLA-4 | ^161^Dy | Activated T-lymphocyte | 14D3 | 3161004B | Fluidigm |
| 15 | CCR2 | ^163^Dy | Resident macrophage | K036C2 | V17774 | Long Wood |
| 16 | Arginase1 | ^164^Dy | Alternatively activated macrophages | 3164027D | D4E3M | Fluidigm |
| 17 | IL-2 | ^166^Er | T-cell growth factor | MQ1-17H12 | 3166002B | Fluidigm |
| 18 | CD27 | ^167^Er | Co-stimulatory marker expressed on T-cell, B-cell and NK cell | O323 | 3167002B | Fluidigm |
| 19 | CD206 | ^168^Er | Alternatively activated macrophages | 15-2 | 3168008B | Fluidigm |
| 20 | CD45RA | ^169^Tm | Helper T-cell | HI100 | 3169008B | Fluidigm |
| 21 | CD3 | ^170^Er | CD3 T-cell co-receptor | UCHT1 | 3170001B | Fluidigm |
| 22 | CCR6 | ^171^Yb | Functional marker for Th17 cell | G034E3 | 353427 | Biolegend |
| 23 | PHH3 | ^175^Lu | Cell proliferation marker | 3176023D | HTA28 | Fluidigm |
| 24 | CD127/IL-7RA | ^176^Yb | Treg cell marker | A019D5 | 3176004B | Fluidigm |
| 25 | HLA-DR | ^174^Yb | Cytotoxic-T-cell activation | L243 | 3174001B | Fluidigm |
| 26 | CD45 | ^89^Y | Pan immune cell marker | HI30 | 3089003B | Fluidigm |

αSMA: alpha smooth muscle actin; IL-1β: Interleukin 1 beta; HLA-DR: human leukocyte antigen-DR; PD-L1: Programmed death ligand 1

**Table S4.** Baseline characteristics of patient donors of right ventricular tissue

| **Patient ID#** | **Age at Death (yr)** | **Sex** | **Cause of Death** | **Final Anatomic Diagnosis** |
| --- | --- | --- | --- | --- |
| A | 63 | M | Progressive supranuclear palsy, leading to dysphagia, complicated by aspiration pneumonia | Not available |
| B | 66 | M | Atherosclerotic and hypertrophic cardiovascular disease | Echocardiography showed dilated right ventricle. At autopsy, there were many signs of atherosclerotic and hypertensive changes.Atherosclerotic and hypertrophic cardiovascular disease, biventricular cardiac hypertrophy (500 g), remote myocardial infarct, posterior papillary muscle ,coronary artery atherosclerosis, left main coronary artery at origin: 50-75% occluded left anterior descending artery: 75-100% occluded, left circumflex artery: 50% occluded, right coronary artery: 50% occluded pulmonary artery atherosclerosis (consistent with sustained pulmonary hypertension) right ventricle dilatation, arterionephrosclerosis |
| C | 61 | F | Metastatic dedifferentiated endometroid adenocarcinoma of ovary complicated by intraperitoneal hemorrhage from liver biopsy | Not available |
| D | 92 | M | Cardiac amyloidosis leading to restrictive cardiomyopathy complicated by atherosclerotic and hypertensive cardiovascular disease | Amyloid cardiomyopathy, multifocal cardiac amyloid deposition confirmed with congo red stain, atherosclerotic and hypertrophic cardiovascular disease (clinical history of systemic and pulmonary hypertension), cardiomegaly (500 gm) with biventricular hypertrophy coronary artery atherosclerosis, moderate: left anterior descending artery, 20-40% stenosis right coronary artery, 40% stenosis, subendocardial infarct, remote, posterior papillary muscle, left ventricle |
| E | 32 | M | Acute asthmatic attack | Not available |
| F | 65 | M | Primary pulmonary hypertension, leading to right ventricular dilation and hypertrophy, leading to right heart failure | primary pulmonary hypertension, cardiomegaly (890 gm), with right ventricular dilation and biventricular hypertrophy, thickening and enlargement of the tricuspid valve, arterio and arteriolonephrosclerosis, bilateral kidneys, focal segmental glomerolusclerosis, bilateral kidneys, cystic emphysematous blebs, bilateral lungs, petechial hemorrhage, diffuse, stomach, multiple cortical cysts (up to 2.5 cm), bilateral kidneys atherosclerosis, thoracic and abdominal aorta |

**Table S5:** Primer sequence of genes amplified to determine the gene expression levels in vehicle and IL1β treated rat cardiac fibroblast cells.

|  | **Forward Primer** | **Reverse Primer** |
| --- | --- | --- |
| **Rat** |  |  |
| Col-1 | 5'-TGCCGTGACCTCAAGATGTG-3′ | 5'-CACAAGCGTGCTGTAGGTGA-3′ |
| Col-III | 5'-ATGGTGGCTTTCAGTTCAGC-3′ | 5'-TGGGGTTTCAGAGAGTTTGG-3′ |
| MMP-9 | 5'-GTACAGCCTGTTTCTGGTGGC-3' | 5'-GGCCTTGGGTCAGGTTTAGAG-3' |
| MyD88 | 5'-TTCTCCAACGCTGTCCTGTC-3' | 5'-AACTGAGATGTGTGCCCAGG-3' |
| IL-6 | 5'-TTGTTGACAGCCACTGCCTTCCC-3' | 5'-TCTGACAGTGCATCATCGCTGTTCA-3' |
| CCL2 | 5’-GCCCCACTCACCTGCTGCTAC-3’ | 5’-GGTTCTGATCTCATTTGGTTCCG-3’ |
| CCR2 | 5'-ATCCACGGCATACTATCAACATCTC-3’ | 5’-GACAAGGCTCACCATCATCGTAG-3' |
| IL1R | 5’-CTT GCC GCA CGT CCT ACA CAT ACC-3’ | 5’-CGG GGA AGA AAA TCA GAG CAG GAG-3’ |
| **Housekeeping genes** |  |  |
| GAPDH | 5'-CGCTAACATCAAATGGGGTG-3′ | 5'-TTGCTGACAATCTTGAGGGAG-3′ |
| 18sRNA | 5’-AAACGGCTACCACATCCA-3’ | 5’-CTCATTCCAATTACAGGG-3’ |
| **Human** |  |  |
| IL-2 | 5'-CAT TGC ACT AAG TCT TGC ACT TGT CA-3' | 5'-CGT TGA TAT TGC TGA TTA AGT CCC TG-3' |
| IL-6 | 5'-ATGAACTCCTTCTCCACAAGCGC-3' | 5'-GAAGAGCCCTCAGGCTGGACTG-3' |
| IL-8 | 5'-TGACTTCCAAGCTGGCCGTGGCT-3' | 5'-TCTCAGCCCTCTTCAAAAACTTCTC-3' |
| IL-10 | 5'-AAGCTGAGAACCAAGACCCAGACATCAAGGCG-3' | 5'-AGCTATCCCAGAGCCCCAGATCCGATTTTGG-3' |
| ICAM 1 | 5'-GAT TGT CAT CAT CAC TGT GGT AG-3' | 5'-GCC TGT TGT AGT CTG TAT TTC TT-3' |
| CD4 | 5'- GGA GTC CCT TTT AGG CAC TTG C -3' | 5'-AAGACAGTGCATGTCCAGGTG-3' |
| TGF-B | 5'-GGCCCTGCCCCTACATTT-3' | 5'-CCGGGTTATGCTGGTTGTACA-3' |
| **Housekeeping genes** |  |  |
| β-Actin | 5’-CTTTCGTGTAAATTATGTAATGCA-3’ | 5’-TACATCTCAAGTTGGGGGA-3’ |
| GAPDH | 5'-ACCACAGTCCATGCCATCAC-3' | 5'-TCCACCACCCTGTTGCTGTA-3' |

**Table S6: Summary of differentially regulated extracellular matrix genes in the Veh and IL-1β (10ng/mL) treated hVCF using taqman gene array**

| **Gene** | **Gene name** | **2^-ΔCt (Veh)** | **2^-ΔCt (IL-1B)** | **Standard deviation (Veh)** | **Standard deviation (IL-1B)** | **Fold Change (IL-1B/Veh)** | **P value** | **(-logP)** | **FDR q Value** |
| --- | --- | --- | --- | --- | --- | --- | --- | --- | --- |
| *NCAM1* | neural cell adhesion molecule 1 | 0.75 | 0.41 | 1.16 | 0.14 | 0.55 | 0.17 | 0.78 | 10.00 |
| *TNC* | tenascin C | 4.47 | 10.87 | 2.95 | 9.20 | 2.43 | 0.25 | 0.60 | 7.53 |
| *FN1* | fibronectin 1 | 20.34 | 40.36 | 20.26 | 0.00 | 1.98 | 0.28 | 0.56 | 5.54 |
| *COL12A1* | collagen type XII alpha 1 chain | 2.44 | 6.71 | 2.50 | 5.77 | 2.75 | 0.31 | 0.51 | 4.64 |
| *CD44* | CD44 molecule | 3.29 | 8.79 | 3.32 | 7.59 | 2.67 | 0.32 | 0.50 | 3.79 |
| *MMP12* | matrix metallopeptidase 12 | 4.28 | 48.63 | 3.03 | 70.22 | 11.36 | 0.32 | 0.49 | 3.24 |
| *THBS3* | thrombospondin 3 | 72.39 | 0.25 | 41.93 | 0.24 | 0.00 | 0.38 | 0.43 | 3.21 |
| *COL7A1* | collagen type VII alpha 1 chain | 4.34 | 0.41 | 2.89 | 0.14 | 0.10 | 0.38 | 0.42 | 2.85 |
| *PGK1* | phosphoglycerate kinase 1 | 0.76 | 1.68 | 1.17 | 0.58 | 2.21 | 0.41 | 0.39 | 2.73 |
| *MMP14* | matrix metallopeptidase 14 | 2.25 | 5.39 | 2.67 | 4.65 | 2.39 | 0.41 | 0.38 | 2.48 |
| *ITGB3* | integrin subunit beta 3 | 2.53 | 5.01 | 2.12 | 4.94 | 1.98 | 0.42 | 0.38 | 2.28 |
| *ACTB* | actin beta | 1.03 | 1.98 | 0.83 | 1.98 | 1.93 | 0.42 | 0.38 | 2.10 |
| *ITGB2* | integrin subunit beta 2 | 7.89 | 0.27 | 5.67 | 0.13 | 0.03 | 0.44 | 0.36 | 2.01 |
| *CTNNA1* | catenin alpha 1 | 1.08 | 2.59 | 1.33 | 2.34 | 2.41 | 0.44 | 0.35 | 1.89 |
| *COL14A1* | collagen type XIV alpha 1 chain | 0.70 | 0.75 | 1.22 | 0.77 | 1.06 | 0.48 | 0.32 | 1.91 |
| *COL4A2* | collagen type IV alpha 2 chain | 8.03 | 16.16 | 8.02 | 16.12 | 2.01 | 0.48 | 0.32 | 1.79 |
| *UBC* | ubiquitin C | 4.03 | 8.17 | 4.05 | 8.17 | 2.02 | 0.48 | 0.32 | 1.69 |
| *ITGA1* | integrin subunit alpha 1 | 1.98 | 4.11 | 2.07 | 4.08 | 2.07 | 0.48 | 0.32 | 1.60 |
| *THBS2* | thrombospondin 2 | 4.02 | 8.08 | 4.02 | 8.12 | 2.01 | 0.48 | 0.32 | 1.52 |
| *MMP7* | matrix metallopeptidase 7 | 0.06 | 13.29 | 0.12 | 22.20 | 235.17 | 0.48 | 0.31 | 1.45 |
| *ITGB4* | integrin subunit beta 4 | 2.61 | 0.81 | 1.96 | 0.37 | 0.31 | 0.49 | 0.31 | 1.41 |
| *ITGA6* | integrin subunit alpha 6 | 4.51 | 1.01 | 2.99 | 0.86 | 0.22 | 0.50 | 0.30 | 1.35 |
| *CNTN1* | contactin 1 | 37.21 | 5.63 | 21.99 | 8.88 | 0.15 | 0.50 | 0.30 | 1.30 |
| *COL15A1* | collagen type XV alpha 1 chain | 0.78 | 0.84 | 1.17 | 0.30 | 1.08 | 0.53 | 0.28 | 1.32 |
| *MMP8* | matrix metallopeptidase 8 | 0.13 | 0.31 | 0.16 | 0.26 | 2.37 | 0.53 | 0.28 | 1.27 |
| *ITGA7* | integrin subunit alpha 7 | 8.46 | 13.50 | 6.76 | 16.86 | 1.60 | 0.57 | 0.24 | 1.31 |
| *MMP1* | matrix metallopeptidase 1 | 2.01 | 3.41 | 1.76 | 4.17 | 1.69 | 0.57 | 0.24 | 1.27 |
| *PECAM1* | platelet and endothelial cell adhesion molecule 1 | 1.98 | 3.35 | 1.74 | 4.07 | 1.69 | 0.57 | 0.24 | 1.23 |
| *SELL* | selectin L | 1.93 | 3.10 | 1.73 | 4.08 | 1.60 | 0.63 | 0.20 | 1.29 |
| *COL1A1* | collagen type I alpha 1 chain | 11.33 | 19.72 | 13.09 | 19.88 | 1.74 | 0.64 | 0.19 | 1.28 |
| *MMP2* | matrix metallopeptidase 2 | 1.83 | 3.28 | 2.21 | 3.25 | 1.79 | 0.65 | 0.19 | 1.25 |
| *ADAMTS13* | ADAM metallopeptidase with thrombospondin type 1 motif 13 | 0.78 | 1.52 | 1.18 | 0.88 | 1.94 | 0.65 | 0.19 | 1.22 |
| *MMP11* | matrix metallopeptidase 11 | 7.37 | 2.36 | 4.56 | 3.55 | 0.32 | 0.67 | 0.18 | 1.21 |
| *SPP1* | secreted phosphoprotein 1 | 2.63 | 1.24 | 1.99 | 1.05 | 0.47 | 0.67 | 0.17 | 1.19 |
| *SELE* | selectin E | 119.05 | 34.97 | 68.81 | 59.14 | 0.29 | 0.68 | 0.17 | 1.17 |
| *MMP10* | matrix metallopeptidase 10 | 15.01 | 4.72 | 8.87 | 7.09 | 0.31 | 0.69 | 0.16 | 1.15 |
| *ICAM1* | intercellular adhesion molecule 1 | 11.32 | 3.90 | 6.76 | 5.17 | 0.34 | 0.71 | 0.15 | 1.15 |
| *LAMA3* | laminin subunit alpha 3 | 4.47 | 1.86 | 2.96 | 1.90 | 0.42 | 0.71 | 0.15 | 1.13 |
| *RPLP0* | ribosomal protein lateral stalk subunit P0 | 3.30 | 4.42 | 3.32 | 3.78 | 1.34 | 0.72 | 0.14 | 1.11 |
| *TGFB1* | transforming growth factor beta 1 | 12.41 | 16.42 | 12.31 | 14.22 | 1.32 | 0.73 | 0.14 | 1.09 |
| *LAMC1* | laminin subunit gamma 1 | 8.07 | 10.68 | 8.03 | 9.24 | 1.32 | 0.73 | 0.14 | 1.06 |
| *SGCE* | sarcoglycan epsilon | 5.03 | 6.67 | 5.02 | 5.77 | 1.33 | 0.73 | 0.14 | 1.04 |
| *ITGAV* | integrin subunit alpha V | 1.98 | 2.72 | 2.07 | 2.30 | 1.38 | 0.73 | 0.14 | 1.02 |
| *ITGB1* | integrin subunit beta 1 | 8.15 | 10.74 | 8.14 | 9.30 | 1.32 | 0.73 | 0.13 | 1.00 |
| *KAL1* | anosmin1 | 1.56 | 2.21 | 1.72 | 1.87 | 1.42 | 0.74 | 0.13 | 0.99 |
| *COL5A1* | collagen type V alpha 1 chain | 0.77 | 1.41 | 1.17 | 1.01 | 1.83 | 0.75 | 0.12 | 0.98 |
| *MMP3* | matrix metallopeptidase 3 | 18.20 | 7.31 | 11.03 | 7.49 | 0.40 | 0.76 | 0.12 | 0.97 |
| *VTN* | vitronectin | 1.08 | 1.66 | 1.33 | 1.37 | 1.53 | 0.76 | 0.12 | 0.95 |
| *MMP16* | matrix metallopeptidase 16 | 1.10 | 1.04 | 1.25 | 0.36 | 0.94 | 0.76 | 0.12 | 0.94 |
| *LAMA2* | laminin subunit alpha 2 | 1.08 | 1.02 | 1.23 | 0.36 | 0.95 | 0.76 | 0.12 | 0.92 |
| *COL11A1* | collagen type XI alpha 1 chain | 1.12 | 1.14 | 1.37 | 1.19 | 1.02 | 0.82 | 0.08 | 0.97 |
| *TIMP3* | TIMP metallopeptidase inhibitor 3 | 10.37 | 9.89 | 8.26 | 9.87 | 0.95 | 0.85 | 0.07 | 0.98 |
| *ITGA4* | integrin subunit alpha 4 | 5.24 | 5.02 | 4.20 | 4.98 | 0.96 | 0.85 | 0.07 | 0.97 |
| *LAMB3* | laminin subunit beta 3 | 2.58 | 2.05 | 1.95 | 0.71 | 0.80 | 0.95 | 0.02 | 1.05 |
| *SELP* | selectin P | 1.73 | 2.15 | 2.09 | 1.53 | 1.25 | 0.96 | 0.02 | 1.05 |
| *COL6A1* | collagen type VI alpha 1 chain | 1.00 | 1.36 | 1.40 | 1.07 | 1.37 | 0.97 | 0.01 | 1.04 |
| *CTGF* | connective tissue growth factor | 3.36 | 2.87 | 2.75 | 3.25 | 0.85 | 0.97 | 0.01 | 1.02 |
| *ADAMTS8* | ADAM metallopeptidase with thrombospondin type 1 motif 8 | 3.51 | 2.39 | 2.43 | 3.44 | 0.68 | 0.99 | 0.00 | 1.02 |
| *LAMB1* | laminin subunit beta 1 | 2.91 | 3.37 | 3.40 | 2.90 | 1.16 | 0.99 | 0.00 | 1.01 |
| *PPIA* | peptidylprolyl isomerase A | 0.79 | 1.17 | 1.18 | 0.77 | 1.49 | 1.00 | 0.00 | 1.00 |

**Table S7: Summary of differentially regulated inflammation genes in the Veh and IL-1β (10ng/mL) treated hVCF using taqman gene array**

| **Gene** | **Gene name** | | **2^-ΔCt (Veh)** | | **2^-ΔCt (IL-1B)** | | **Standard deviation (Veh)** | | **Standard deviation (IL-1B)** | | **Fold Change (IL-1B/Veh)** | | **P value** | | **(-logP)** | | **FDR q Value** |
| --- | --- | --- | --- | --- | --- | --- | --- | --- | --- | --- | --- | --- | --- | --- | --- | --- | --- |
| *ITGAM* | integrin subunit alpha M | | 13.70 | | 98.40 | | 22.00 | | 65.46 | | 7.18 | | 0.10 | | 1.00 | | 6.35 |
| *PLCG2* | phospholipase C gamma 2 | | 1.07 | | 1.08 | | 0.47 | | 0.46 | | 1.01 | | 0.12 | | 0.91 | | 3.85 |
| *PTGS2* | prostaglandin-endoperoxide synthase 2 | | 0.98 | | 1.84 | | 0.01 | | 1.91 | | 1.87 | | 0.18 | | 0.75 | | 3.76 |
| *ANXA1* | annexin A1 | | 10.65 | | 24.32 | | 9.22 | | 13.96 | | 2.28 | | 0.23 | | 0.64 | | 3.63 |
| *BDKRB2* | bradykinin receptor B2 | | 1.07 | | 22.67 | | 0.46 | | 27.02 | | 21.13 | | 0.24 | | 0.62 | | 3.01 |
| *PLA2G4C* | phospholipase A2 group IVC | | 1.04 | | 1.25 | | 0.35 | | 1.08 | | 1.19 | | 0.26 | | 0.59 | | 2.69 |
| *PTGIS* | prostaglandin I2 synthase | | 1.33 | | 2.57 | | 1.08 | | 2.31 | | 1.93 | | 0.28 | | 0.55 | | 2.53 |
| *ANXA5* | annexin A5 | | 1.19 | | 0.59 | | 0.78 | | 0.39 | | 0.50 | | 0.31 | | 0.52 | | 2.41 |
| *NR3C1* | nuclear receptor subfamily 3 group C member 1 | | 1.02 | | 1.81 | | 0.45 | | 1.19 | | 1.77 | | 0.34 | | 0.46 | | 2.41 |
| *PLCG1* | phospholipase C gamma 1 | | 1.01 | | 1.69 | | 0.02 | | 0.60 | | 1.67 | | 0.35 | | 0.46 | | 2.20 |
| *PLCE1* | phospholipase C epsilon 1 | | 1.58 | | 0.76 | | 1.74 | | 0.62 | | 0.48 | | 0.36 | | 0.45 | | 2.05 |
| *VCAM1* | vascular cell adhesion molecule 1 | | 1.09 | | 2.44 | | 0.39 | | 2.51 | | 2.24 | | 0.36 | | 0.45 | | 1.88 |
| *ITGB1* | integrin subunit beta 1 | | 1.00 | | 1.34 | | 0.00 | | 0.58 | | 1.34 | | 0.36 | | 0.44 | | 1.76 |
| *BDKRB1* | bradykinin receptor B1 | | 1.18 | | 22.84 | | 0.78 | | 36.92 | | 19.32 | | 0.37 | | 0.43 | | 1.65 |
| *PLCB4* | phospholipase C beta 4 | | 1.42 | | 1.76 | | 1.02 | | 2.01 | | 1.24 | | 0.40 | | 0.40 | | 1.68 |
| *IL1R2* | interleukin 1 receptor type 2 | | 1.04 | | 1.68 | | 0.37 | | 0.75 | | 1.61 | | 0.41 | | 0.39 | | 1.61 |
| *PDE4B* | phosphodiesterase 4B | | 1.04 | | 1.94 | | 0.36 | | 2.69 | | 1.86 | | 0.41 | | 0.39 | | 1.53 |
| *CACNAB4* | calcium channels | | 1.42 | | 0.64 | | 1.47 | | 0.22 | | 0.45 | | 0.42 | | 0.38 | | 1.47 |
| *KLKB1* | kallikrein B1 | | 16.40 | | 9.42 | | 16.42 | | 6.16 | | 0.57 | | 0.43 | | 0.37 | | 1.42 |
| *CES1* | carboxylesterase 1 | | 2.11 | | 0.83 | | 2.53 | | 0.38 | | 0.39 | | 0.44 | | 0.36 | | 1.37 |
| *TNFRSF1B* | TNF receptor superfamily member 1B | | 2.11 | | 8.22 | | 2.49 | | 9.89 | | 3.89 | | 0.44 | | 0.36 | | 1.31 |
| *PTGS1* | prostaglandin-endoperoxide synthase 1 | | 1.51 | | 7.03 | | 1.56 | | 5.67 | | 4.67 | | 0.45 | | 0.35 | | 1.28 |
| *A2M* | alpha-2-macroglobulin | | 1.67 | | 3.34 | | 1.50 | | 3.18 | | 2.00 | | 0.46 | | 0.34 | | 1.25 |
| *LTB4R* | leukotriene B4 receptor | | 1.16 | | 3.18 | | 0.77 | | 4.21 | | 2.73 | | 0.46 | | 0.34 | | 1.21 |
| *TBXA2R* | thromboxane A2 receptor | | 1.04 | | 1.15 | | 0.37 | | 1.18 | | 1.11 | | 0.48 | | 0.32 | | 1.21 |
| *HRH1* | histamine receptor H1 | | 1.64 | | 2.47 | | 1.44 | | 1.43 | | 1.51 | | 0.52 | | 0.29 | | 1.25 |
| *MAPK14* | mitogen-activated protein kinase 14 | | 1.03 | | 1.45 | | 0.36 | | 0.97 | | 1.41 | | 0.52 | | 0.29 | | 1.21 |
| *PTGER2* | prostaglandin E receptor 2 | | 1.09 | | 1.51 | | 0.46 | | 1.54 | | 1.38 | | 0.52 | | 0.28 | | 1.18 |
| *PLA2G5* | phospholipase A2 group V | | 1.17 | | 1.18 | | 0.78 | | 0.79 | | 1.01 | | 0.53 | | 0.28 | | 1.15 |
| *LTA4H* | leukotriene A4 hydrolase | | 1.40 | | 1.41 | | 1.00 | | 1.01 | | 1.01 | | 0.53 | | 0.28 | | 1.11 |
| *IL1R1* | interleukin 1 receptor type 1 | | 3.08 | | 6.75 | | 4.38 | | 8.13 | | 2.19 | | 0.53 | | 0.28 | | 1.08 |
| *CASP1* | caspase 1 | | 2.04 | | 3.72 | | 1.92 | | 3.80 | | 1.83 | | 0.53 | | 0.27 | | 1.05 |
| *IL2RG* | interleukin 2 receptor subunit gamma | | 0.05 | | 0.03 | | 0.07 | | 0.02 | | 0.55 | | 0.57 | | 0.24 | | 1.09 |
| *PDE4D* | phosphodiesterase 4D | | 1.33 | | 2.08 | | 1.07 | | 2.53 | | 1.56 | | 0.60 | | 0.22 | | 1.11 |
| *ICAM1* | intercellular adhesion molecule 1 | | 13.71 | | 6.56 | | 22.68 | | 2.84 | | 0.48 | | 0.62 | | 0.21 | | 1.11 |
| *PTAFR* | platelet activating factor receptor | | 1.69 | | 2.16 | | 1.51 | | 0.93 | | 1.28 | | 0.62 | | 0.21 | | 1.09 |
| *PDE4A* | phosphodiesterase 4A | | 1.17 | | 5.84 | | 0.76 | | 8.82 | | 5.01 | | 0.64 | | 0.19 | | 1.09 |
| *CACNA1C* | calcium voltage-gated channel subunit alpha1 C | | 1.17 | | 1.51 | | 0.78 | | 0.88 | | 1.29 | | 0.64 | | 0.19 | | 1.07 |
| *NFKB1* | nuclear factor kappa B subunit 1 | | 1.19 | | 1.54 | | 0.79 | | 0.89 | | 1.29 | | 0.65 | | 0.19 | | 1.05 |
| *TNFSF13B* | tumor necrosis factor superfamily member 13b | | 1.69 | | 2.60 | | 1.55 | | 3.46 | | 1.54 | | 0.66 | | 0.18 | | 1.03 |
| *ITGB2* | integrin subunit beta 2 | | 1.76 | | 1.17 | | 2.00 | | 0.78 | | 0.67 | | 0.66 | | 0.18 | | 1.01 |
| *PLA2G1B* | phospholipase A2 group IB | | 0.04 | | 0.07 | | 0.05 | | 0.05 | | 1.65 | | 0.66 | | 0.18 | | 0.99 |
| *PTGDR* | prostaglandin D2 receptor | | 1.04 | | 2.25 | | 0.36 | | 2.98 | | 2.16 | | 0.67 | | 0.17 | | 0.98 |
| *CD40* | CD40 molecule | | 1.69 | | 2.16 | | 1.52 | | 0.92 | | 1.28 | | 0.67 | | 0.17 | | 0.96 |
| *IL1RAPL2* | interleukin 1 receptor accessory protein like 2 | | 1.00 | | 1.35 | | 0.01 | | 0.58 | | 1.35 | | 0.67 | | 0.17 | | 0.94 |
| *PTGFR* | prostaglandin F receptor | | 1.17 | | 1.00 | | 0.77 | | 0.01 | | 0.86 | | 0.68 | | 0.17 | | 0.93 |
| *CYSLTR1* | cysteinyl leukotriene receptor 1 | | 1.74 | | 1.31 | | 1.59 | | 0.43 | | 0.75 | | 0.70 | | 0.16 | | 0.94 |
| *PTGIR* | prostaglandin I2 (prostacyclin) receptor (IP) | | 1.33 | | 3.40 | | 1.07 | | 2.68 | | 2.56 | | 0.73 | | 0.14 | | 0.95 |
| *IL1RL1* | interleukin 1 receptor like 1 | | 1.19 | | 1.79 | | 0.77 | | 2.03 | | 1.51 | | 0.77 | | 0.11 | | 0.99 |
| *PLCD1* | phospholipase C delta 1 | | 1.08 | | 3.01 | | 0.47 | | 3.12 | | 2.79 | | 0.81 | | 0.09 | | 1.02 |
| *TNF* | tumor necrosis factor | | 1.07 | | 1.08 | | 0.48 | | 0.48 | | 1.01 | | 0.86 | | 0.07 | | 1.06 |
| *TBXAS1* | thromboxane A synthase 1 | | 1.68 | | 1.89 | | 1.50 | | 1.25 | | 1.13 | | 0.88 | | 0.05 | | 1.07 |
| *ADRB2* | adrenoceptor beta 2 | | 1.04 | | 1.14 | | 0.36 | | 1.17 | | 1.10 | | 0.89 | | 0.05 | | 1.06 |
| *MC2R* | melanocortin 2 receptor | | 1.17 | | 1.50 | | 0.78 | | 0.87 | | 1.28 | | 0.91 | | 0.04 | | 1.06 |
| *ANXA3* | annexin A3 | | 1.97 | | 1.78 | | 2.61 | | 1.27 | | 0.90 | | 0.91 | | 0.04 | | 1.05 |
| *CACNAB2* | calcium channels | | 1.03 | | 1.01 | | 0.46 | | 0.44 | | 0.98 | | 0.97 | | 0.02 | | 1.09 |
| *MAPK3* | mitogen-activated protein kinase 3 | | 1.09 | | 1.10 | | 0.48 | | 0.49 | | 1.00 | | 0.98 | | 0.01 | | 1.08 |
| *IL2RB* | interleukin 2 receptor subunit beta | | 1.26 | | 1.28 | | 1.07 | | 0.49 | | 1.01 | | 0.98 | | 0.01 | | 1.07 |
| *TNFRSF1A* | TNF receptor superfamily member 1A | | 4.81 | | 11.85 | | 6.95 | | 12.29 | | 2.46 | | 0.98 | | 0.01 | | 1.05 |
| *PLCB3* | phospholipase C beta 3 | | 2.79 | | 0.89 | | 3.40 | | 0.72 | | 0.32 | | 0.99 | | 0.01 | | 1.04 |
| *LTC4S* | leukotriene C4 synthase | | 1.77 | | 1.97 | | 1.65 | | 2.17 | | 1.11 | | 0.99 | | 0.00 | | 1.02 |
| *CACNA1D1* | calcium channels | | 1.01 | | 1.01 | | 0.01 | | 0.88 | | 1.01 | | 0.99 | | 0.00 | | 1.01 |
| *MAPK8* | mitogen-activated protein kinase 8 | | 1.78 | | 3.24 | | 2.02 | | 4.31 | | 1.82 | | 0.99 | | 0.00 | | 0.99 |

**Table S8:** Biological process gene ontology (GO:BP) and cellular component gene ontology (GO:CC), obtained through GProfiler, of secreted proteins expressed in conditional media of human ventricular cardiac fibroblasts treated with IL-1β (10ng/mL). Ontology terms represented with P value of 0.001 or less.

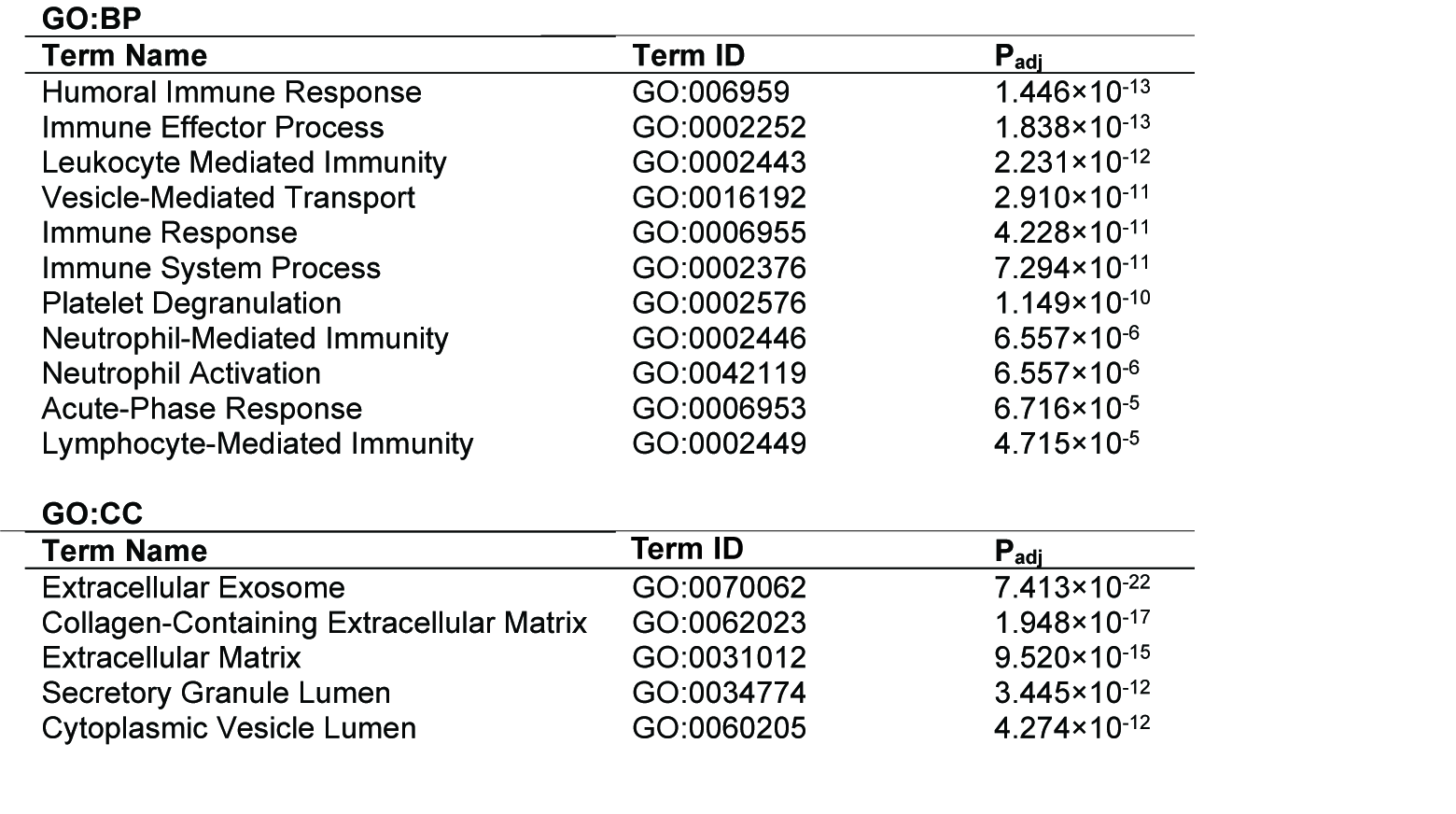

**Table S9:** Biological process gene ontology (GO:BP) level 3, immune subset identified through Database for Annotation, Visualization, and Integrated Discovery (DAVID) v6.8 of secreted proteins expressed uniquely in conditional media of human ventricular cardiac fibroblasts treated with IL-1β (10ng/mL). Multiplicity is controlled through Benjamini-Hochberg test.

| **Term** | **Gene Name** | **%** | **P Value** | **Fold Enrichment** | **Benjamini-Hochberg** |
| --- | --- | --- | --- | --- | --- |
| GO:0006952~defense response | complement factor H(CFH), amyloid P component, serum(APCS), attractin(ATRN), complement C1q C chain(C1QC), C-C motif chemokine ligand 2(CCL2), orosomucoid 1(ORM1), serpin family A member 3(SERPINA3), orosomucoid 2(ORM2), apolipoprotein A1(APOA1), complement C4A (Rodgers blood group)(C4A), apolipoprotein A2(APOA2), serpin family A member 1(SERPINA1), complement component 4 binding protein alpha(C4BPA), peptidoglycan recognition protein 2(PGLYRP2), complement C5(C5), complement C6(C6), CD5 molecule like(CD5L), histone cluster 2 H2B family member e(HIST2H2BE), interleukin 1 beta(IL1B), C-X-C motif chemokine ligand 5(CXCL5), C-X-C motif chemokine ligand 1(CXCL1), haptoglobin(HP), integrin subunit beta 1(ITGB1), kallikrein B1(KLKB1), low density lipoprotein receptor(LDLR), immunoglobulin heavy constant gamma 4 (G4m marker)(IGHG4), interleukin 6(IL6), kininogen 1(KNG1), inter-alpha-trypsin inhibitor heavy chain family member 4(ITIH4), complement factor B(CFB), intercellular adhesion molecule 1(ICAM1), joining chain of multimeric IgA and IgM(JCHAIN), serpin family F member 1(SERPINF1), C-X-C motif chemokine ligand 8(CXCL8), hemopexin(HPX) | 38.88889 | 6.12E-14 | 4.316775 | 3.76E-11 |
| GO:0006959~humoral immune response | immunoglobulin kappa variable 1D-33(IGKV1D-33), complement factor H(CFH), complement C1q C chain(C1QC), C-C motif chemokine ligand 2(CCL2), bone marrow stromal cell antigen 1(BST1), immunoglobulin kappa variable 4-1(IGKV4-1), immunoglobulin heavy constant gamma 4 (G4m marker)(IGHG4), complement C4A (Rodgers blood group)(C4A), immunoglobulin kappa variable 3-20(IGKV3-20), interleukin 6(IL6), complement component 4 binding protein alpha(C4BPA), complement factor B(CFB), complement C5(C5), joining chain of multimeric IgA and IgM(JCHAIN), complement C6(C6), histone cluster 2 H2B family member e(HIST2H2BE), hemopexin(HPX) | 18.88889 | 1.99E-13 | 12.83395 | 6.12E-11 |
| GO:0006956~complement activation | immunoglobulin kappa variable 1D-33(IGKV1D-33), complement C4A (Rodgers blood group)(C4A), immunoglobulin kappa variable 3-20(IGKV3-20), complement component 4 binding protein alpha(C4BPA), complement factor H(CFH), complement factor B(CFB), complement C5(C5), complement C6(C6), complement C1q C chain(C1QC), immunoglobulin kappa variable 4-1(IGKV4-1), immunoglobulin heavy constant gamma 4 (G4m marker)(IGHG4) | 12.22222 | 5.62E-10 | 17.60232 | 1.15E-07 |
| GO:0002684~positive regulation of immune system process | vascular endothelial growth factor C(VEGFC), immunoglobulin kappa variable 1D-33(IGKV1D-33), interleukin 1 beta(IL1B), complement factor H(CFH), C-X-C motif chemokine ligand 5(CXCL5), C-X-C motif chemokine ligand 1(CXCL1), complement C1q C chain(C1QC), bone marrow stromal cell antigen 1(BST1), C-C motif chemokine ligand 2(CCL2), immunoglobulin kappa variable 4-1(IGKV4-1), immunoglobulin heavy constant gamma 4 (G4m marker)(IGHG4), complement C4A (Rodgers blood group)(C4A), immunoglobulin kappa variable 3-20(IGKV3-20), interleukin 6(IL6), complement component 4 binding protein alpha(C4BPA), peptidoglycan recognition protein 2(PGLYRP2), complement factor B(CFB), intercellular adhesion molecule 1(ICAM1), complement C5(C5), C-X-C motif chemokine ligand 8(CXCL8), complement C6(C6), hemopexin(HPX) | 24.44444 | 1.87E-08 | 4.290566 | 2.3E-06 |
| GO:0002443~leukocyte mediated immunity | immunoglobulin kappa variable 1D-33(IGKV1D-33), interleukin 1 beta(IL1B), C-X-C motif chemokine ligand 5(CXCL5), complement C1q C chain(C1QC), immunoglobulin kappa variable 4-1(IGKV4-1), immunoglobulin heavy constant gamma 4 (G4m marker)(IGHG4), complement C4A (Rodgers blood group)(C4A), immunoglobulin kappa variable 3-20(IGKV3-20), interleukin 6(IL6), complement component 4 binding protein alpha(C4BPA), complement C5(C5), intercellular adhesion molecule 1(ICAM1), complement C6(C6), hemopexin(HPX) | 15.55556 | 4.4E-08 | 7.36277 | 4.51E-06 |
| GO:0002697~regulation of immune effector process | interleukin 1 beta(IL1B), complement factor H(CFH), C-C motif chemokine ligand 2(CCL2), apolipoprotein A1(APOA1), apolipoprotein A2(APOA2), complement C4A (Rodgers blood group)(C4A), interleukin 6(IL6), complement component 4 binding protein alpha(C4BPA), peptidoglycan recognition protein 2(PGLYRP2), complement factor B(CFB), complement C5(C5), complement C6(C6), hemopexin(HPX) | 14.44444 | 1.35E-07 | 7.488989 | 1.19E-05 |
| GO:0050776~regulation of immune response | immunoglobulin kappa variable 1D-33(IGKV1D-33), interleukin 1 beta(IL1B), alpha-1-microglobulin/bikunin precursor(AMBP), complement factor H(CFH), integrin subunit beta 1(ITGB1), complement C1q C chain(C1QC), immunoglobulin kappa variable 4-1(IGKV4-1), immunoglobulin heavy constant gamma 4 (G4m marker)(IGHG4), apolipoprotein A1(APOA1), complement C4A (Rodgers blood group)(C4A), immunoglobulin kappa variable 3-20(IGKV3-20), apolipoprotein A2(APOA2), interleukin 6(IL6), complement component 4 binding protein alpha(C4BPA), peptidoglycan recognition protein 2(PGLYRP2), complement factor B(CFB), intercellular adhesion molecule 1(ICAM1), complement C5(C5), complement C6(C6), hemopexin(HPX) | 22.22222 | 2.17E-07 | 4.105805 | 1.67E-05 |
| GO:0002250~adaptive immune response | immunoglobulin kappa variable 1D-33(IGKV1D-33), interleukin 1 beta(IL1B), complement C1q C chain(C1QC), immunoglobulin kappa variable 4-1(IGKV4-1), immunoglobulin heavy constant gamma 4 (G4m marker)(IGHG4), complement C4A (Rodgers blood group)(C4A), immunoglobulin kappa variable 3-20(IGKV3-20), interleukin 6(IL6), complement component 4 binding protein alpha(C4BPA), complement C5(C5), intercellular adhesion molecule 1(ICAM1), joining chain of multimeric IgA and IgM(JCHAIN), complement C6(C6), hemopexin(HPX) | 15.55556 | 2.98E-07 | 6.240824 | 2.03E-05 |
| GO:0050778~positive regulation of immune response | immunoglobulin kappa variable 1D-33(IGKV1D-33), interleukin 1 beta(IL1B), complement factor H(CFH), complement C1q C chain(C1QC), immunoglobulin kappa variable 4-1(IGKV4-1), immunoglobulin heavy constant gamma 4 (G4m marker)(IGHG4), complement C4A (Rodgers blood group)(C4A), immunoglobulin kappa variable 3-20(IGKV3-20), interleukin 6(IL6), complement component 4 binding protein alpha(C4BPA), peptidoglycan recognition protein 2(PGLYRP2), complement factor B(CFB), complement C5(C5), complement C6(C6), hemopexin(HPX) | 16.66667 | 1.3E-05 | 4.105805 | 0.000445 |
| GO:0002685~regulation of leukocyte migration | vascular endothelial growth factor C(VEGFC), interleukin 6(IL6), C-X-C motif chemokine ligand 5(CXCL5), intercellular adhesion molecule 1(ICAM1), C-X-C motif chemokine ligand 1(CXCL1), complement C5(C5), C-X-C motif chemokine ligand 8(CXCL8), C-C motif chemokine ligand 2(CCL2) | 8.888889 | 1.85E-05 | 9.601268 | 0.0006 |
| GO:0002687~positive regulation of leukocyte migration | vascular endothelial growth factor C(VEGFC), interleukin 6(IL6), C-X-C motif chemokine ligand 5(CXCL5), intercellular adhesion molecule 1(ICAM1), C-X-C motif chemokine ligand 1(CXCL1), C-X-C motif chemokine ligand 8(CXCL8), C-C motif chemokine ligand 2(CCL2) | 7.777778 | 2.39E-05 | 12.02361 | 0.000669 |
| GO:0002683~negative regulation of immune system process | apolipoprotein A2(APOA2), follistatin like 3(FSTL3), complement component 4 binding protein alpha(C4BPA), alpha-1-microglobulin/bikunin precursor(AMBP), peptidoglycan recognition protein 2(PGLYRP2), amyloid P component, serum(APCS), complement C5(C5), complement C1q C chain(C1QC), C-C motif chemokine ligand 2(CCL2), NME/NM23 nucleoside diphosphate kinase 1(NME1), apolipoprotein A1(APOA1) | 12.22222 | 3.74E-05 | 5.307917 | 0.000921 |
| GO:0030595~leukocyte chemotaxis | vascular endothelial growth factor C(VEGFC), interleukin 6(IL6), interleukin 1 beta(IL1B), C-X-C motif chemokine ligand 5(CXCL5), C-X-C motif chemokine ligand 1(CXCL1), complement C5(C5), C-X-C motif chemokine ligand 8(CXCL8), C-C motif chemokine ligand 2(CCL2) | 8.888889 | 7.45E-05 | 7.720607 | 0.001697 |
| GO:0045087~innate immune response | complement factor H(CFH), amyloid P component, serum(APCS), complement C1q C chain(C1QC), C-C motif chemokine ligand 2(CCL2), immunoglobulin heavy constant gamma 4 (G4m marker)(IGHG4), complement C4A (Rodgers blood group)(C4A), complement component 4 binding protein alpha(C4BPA), peptidoglycan recognition protein 2(PGLYRP2), complement factor B(CFB), complement C5(C5), intercellular adhesion molecule 1(ICAM1), joining chain of multimeric IgA and IgM(JCHAIN), complement C6(C6), histone cluster 2 H2B family member e(HIST2H2BE), hemopexin(HPX) | 16.66667 | 0.000116 | 3.359295 | 0.0023 |
| GO:0097529~myeloid leukocyte migration | vascular endothelial growth factor C(VEGFC), interleukin 6(IL6), interleukin 1 beta(IL1B), C-X-C motif chemokine ligand 1(CXCL1), complement C5(C5), C-X-C motif chemokine ligand 8(CXCL8), C-C motif chemokine ligand 2(CCL2) | 7.777778 | 0.000195 | 8.242598 | 0.003532 |
| GO:0050777~negative regulation of immune response | apolipoprotein A2(APOA2), complement component 4 binding protein alpha(C4BPA), peptidoglycan recognition protein 2(PGLYRP2), alpha-1-microglobulin/bikunin precursor(AMBP), apolipoprotein A1(APOA1) | 5.555556 | 0.00409 | 7.610761 | 0.041237 |
| GO:0002699~positive regulation of immune effector process | interleukin 6(IL6), interleukin 1 beta(IL1B), complement C6(C6), C-C motif chemokine ligand 2(CCL2), hemopexin(HPX) | 5.555556 | 0.011123 | 5.708071 | 0.079545 |
| GO:0002377~immunoglobulin production | interleukin 6(IL6), immunoglobulin kappa variable 4-1(IGKV4-1), polypeptide N-acetylgalactosaminyltransferase 2(GALNT2), hemopexin(HPX) | 4.444444 | 0.014289 | 7.80103 | 0.094489 |
| GO:0031349~positive regulation of defense response | interleukin 6(IL6), interleukin 1 beta(IL1B), peptidoglycan recognition protein 2(PGLYRP2), kallikrein B1(KLKB1), C-C motif chemokine ligand 2(CCL2), low density lipoprotein receptor(LDLR), hemopexin(HPX) | 7.777778 | 0.019619 | 3.260132 | 0.113338 |

**Table S10:** Biological process gene ontology (GO:BP) level 3, wounding subset identified through Database for Annotation, Visualization, and Integrated Discovery (DAVID) v6.8 of secreted proteins expressed uniquely in conditional media of human ventricular cardiac fibroblasts treated with IL-1β (10ng/mL). Multiplicity is controlled through Benjamini-Hochberg test.

| **Term** | **Gene Name** | **%** | **P Value** | **Fold Enrichment** | **Benjamini-Hochberg** |
| --- | --- | --- | --- | --- | --- |
| GO:0009611~response to wounding | tropomyosin 1 (alpha)(TPM1), hemoglobin subunit beta(HBB), pyruvate kinase, muscle(PKM), amyloid P component, serum(APCS), kallikrein B1(KLKB1), C-C motif chemokine ligand 2(CCL2), enolase 3(ENO3), annexin A2(ANXA2), apolipoprotein A1(APOA1), serpin family A member 1(SERPINA1), interleukin 6(IL6), kininogen 1(KNG1) | 13.33333 | 0.000603 | 3.461782 | 0.008429 |
| GO:1903035~negative regulation of response to wounding | kallikrein B1(KLKB1), kininogen 1(KNG1), amyloid P component, serum(APCS), annexin A2(ANXA2) | 4.44444 | 0.005569 | 11.01322 | 0.049641 |

**Table S11:** Biological process gene ontology (GO:BP) level 3, extracellular matrix subset identified through Database for Annotation, Visualization, and Integrated Discovery (DAVID) v6.8 of secreted proteins expressed uniquely in conditional media of human ventricular cardiac fibroblasts treated with IL-1β (10ng/mL). Multiplicity is controlled through Benjamini-Hochberg test.

| **Term** | **Gene Name** | **%** | **P Value** | **Fold Enrichment** | **Benjamini-Hochberg** |
| --- | --- | --- | --- | --- | --- |
| GO:0043062~extracellular structure organization | heat shock protein family A (Hsp70) member 8(HSPA8), ABI family member 3 binding protein (ABI3BP), versican (VCAN), intercellular adhesion molecule 1(ICAM1), integrin subunit beta 1(ITGB1), collagen type V alpha 2 chain (COL5A2), kallikrein B1(KLKB1), transthyretin(TTR), peroxidasin(PXDN), annexin A2(ANXA2), collagen type VIII alpha 1 chain(COL8A1) | 12.22222 | 1.06E-05 | 6.147677 | 0.000384 |

**Table S12:** List of metabolites that are differentially expressed in the conditioned media of primary human ventricular cardiac fibroblasts treated with IL-1β (10ng/mL)

| **Metabolite** | **Expression** | **HMDA Class** | **Role** |
| --- | --- | --- | --- |
| N-acetyl spermine | Higher | Polyamine | Cell growth and differentiation, androgenic effects of polyamines |
| Spermidine | Higher | Polyamine | Inhibits pro-inflammatory cytokine synthesis, Induction of autophagy and longevity |
| N6-Acetyl-L-lysine | Higher | Acetylated amino-acid | Biomarker of active genes, stroke risk |
| Acetyllysine | Lower | Acetylated amino-acid | Marker of exercise incapacity in patients with HF |
| N-acetyl-glucosamine-1-phosphate | Higher | n-acyl alpha hexosamines | Amino sugar metabolism |
| Glutathione | Higher | Antioxidant-waste disposal system | Glutathione induced immune stimulatory activity |
| Adenosine | Lower | Purine nucleoside | Endogenous regulator of innate immunity, coronary vasodilator |
| Nicotinamide Riboside | Higher | Nicotinate and nicotinamide metabolism | Nicotinamide decrease IL-8 production |
| D-sedoheptulose-1-7-phosphate | Lower | Hexose phosphates | type III secretion system dependant NFKB activation |
| p-aminobenzoate | Lower | organic compund | Essential nutrient for bacteria and called vitamin Bx |
| 2-oxo-4-methylthiobutanoate | Higher | Thia fatty acid | IL-3 dependant potent inducer of apoptosis |
| 1,3-diphosphateglycerate | Lower | 3-carbon organic molecule | Intermediate of glycolysis during respiration |
| Citrate-isocitrate | Lower | Tricarboxylic acids and derivatives | Citric acid cycle |
| Isocitrate | Higher | Tricarboxylic acids and derivatives | Citric acid cycle |
| Oxaloacetate | Lower | short chain keto acid | oxaloacetate can be converted to citric acid |
| Succinate | Lower | dicarboxylic acid | Complex II of the eletron transport chain |
| Lactate | Higher | Produced in the muscle | Energy source for the fibres of the heart, Lactate-pyruvate ratio is an indicator of oxidative stress |
| Citrate | Lower | Energy supply to the body |  |
| 2,3-dihydroxybenzoic acid | Lower | Human benzoic acid intermediate | intracellular iron deposition and tissue fibrosis |
| Cholic acid | Lower | primary bile acid | Facilitates fat absorption and cholesterol excretion |
| Pyroglutamic acid | Lower | Cyclized derivative of L-glutamic acid | Metabotoxin that causes adverse health effects |
| Atrolactic acid | Higher |  |  |

**Table S13:** Metabolite concentrations in the conditioned media of IL-1β treated primary human ventricular cardiac fibroblasts (P5). The p values represent two sided Student t test (P value threshold 0.05) and the FDR values calculated from parento scaling normalized data. The mean peak values representing the concentrations of metabolites were calculated from the raw data. FDR is the false discovery rate.

| **KEGG ID** | **Metabolite names** | **Mean peak area intensity +/- SD (Veh)** | **Mean peak area intensity +/-SD (IL-1β)** | **Standard deviation (Veh)** | **Standard deviation (IL-1β)** | **Fold change (IL-1β/Veh)** | **P value** | **(-logP)** | **FDR (q value)** |
| --- | --- | --- | --- | --- | --- | --- | --- | --- | --- |
| C00042 | Thiamine pyrophosphate |  | 4971.7 |  | 2891.1 |  |  |  |  |
| C00408 | dTMP |  | 4179.9 |  |  |  |  |  |  |
| C01026 | dTDP-nega |  | 4380.8 |  | 2680.7 |  |  |  |  |
| C00352 | dGMP |  | 4108.9 |  | 1163.4 |  |  |  |  |
| C00818 | D-glucosamine-1-phosphate |  | 7432.9 |  |  |  |  |  |  |
| C00352 | dGTP | 6598.3 |  | 2865.0 |  |  |  |  |  |
| C00364 | dUMP-nega |  | 4955.1 |  |  |  |  |  |  |
| C00160 | GTP-nega | 2065.4 |  | 583.6 |  |  |  |  |  |
| C03150 | O-acetyl-L-serine | 5754.5 |  | 1125.5 |  |  |  |  |  |
| C00378 | trans, trans-farnesyl diphosphate | 3717.6 |  | 583.4 |  |  |  |  |  |
| C00015 | UTP-nega | 4693.3 |  | 1919.1 |  |  |  |  |  |
| C00568 | Phenylpropiolic acid | 75714.3 | 2907872.1 | 11576.5 | 4908410.4 | 38.4 | 0.4 | 0.4 | 0.9 |
| C05382 | dTMP-nega | 6417.4 | 27713.6 | 4444.4 | 35939.7 | 4.3 | 0.4 | 0.4 | 0.9 |
| C00493 | spermine | 2891.8 | 11517.5 | 585.3 | 10537.9 | 4.0 | 0.4 | 0.4 | 0.9 |
| C00167 | Uric acid | 2489.1 | 8204.5 | 7.3 | 4627.5 | 3.3 | 0.2 | 0.7 | 0.9 |
| C00705 | deoxyadenosine | 2065.3 | 6608.9 | 583.5 | 2337.6 | 3.2 | 0.1 | 0.9 | 0.9 |
| C05463 | thymine | 6254.2 | 19984.2 | 128.8 | 19807.2 | 3.2 | 0.4 | 0.4 | 0.9 |
| C02504 | 2-Isopropylmalic acid | 72521.7 | 227037.0 | 4814.3 | 257779.9 | 3.1 | 0.4 | 0.4 | 0.9 |
| C00131 | dehydroascorbic acid | 7522.3 | 22906.3 | 6535.8 | 16814.3 | 3.0 | 0.2 | 0.7 | 0.9 |
| C03492 | 5-methoxytryptophan | 2891.0 | 8670.5 | 582.2 | 4086.7 | 3.0 | 0.2 | 0.7 | 0.9 |
| C00197 | 3-S-methylthiopropionate | 2477.6 | 7428.4 |  |  | 3.0 |  |  |  |
| C00465 | Pyrophosphate | 37180.3 | 88984.7 | 14867.3 | 109895.1 | 2.4 | 0.5 | 0.3 | 0.9 |
| HMDB02005 | N-acetyl spermidine | 1660.8 | 3643.9 |  | 2816.1 | 2.2 |  |  |  |
| HMDB0002108 | N-acetyl spermine | 11549.4 | 25276.4 | 2851.9 | 3304.5 | 2.2 | 0.0 | 2.3 | 0.5 |
| C01180 | 2-oxo-4-methylthiobutanoate | 12347.3 | 25439.1 | 5858.9 | 6388.9 | 2.1 | 0.1 | 1.2 | 0.9 |
| C00559 | deoxyinosine | 4126.9 | 8258.6 | 1165.6 | 5839.3 | 2.0 | 0.4 | 0.4 | 0.9 |
| C04483 | deoxyribose-phosphate | 3615.0 | 7186.8 | 748.0 | 2044.7 | 2.0 | 0.1 | 0.8 | 0.9 |
| HMDB00734 | Kynurenic acid | 2025905.7 | 3989798.0 | 1315608.2 | 3652608.6 | 2.0 | 0.4 | 0.4 | 0.9 |
| C00719 | biotin | 46310.7 | 89300.0 | 7237.8 | 55766.8 | 1.9 | 0.3 | 0.6 | 0.9 |
| C05852 | hypoxanthine | 4955.3 | 9451.3 | 1655.2 | 6602.7 | 1.9 | 0.3 | 0.5 | 0.9 |
| C00158 | coenzyme A-posi | 2477.5 | 4719.2 |  | 1214.7 | 1.9 |  |  |  |
|  | orotate | 71855.9 | 135862.4 | 10838.3 | 167204.0 | 1.9 | 0.5 | 0.3 | 0.9 |
| C00049 | Atrolactic acid | 37916.4 | 64371.3 | 13937.3 | 6295.8 | 1.7 | 0.0 | 1.4 | 0.9 |
| C00332 | Acetyllysine | 9619.4 | 16274.3 | 3739.2 | 12015.7 | 1.7 | 0.4 | 0.4 | 0.9 |
| C00061 | fructose-6-phosphate | 5367.8 | 9029.1 | 2925.5 | 2388.3 | 1.7 | 0.3 | 0.5 | 0.9 |
| C00026 | allantoin | 16184.1 | 26647.1 | 8184.3 | 10594.0 | 1.6 | 0.2 | 0.6 | 0.9 |
| C00356 | 3-hydroxybuterate | 13454.6 | 21576.4 | 7533.7 | 1480.3 | 1.6 | 0.1 | 0.9 | 0.9 |
| C00016 | folate | 15133.8 | 24166.1 | 2663.9 | 11972.4 | 1.6 | 0.3 | 0.6 | 0.9 |
| C00438 | Nicotinamide Riboside | 7432.0 | 11326.9 | 1649.8 | 7112.1 | 1.5 | 0.4 | 0.4 | 0.9 |
| C00882 | D-glucarate | 3302.7 | 4973.1 |  | 2382.0 | 1.5 |  |  |  |
| C00463 | isocitrate | 8744.2 | 12709.3 | 1181.3 | 2535.4 | 1.5 | 0.1 | 1.2 | 0.9 |
| C00065 | spermidine | 99278.5 | 144242.8 | 10511.3 | 19851.1 | 1.5 | 0.0 | 1.6 | 0.8 |
| C00198 | glucose-6-phosphate | 39261.6 | 56600.9 | 13324.8 | 17064.0 | 1.4 | 0.3 | 0.5 | 0.9 |
| C02918 | N-acetyl-glucosamine-1-phosphate | 15951.1 | 21545.5 | 14816.4 | 11516.4 | 1.4 | 0.6 | 0.2 | 1.0 |
| C00294 | Kynurenine | 7570.7 | 10146.8 | 2108.0 | 4575.4 | 1.3 | 0.5 | 0.3 | 0.9 |
| C00447 | sn-glycerol-3-phosphate | 111168.2 | 142903.5 | 42611.2 | 41244.6 | 1.3 | 0.4 | 0.4 | 0.9 |
| C00120 | carnitine | 128962.3 | 164318.6 | 13057.3 | 96747.9 | 1.3 | 0.6 | 0.2 | 0.9 |
| C00311 | lactate | 43867545.6 | 55529809.0 | 15572477.1 | 20366790.9 | 1.3 | 0.5 | 0.3 | 0.9 |
| C00078 | UMP | 3280.4 | 4130.3 |  |  | 1.3 |  |  |  |
| C03406 | malate | 335237.8 | 420964.6 | 70765.7 | 238599.8 | 1.3 | 0.6 | 0.2 | 0.9 |
| C01089 | 3-methylphenylacetic acid | 12651.8 | 15859.1 | 7557.7 | 10263.6 | 1.3 | 0.7 | 0.2 | 1.0 |
| C03722 | S-adenosyl-L-homoCysteine-posi | 4955.1 | 6193.9 |  | 1752.2 | 1.2 |  |  |  |
| C01236 | D-glyceraldehdye-3-phosphate | 4438.5 | 5466.9 | 1708.4 | 366.9 | 1.2 | 0.4 | 0.5 | 0.9 |
| C00295 | pantothenate | 668847.6 | 821713.7 | 132994.8 | 149481.3 | 1.2 | 0.3 | 0.6 | 0.9 |
| HMDB01107 | acetoacetate | 91474.3 | 111640.6 | 18913.6 | 31326.2 | 1.2 | 0.4 | 0.4 | 0.9 |
| C02494 | 1-Methyladenosine | 6096.6 | 7418.5 | 2533.9 | 2408.3 | 1.2 | 0.5 | 0.3 | 0.9 |
| C00361 | D-glucono-?-lactone-6-phosphate | 8315.9 | 10098.4 | 1516.8 | 4820.6 | 1.2 | 0.6 | 0.2 | 0.9 |
| C00079 | phosphocreatine | 319775.2 | 386385.4 | 32460.9 | 76082.4 | 1.2 | 0.2 | 0.6 | 0.9 |
| C00956 | anthranilate | 15968.5 | 19135.1 | 9801.9 | 2403.8 | 1.2 | 0.6 | 0.2 | 1.0 |
| C00860 | homocysteine | 11380.9 | 13628.1 | 582.6 | 4090.9 | 1.2 | 0.4 | 0.4 | 0.9 |
| C02571 | acetylphosphate | 41510.2 | 49631.3 | 41607.1 |  | 1.2 |  |  |  |
| C00570 | cholesterol | 59910.2 | 71345.5 | 11868.4 | 29167.1 | 1.2 | 0.6 | 0.2 | 0.9 |
| C00213 | S-methyl-5-thioadenosine | 2117.0 | 2512.9 | 560.2 |  | 1.2 |  |  |  |
| C00668 | glutathione | 19267.5 | 22805.6 | 2827.5 | 14928.9 | 1.2 | 0.7 | 0.2 | 1.0 |
| HMDB0002092 | L-arginino-succinate | 30334.9 | 35286.2 | 4816.1 | 6679.2 | 1.2 | 0.4 | 0.4 | 0.9 |
| C16463 | cysteine sulfinate | 5229.4 | 6063.6 | 515.5 | 5611.1 | 1.2 | 0.8 | 0.1 | 1.0 |
| HMDB00746 | hydroxyproline | 21438.5 | 24848.8 | 11121.6 | 16170.3 | 1.2 | 0.8 | 0.1 | 1.0 |
| C00118 | dihydroorotate | 115973.9 | 133286.9 | 46887.5 | 76175.6 | 1.1 | 0.8 | 0.1 | 1.0 |
| C00086 | xanthosine | 10449.5 | 11556.1 | 5489.5 | 7048.4 | 1.1 | 0.8 | 0.1 | 1.0 |
| C00170 | S-ribosyl-L-homocysteine-posi | 649413.2 | 712348.7 | 59301.2 | 98611.2 | 1.1 | 0.4 | 0.4 | 0.9 |
| C00882 | dGDP-nega | 4541.6 | 4955.2 | 1752.1 |  | 1.1 |  |  |  |
| C01144 | 3-phosphoglycerate | 11909.9 | 12993.6 | 3109.3 | 4219.7 | 1.1 | 0.7 | 0.1 | 1.0 |
| C00130 | inosine | 286305.8 | 309446.0 | 32473.8 | 71274.3 | 1.1 | 0.6 | 0.2 | 1.0 |
| C00864 | phenylpyruvate | 12626.4 | 13632.0 | 5203.5 | 9881.9 | 1.1 | 0.9 | 0.1 | 1.0 |
| HMDB00991 | 2-Aminooctanoic acid | 8918.7 | 9549.6 | 4146.6 | 2349.3 | 1.1 | 0.8 | 0.1 | 1.0 |
| C15608 | fructose-1,6-bisphosphate | 24438.8 | 26137.2 | 7521.5 | 8021.3 | 1.1 | 0.8 | 0.1 | 1.0 |
| C02727 | N-acetyl-glutamine | 64757.5 | 69011.1 | 73113.2 | 91643.7 | 1.1 | 1.0 | 0.0 | 1.0 |
| C00072 | aspartate | 2830883.4 | 2988371.4 | 698971.8 | 383149.0 | 1.1 | 0.7 | 0.1 | 1.0 |
| C00334 | 4-Pyridoxic acid | 24182.9 | 25527.6 | 16473.9 | 7577.5 | 1.1 | 0.9 | 0.0 | 1.0 |
| C00063 | cysteine | 112944.5 | 118985.8 | 33294.7 | 29892.2 | 1.1 | 0.8 | 0.1 | 1.0 |
| C00024 | aconitate | 112225.3 | 117654.5 | 21906.1 | 32475.1 | 1.0 | 0.8 | 0.1 | 1.0 |
| C00299 | Xanthurenic acid | 298163.1 | 312326.3 | 239786.8 | 254273.1 | 1.0 | 0.9 | 0.0 | 1.0 |
| C00109 | 2-oxobutanoate | 127614.1 | 133159.7 | 30704.5 | 56372.8 | 1.0 | 0.9 | 0.1 | 1.0 |
| C00144 | guanine | 10588.4 | 11001.8 | 2682.2 | 3654.7 | 1.0 | 0.9 | 0.1 | 1.0 |
| C00064 | glutathione disulfide-posi | 12032.2 | 12437.5 | 3846.4 | 3441.3 | 1.0 | 0.9 | 0.0 | 1.0 |
| C00135 | homocysteic acid | 26430.5 | 27261.9 | 15159.1 | 15282.9 | 1.0 | 0.9 | 0.0 | 1.0 |
| HMDB06029 | N-Acetylputrescine | 15708.9 | 16189.6 | 2679.5 | 3520.7 | 1.0 | 0.9 | 0.1 | 1.0 |
| C02835 | Indoleacrylic acid | 46067.9 | 47436.6 | 25937.6 | 17503.2 | 1.0 | 0.9 | 0.0 | 1.0 |
| C00458 | deoxyguanosine | 6883.0 | 7071.9 | 3726.6 | 2977.6 | 1.0 | 0.9 | 0.0 | 1.0 |
| C00105 | valine | 2762540.9 | 2834181.4 | 522316.3 | 567688.0 | 1.0 | 0.9 | 0.1 | 1.0 |
| C00750 | taurine | 29801.9 | 30541.5 | 4257.1 | 2544.0 | 1.0 | 0.8 | 0.1 | 1.0 |
| C01157 | Imidazoleacetic acid | 489139.2 | 499923.1 | 81656.9 | 59550.0 | 1.0 | 0.9 | 0.1 | 1.0 |
| C00123 | Maleic acid | 345604.0 | 353087.6 | 95495.8 | 28571.9 | 1.0 | 0.9 | 0.0 | 1.0 |
| C00029 | Urea | 153622.9 | 156769.9 | 18448.3 | 70600.8 | 1.0 | 0.9 | 0.0 | 1.0 |
| C04677 | arginine | 52828700.5 | 53877163.0 | 4037576.4 | 8569760.7 | 1.0 | 0.9 | 0.1 | 1.0 |
| C00575 | Cystine | 5377824.3 | 5478109.0 | 518556.6 | 2035423.5 | 1.0 | 0.9 | 0.0 | 1.0 |
| C00360 | dCTP-nega | 3302.8 | 3302.7 |  |  | 1.0 |  |  |  |
| C00024 | adenine | 404794.4 | 404666.2 | 16632.2 | 81734.9 | 1.0 | 1.0 | 0.0 | >0.999999 |
| C01384 | Methylcysteine | 404794.4 | 404666.2 | 16632.2 | 81734.9 | 1.0 | 1.0 | 0.0 | >0.999999 |
| C00345 | 7-methylguanosine | 13767.4 | 13757.2 | 3440.5 | 8020.5 | 1.0 | 1.0 | 0.0 | >0.999999 |
| C03539 | Taurodeoxycholic acid | 107092.1 | 105712.4 | 13376.5 | 32334.9 | 1.0 | 0.9 | 0.0 | 1.0 |
| C00455 | OBP | 19455.0 | 18993.1 | 2642.8 | 3295.3 | 1.0 | 0.9 | 0.1 | 1.0 |
| C00025 | glutathione disulfide-nega | 36723.2 | 35776.8 | 7699.9 | 8734.9 | 1.0 | 0.9 | 0.0 | 1.0 |
| C00438 | Nicotinamide ribotide | 5505.3 | 5354.8 | 954.9 | 2938.9 | 1.0 | 0.9 | 0.0 | 1.0 |
| PubChem Identifier (KEGG/HMDB) | myo-inositol | 15207718.8 | 14697583.0 | 2057383.8 | 1898898.8 | 1.0 | 0.8 | 0.1 | 1.0 |
| HMDB00563 | Phosphorylcholine | 64146.8 | 61875.2 | 14883.0 | 29437.9 | 1.0 | 0.9 | 0.0 | 1.0 |
| C00263 | hydroxyphenylpyruvate | 830811.3 | 799691.8 | 17272.9 | 52142.2 | 1.0 | 0.4 | 0.4 | 0.9 |
| C00127 | glycerate | 828746.1 | 796036.7 | 35304.8 | 172131.9 | 1.0 | 0.8 | 0.1 | 1.0 |
| C00002 | betaine | 1070121.4 | 1023654.5 | 218533.4 | 252452.4 | 1.0 | 0.8 | 0.1 | 1.0 |
| C00117 | sarcosine | 3900314.9 | 3726779.4 | 438769.7 | 1295396.2 | 1.0 | 0.8 | 0.1 | 1.0 |
| C01159 | 2,3-Diphosphoglyceric acid | 4308.2 | 4103.0 | 912.3 |  | 1.0 |  |  |  |
| C00341 | glucose-1-phosphate | 33921.8 | 32074.7 | 6060.0 | 7642.9 | 0.9 | 0.8 | 0.1 | 1.0 |
| C00149 | Methionine sulfoxide | 87560.1 | 82631.8 | 24939.1 | 17265.0 | 0.9 | 0.8 | 0.1 | 1.0 |
| C00166 | p-hydroxybenzoate | 97729.4 | 91880.7 | 37840.6 | 70355.6 | 0.9 | 0.9 | 0.0 | 1.0 |
| C00082 | uracil | 76914.5 | 71680.1 | 5059.4 | 1892.1 | 0.9 | 0.2 | 0.8 | 0.9 |
| HMDB00475 | betaine aldehyde | 48389.7 | 44940.7 | 17369.5 | 8040.1 | 0.9 | 0.8 | 0.1 | 1.0 |
| C00242 | hexose-phosphate | 60960.3 | 56127.7 | 5297.6 | 10027.7 | 0.9 | 0.5 | 0.3 | 0.9 |
| C00051 | glutathione-nega | 26298.8 | 24125.0 | 4309.5 | 13131.1 | 0.9 | 0.8 | 0.1 | 1.0 |
| C00498 | alanine | 5228301.0 | 4796127.5 | 383683.6 | 1577145.5 | 0.9 | 0.7 | 0.2 | 1.0 |
| C00021 | serine | 20054935.7 | 18395588.8 | 1144971.0 | 6324430.2 | 0.9 | 0.7 | 0.2 | 1.0 |
| C16511 | Hydroxyisocaproic acid | 510464.3 | 464367.5 | 55652.1 | 68877.3 | 0.9 | 0.4 | 0.4 | 0.9 |
| C00036 | Phenyllactic acid | 160473.1 | 145617.7 | 26090.4 | 33411.2 | 0.9 | 0.6 | 0.2 | 0.9 |
| C00417 | Adenylosuccinate | 115070.1 | 104333.9 | 42837.6 | 25925.3 | 0.9 | 0.7 | 0.1 | 1.0 |
| C00035 | glucosamine | 1235908.6 | 1116755.1 | 67073.3 | 174336.1 | 0.9 | 0.3 | 0.5 | 0.9 |
| C00670 | glyoxylate | 15015.9 | 13550.3 | 5797.3 | 4979.6 | 0.9 | 0.8 | 0.1 | 1.0 |
| C00629 | 2-dehydro-D-gluconate | 21688.6 | 19534.4 | 3719.2 | 6107.0 | 0.9 | 0.6 | 0.2 | 1.0 |
| C00005 | nicotinamide | 104498579.0 | 93850575.0 | 10852864.8 | 20082712.0 | 0.9 | 0.5 | 0.3 | 0.9 |
| C02630 | 2-hydroxygluterate | 503624.0 | 451181.1 | 55595.1 | 41761.9 | 0.9 | 0.3 | 0.6 | 0.9 |
| C00013 | ribose-phosphate | 11658.7 | 10416.3 | 6560.1 | 6475.3 | 0.9 | 0.8 | 0.1 | 1.0 |
| C00103 | glutamine | 82092538.1 | 73140380.2 | 2865570.5 | 9106056.4 | 0.9 | 0.2 | 0.7 | 0.9 |
| C00460 | Flavone | 20425.8 | 18121.8 | 6513.3 | 4029.5 | 0.9 | 0.6 | 0.2 | 1.0 |
| C00055 | Creatinine | 21146.8 | 18716.5 | 7090.5 | 7577.5 | 0.9 | 0.7 | 0.1 | 1.0 |
| C00047 | methionine | 1226252.7 | 1085196.8 | 151808.0 | 27338.7 | 0.9 | 0.2 | 0.7 | 0.9 |
| C00279 | D-gluconate | 131302.5 | 115926.5 | 14537.3 | 3919.9 | 0.9 | 0.2 | 0.8 | 0.9 |
| C00329 | glutamate | 9690338.2 | 8553138.1 | 1727038.7 | 1740264.3 | 0.9 | 0.5 | 0.3 | 0.9 |
| C00362 | dihydroxy-acetone-phosphate | 8701.8 | 7667.9 | 7055.1 |  | 0.9 |  |  |  |
| C00100 | Pyroglutamic acid | 958870.2 | 840319.6 | 37528.3 | 63907.1 | 0.9 | 0.1 | 1.3 | 0.9 |
| C00491 | cytosine | 337420.5 | 295633.4 | 45305.2 | 82136.2 | 0.9 | 0.5 | 0.3 | 0.9 |
| C00074 | proline | 31160052.1 | 26950570.5 | 4423268.0 | 4434999.7 | 0.9 | 0.3 | 0.5 | 0.9 |
| C00387 | histidine | 63147455.2 | 54366853.6 | 7322275.8 | 11274981.2 | 0.9 | 0.3 | 0.5 | 0.9 |
| C01152 | 1-Methyl-Histidine | 70150.3 | 60372.8 | 19207.4 | 38344.0 | 0.9 | 0.7 | 0.1 | 1.0 |
| C00127 | Glycerophosphocholine | 5686411.0 | 4885411.6 | 1696986.5 | 1828239.0 | 0.9 | 0.6 | 0.2 | 1.0 |
| C02170 | N-acetyl-glucosamine | 5782.7 | 4965.1 |  |  | 0.9 |  |  |  |
| C00141 | 2-keto-isovalerate | 366764.8 | 313697.6 | 88616.7 | 23274.7 | 0.9 | 0.4 | 0.4 | 0.9 |
| C01717 | leucine-isoleucine | 96782912.2 | 82724090.4 | 2396428.4 | 16193482.9 | 0.9 | 0.2 | 0.7 | 0.9 |
| C00155 | Hydroxyphenylacetic acid | 92066.5 | 78585.8 | 20634.8 | 63072.3 | 0.9 | 0.7 | 0.1 | 1.0 |
| C02612 | 2-Hydroxy-2-methylbutanedioic acid | 186310.0 | 158742.2 | 20949.0 | 78646.1 | 0.9 | 0.6 | 0.2 | 0.9 |
| C00005 | Ng,NG-dimethyl-L-arginine | 282483.8 | 239422.7 | 91068.2 | 159966.0 | 0.8 | 0.7 | 0.2 | 1.0 |
| C00224 | a-ketoglutarate | 5791637.4 | 4903165.0 | 452318.0 | 1020095.9 | 0.8 | 0.2 | 0.6 | 0.9 |
| C00214 | tyrosine | 1290795.3 | 1091576.8 | 182630.2 | 159115.4 | 0.8 | 0.2 | 0.6 | 0.9 |
| C00091 | threonine | 25234183.0 | 21309601.7 | 1044230.8 | 5610259.0 | 0.8 | 0.3 | 0.5 | 0.9 |
| C00777 | S-adenosyl-L-methioninamine | 8330.1 | 7027.9 |  | 2909.9 | 0.8 |  |  |  |
| HMDB01864 | 2-ketohaxanoic acid | 14211.5 | 11963.9 | 6102.7 | 5145.8 | 0.8 | 0.7 | 0.2 | 1.0 |
| C00062 | asparagine | 7246627.4 | 6086825.6 | 467921.7 | 1141953.8 | 0.8 | 0.2 | 0.7 | 0.9 |
| C00108 | Ascorbic acid | 2043146.7 | 1714385.7 | 503577.1 | 886389.7 | 0.8 | 0.6 | 0.2 | 1.0 |
| C00020 | arginosuccinic acid | 44621.7 | 37277.1 | 9257.8 | 4595.3 | 0.8 | 0.3 | 0.5 | 0.9 |
| C00257 | D-glucosamine-6-phosphate | 4955.3 | 4130.0 |  |  | 0.8 |  |  |  |
| C00111 | DL-Pipecolic acid | 8068006.5 | 6611906.7 | 895098.1 | 1866394.7 | 0.8 | 0.3 | 0.5 | 0.9 |
| C03626 | nicotinate | 304839.8 | 249526.4 | 76101.3 | 111833.5 | 0.8 | 0.5 | 0.3 | 0.9 |
| C00327 | creatine | 419903.1 | 339636.4 | 157894.8 | 222803.1 | 0.8 | 0.6 | 0.2 | 1.0 |
| HMDB02222 | 3-phospho-serine | 2068.5 | 1655.3 | 584.2 | 0.1 | 0.8 | 0.4 | 0.4 | 0.9 |
| C00112 | cholesteryl sulfate | 25738.0 | 20506.3 | 1672.3 | 19475.2 | 0.8 | 0.7 | 0.2 | 1.0 |
| C00048 | Guanidoacetic acid | 390047.7 | 304627.8 | 35528.3 | 87129.6 | 0.8 | 0.2 | 0.7 | 0.9 |
| C00156 | purine | 167279.1 | 130449.1 | 13371.5 | 44920.4 | 0.8 | 0.2 | 0.6 | 0.9 |
| C00041 | Aminoadipic acid | 529205.7 | 412010.3 | 86977.4 | 191935.6 | 0.8 | 0.4 | 0.4 | 0.9 |
| C08276 | 4-phosphopantothenate | 3768.7 | 2932.9 | 1775.2 | 1777.1 | 0.8 | 0.7 | 0.2 | 1.0 |
| C00979 | orotidine-5-phosphate | 12460.7 | 9642.7 | 9018.9 | 4305.5 | 0.8 | 0.7 | 0.1 | 1.0 |
| C05512 | dephospho-CoA-nega | 8670.6 | 6606.7 | 7591.2 | 2335.4 | 0.8 | 0.7 | 0.1 | 1.0 |
| C00044 | guanosine | 5379.9 | 4096.6 | 566.3 | 876.9 | 0.8 | 0.2 | 0.8 | 0.9 |
| C00186 | lysine | 4025561.1 | 3046125.2 | 495674.1 | 885946.0 | 0.8 | 0.2 | 0.8 | 0.9 |
| C00504 | fumarate | 419519.0 | 317257.2 | 96207.7 | 26821.7 | 0.8 | 0.2 | 0.8 | 0.9 |
| C00624 | N-acetyl-L-ornithine | 16523.5 | 12410.6 | 9718.9 | 3636.7 | 0.8 | 0.5 | 0.3 | 0.9 |
| C00188 | tryptophan | 9100800.7 | 6668861.2 | 1232518.8 | 2902576.3 | 0.7 | 0.3 | 0.6 | 0.9 |
| C01137 | shikimate | 10059390.7 | 7363365.9 | 6498214.5 | 7434861.0 | 0.7 | 0.7 | 0.2 | 1.0 |
| C06369 | 2-deoxyglucose-6-phosphate | 84255.2 | 61637.5 | 16652.7 | 20209.9 | 0.7 | 0.2 | 0.7 | 0.9 |
| C00100 | pyridoxine | 1141287.4 | 834836.3 | 133669.3 | 233605.1 | 0.7 | 0.1 | 0.9 | 0.9 |
| C01879 | riboflavin | 65769.9 | 47704.5 | 13442.2 | 9184.9 | 0.7 | 0.1 | 0.9 | 0.9 |
| C02226 | citrulline | 1421175.4 | 1024245.4 | 184904.1 | 885199.0 | 0.7 | 0.5 | 0.3 | 0.9 |
| C00253 | ornithine | 150337.9 | 108204.9 | 14488.1 | 65952.1 | 0.7 | 0.3 | 0.5 | 0.9 |
| C00106 | xanthine | 69089.5 | 49199.1 | 28962.9 | 31747.4 | 0.7 | 0.5 | 0.3 | 0.9 |
| C00356 | homoserine | 17014.4 | 12111.4 | 6323.8 | 3504.2 | 0.7 | 0.3 | 0.5 | 0.9 |
| C00083 | methylnicotinamide | 6287.9 | 4467.0 | 2038.5 | 280.6 | 0.7 | 0.3 | 0.5 | 0.9 |
| C00134 | pyruvate | 96006.4 | 67785.0 | 32212.7 | 26437.7 | 0.7 | 0.3 | 0.5 | 0.9 |
| C00043 | uridine | 12888.4 | 9087.1 | 8236.7 | 1273.8 | 0.7 | 0.5 | 0.3 | 0.9 |
| C00153 | O8P-O1P | 8808.9 | 6195.2 | 5977.9 | 2537.0 | 0.7 | 0.5 | 0.3 | 0.9 |
| C00526 | D-erythrose-4-phosphate | 33859.6 | 23249.9 | 15597.9 | 8949.4 | 0.7 | 0.4 | 0.4 | 0.9 |
| C00104 | indole | 28360.4 | 19391.9 | 7842.7 | 10130.9 | 0.7 | 0.3 | 0.5 | 0.9 |
| C01103 | phenylalanine | 13507914.9 | 9092541.0 | 1674663.1 | 4012913.3 | 0.7 | 0.2 | 0.8 | 0.9 |
| C00137 | N-acetyl-glutamate | 81018.0 | 53410.8 | 44732.2 | 85077.3 | 0.7 | 0.6 | 0.2 | 1.0 |
| C00254 | putrescine | 21416.2 | 13740.4 | 7802.8 | 9796.6 | 0.6 | 0.4 | 0.4 | 0.9 |
| C00187 | choline | 373867286.4 | 238201833.8 | 17644489.7 | 206309267.9 | 0.6 | 0.3 | 0.5 | 0.9 |
| C01589 | Indole-3-carboxylic acid | 338185.9 | 212961.0 | 114146.8 | 82865.3 | 0.6 | 0.2 | 0.7 | 0.9 |
| C00440 | 6-phospho-D-gluconate | 6603.2 | 4130.4 | 1176.8 | 3504.1 | 0.6 | 0.4 | 0.4 | 0.9 |
| C00337 | dimethylglycine | 60451143.1 | 37527151.1 | 1421161.9 | 32790243.7 | 0.6 | 0.3 | 0.5 | 0.9 |
| C00164 | Acetylcarnitine DL | 42678.9 | 26105.0 | 8913.3 | 14832.0 | 0.6 | 0.2 | 0.8 | 0.9 |
| C01479 | phosphoenolpyruvate | 6706.2 | 4074.0 |  |  | 0.6 |  |  |  |
| C00534 | quinolinate | 79790.6 | 48043.3 | 57876.0 | 60230.7 | 0.6 | 0.5 | 0.3 | 0.9 |
| C00576 | Carbamoyl phosphate | 151654.7 | 90784.6 | 102908.5 | 57305.9 | 0.6 | 0.4 | 0.4 | 0.9 |
| C00021 | SBP | 70103.8 | 41329.0 | 13652.9 | 18017.2 | 0.6 | 0.1 | 1.0 | 0.9 |
| C00083 | Methylmalonic acid | 320943.6 | 185908.2 | 100520.0 | 66474.3 | 0.6 | 0.1 | 0.9 | 0.9 |
| HMDB03320 | itaconic acid | 357129.6 | 206283.5 | 95949.1 | 136817.2 | 0.6 | 0.2 | 0.7 | 0.9 |
| C00185 | Cholic acid | 12321.9 | 7106.8 | 1497.0 | 3846.6 | 0.6 | 0.1 | 1.0 | 0.9 |
| C00300 | cyclic-AMP | 82874.5 | 47625.7 | 32434.8 | 16399.8 | 0.6 | 0.2 | 0.8 | 0.9 |
| C00122 | glucono-?-lactone | 7176.7 | 4074.2 | 847.0 |  | 0.6 |  |  |  |
| C00196 | 2,3-dihydroxybenzoic acid | 91493.3 | 51473.9 | 11087.4 | 27302.1 | 0.6 | 0.1 | 1.1 | 0.9 |
| C00093 | succinate | 299961.8 | 164899.1 | 25435.1 | 12191.3 | 0.5 | 0.0 | 2.9 | 0.2 |
| C00148 | Pyridoxamine | 4543.6 | 2425.5 | 1749.0 |  | 0.5 |  |  |  |
| C00236 | 1,3-diphopshateglycerate | 10876.4 | 5773.0 | 2439.4 | 1863.0 | 0.5 | 0.0 | 1.3 | 0.9 |
| C00695 | citrate | 7384821.6 | 3807418.2 | 957557.5 | 2165632.5 | 0.5 | 0.1 | 1.2 | 0.9 |
| C00140 | N-acetyl-L-aspartic acid | 4961.5 | 2524.0 |  |  | 0.5 |  |  |  |
| C12270 | N-Acetyl-L-alanine | 146626.2 | 73479.4 | 52193.6 | 47684.6 | 0.5 | 0.1 | 0.8 | 0.9 |
| C00008 | allantoate | 7383.3 | 3650.4 |  | 542.1 | 0.5 |  |  |  |
| C00258 | glycolate | 75951.1 | 37336.0 | 47589.1 | 28155.8 | 0.5 | 0.3 | 0.5 | 0.9 |
| C00073 | N6-Acetyl-L-lysine | 10563.8 | 5027.3 | 1376.4 | 1269.7 | 0.5 | 0.0 | 1.7 | 0.8 |
| HMDB00653 | Citraconic acid/itaconic acid | 326088.9 | 153454.6 | 44439.1 | 180435.3 | 0.5 | 0.2 | 0.7 | 0.9 |
| C00318 | CDP-nega | 8451.8 | 3770.1 | 4733.6 | 822.7 | 0.4 | 0.3 | 0.6 | 0.9 |
| C00022 | S-adenosyl-L-homocysteine-nega | 14866.3 | 6606.5 | 10150.3 | 1168.4 | 0.4 | 0.4 | 0.4 | 0.9 |
| C00791 | cystathionine | 15138.0 | 6607.9 | 5613.8 |  | 0.4 |  |  |  |
| C00077 | p-aminobenzoate | 23980.8 | 9477.5 | 12853.3 | 1842.8 | 0.4 | 0.1 | 0.9 | 0.9 |
| C02727 | adenosine | 9019.4 | 3256.9 | 4819.8 | 751.9 | 0.4 | 0.1 | 1.0 | 0.9 |
|  | oxaloacetate | 213838.6 | 72676.7 | 61963.8 | 10111.8 | 0.3 | 0.0 | 1.8 | 0.8 |
| C00114 | citrate-isocitrate | 16692631.3 | 5644964.4 | 920501.0 | 4359663.8 | 0.3 | 0.0 | 1.9 | 0.8 |
| HMDB00582 | D-sedoheptulose-1-7-phosphate | 100424.4 | 31726.2 | 40233.3 | 12853.3 | 0.3 | 0.0 | 1.3 | 0.9 |
| C02291 | cytidine | 42962.9 | 10842.4 | 21436.0 | 1281.7 | 0.3 | 0.1 | 1.2 | 0.9 |
